# Structural Basis for a Scaffolding Role of the COM Domain in Nonribosomal Peptide Synthetases

**DOI:** 10.1101/2025.06.08.658476

**Authors:** Julia Diecker, Benedikt Hermanns, Jennifer Rüschenbaum, René Rasche, Wolfgang Dörner, Alexander Schröder, Daniel Kümmel, Henning D. Mootz

## Abstract

Nonribosomal peptide synthetases (NRPSs) are multi-domain enzymes that catalyze the biosynthesis of therapeutically relevant natural products. Efficient peptide synthesis relies on intricate domain interactions, whose underlying principles remain poorly understood. The communication-mediating (COM) domains facilitate interactions between separate NRPS subunits like other docking domains, however, exhibit distinctive features that are unusual within this family: COM domains co-occur with epimerization (E) domains, are partially embedded within the adjacent condensation (C) domain and can also be found as an internal *cis*-COM domain with unknown function. We present the first crystal structure of a *cis*-COM domain within an E-COM-C domain arrangement from modules 4 and 5 of bacitracin synthetase 3 (BacC). The structure reveals a compactly folded COM domain sandwiched between E and C domains, suggesting a role in orienting these domains for efficient peptidyl carrier protein (PCP) shuttling. Through mutational analyses, dipeptide formation assays, and proximity-dependent photo-crosslinking experiments, we investigated both *cis*- and *trans*-COM domains and provide evidence supporting a principal role of COM domains as scaffolds of NRPS architecture. Their function as docking domains may be a secondary consequence of their division into separate donor and acceptor parts.

## Introduction

Non-ribosomal peptide synthetases are multi-functional protein templates for the biosynthesis of a plethora of peptide-based natural products (Fig. 1A).^[1-2]^ They activate amino acid building blocks under consumption of ATP, bind them as thioester on the 4’-phosphopantetheine (Ppant) moiety of the peptidyl-carrier protein (PCP; or thiolation domain; T). PCPs then pass on the aminoacyl and growing peptidyl intermediates to the catalytic sites for peptide bond formation, optional chemical modification like epimerization or *N*-methylation, and finally off-loading for product release. The corresponding adenylation (A), PCP, condensation (C), epimerization (E), *N*-methylation (M) and thioesterase (TE) domains are typically organized as a set of domains in a module that is responsible for the incorporation of one monomer. The number and order of modules determines the product sequence; an arrangement that has enabled the rational reprogramming of NRPSs in combinatorial approaches.^[3-4]^ A central mechanistic question is how the domain interplay is facilitated, in particular how the PCPs interacts with their various catalytic domain partners, as highlighted for the BacC4 module in Fig. 1C.^[5-9]^ Recent research in this regard has provided snapshots from crystal and cryo-EM structures^[10-18]^ as well as in-solution approaches that report on the conformational dynamics.^[19-24]^

**Figure 1.**
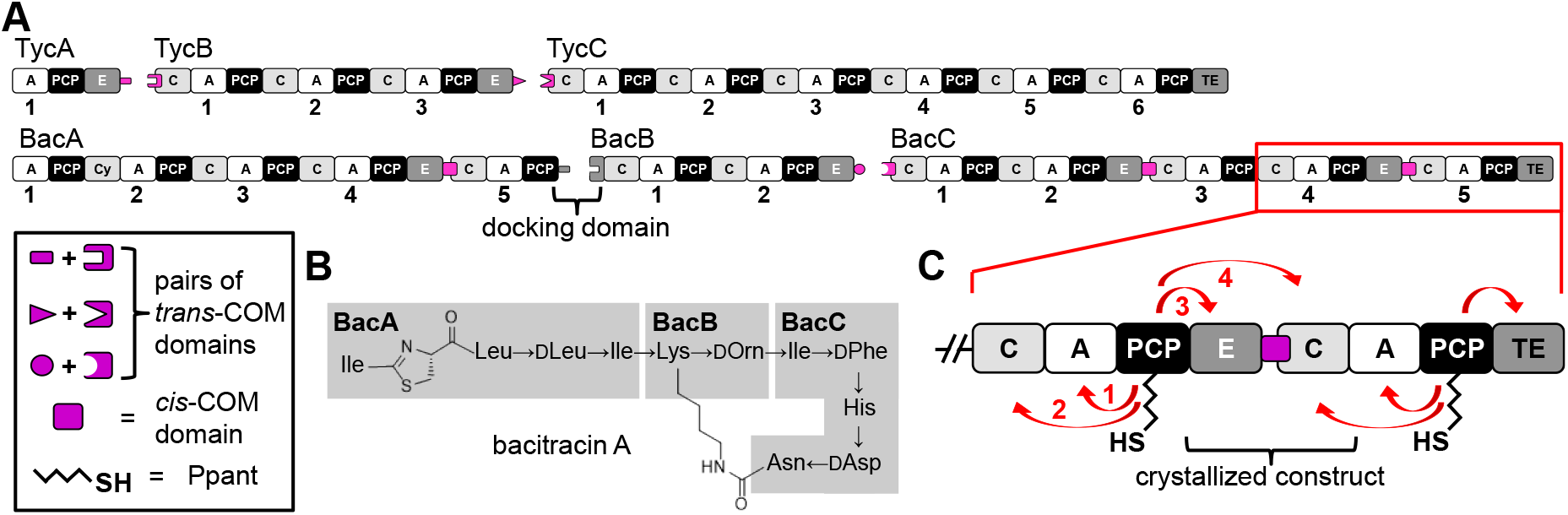
COM domains in NRPSs. (A) Shown are the tyrocidine A (top) and bacitracin A (bottom) NRPSs with *trans*- and *cis*-COM domains. (B) Structure of bacitracin A. (C) Interaction partners of the PCP of the BacC4-C5 modules are highlighted according to the biosynthetic logic. The BacC4 PCP shuttles between the domains of the crystallized E-COM-C construct.

Another important structural aspect pertains to intermolecular interaction of NRPS domains and modules that are located on separate polypeptide chains. Such arrangements are often observed for bacterial NRPSs. For example, the tyrocidine A and bacitracin A synthetases each consist of three enzymes (Figure 1A).^[25-26]^ Cognate pairs of docking domains at the respective C- and N-termini provide specific recognition between them. Several types of docking domains have been functionally and structurally characterized in NRPSs and the related polyketide synthases (PKSs),^[27-31]^ including the type found between the PCP of the BacA and C domain of the BacB synthetase (Figure 1A).^[32]^

However, the first identified type of docking domains, the so-called communication-mediating (COM) domains,^[33-34]^ are still insufficiently understood. The pair of donor COM (COM^D^) and acceptor COM domains (COM^A^) are the only known docking domains found at interfacing E and C domains, for example, between the tyrocidine TycA/TycB and TycB/TycC^[25]^ as well as the bacitracin BacB/BacC subunits (Fig. 1).^[26]^ Interestingly, there are also intramolecular *cis*-COM domains with high sequence homology,^[35]^ as for example in BacA and BacC (Fig. 1A), suggesting *trans*-COM domains might have arisen from *cis*-COM domains in gene splitting events. However, these *cis*-COM domains have received only little attention so far^[35-37]^ or were merely regarded as a linker between E and C domains.^[38]^

Initial biochemical analyses suggested *trans*-COM domains were only short interacting peptide sequences at the protein termini.^[33-34, 39]^ Later, a helix-hand model was proposed, inspired by an artificial protein-protein contact observed in the crystal lattice of the SrfA-C NRPS,^[10]^ in which an appended myc-tag appeared to mimic a C-terminal helix of the COM^D^ part. Notably, the model suggested that the COM^A^ domain also engages β-strand and turn elements from the internal, globular part of the C domain to form palm and fingers of the hand motif that embeds the COM^D^ helix. MD-modeling of a *trans*-COM domain supported the model based on the SrfA-C structure.^[36]^ Photo-crosslinking and MS mapping studies also corroborated a helix-hand interaction of an amphipathic helix with a hydrophobic palm and charged finger and hand motifs, but suggested a reversed orientation of the donor COM helix in an helix-up conformation.^[35]^ A structure of a native *trans*-COM complex has remained elusive^[36, 40-41]^ until very recently when the helix-hand proposal with the helix-up conformation was confirmed.^[17]^

Swapping and inserting native and artificial docking domains provides an enormous potential for engineering novel NRPSs with designer activity,^[42]^ however, this promise has not materialized so far for the COM docking domains. Despite some early successes in COM domain swapping experiments,^[33-34]^ subsequent studies yielded only modest and partially conflicting results.^[39, 43-45]^ We hypothesized that the presence of the *cis*-COM domains hinted at a more general role of COM domains, beyond intermolecular recognition mediated by *trans*-COM domains.

In this work, we report the first structure of a *cis*-COM domain in the context of its flanking E and C domains. The crystal structure revealed a compact COM domain forming extensive contacts with both catalytic domains and suggested the COM domain favorably orients the catalytic domains for efficient interaction with the common PCP partner. To probe this hypothesis, we biochemically characterized structural mutants of both *cis*- and *trans*-COM domains in peptide formation assays and analyzed conformational changes using a photo-crosslinking approach. Together, our data support the model of a novel scaffolding role of the COM domain as an essential structural part in the multi-domain organization of NRPS templates.

## Results and Discussion

### Crystal Structure of a Cis-COM Domain Sandwiched Between E and C Domains

We aimed to crystallize one of the underexplored *cis*-COM domains^[35]^ (see Fig. S1 for a sequence alignment of *trans*-COM and *cis*-COM domains). We selected the *cis*-COM domain between modules 4 and 5 of the BacC subunit including its upstream E and downstream C domains (termed E-COM-C; Fig. 1). We cloned the corresponding gene fragment encoding a 913 aa sequence following PCR-amplification from chromosomal DNA of *Bacillus licheniformis* ATCC10716 and expressed it in *E. coli*.^[26]^ Recombinant selenomethionine-substituted E-COM-C was purified by Ni-NTA and size-exclusion chromatography, crystalized and the structure determined by SAD phasing at ∼3.3 Å resolution. The structure could be confidentially modeled using computational models as template and the positions of the anomalous signals from selenomethionine as reference points (Tab. S1).

In the E-COM-C structure, the *cis*-COM domain appears as a small globular domain sandwiched between and partially overlapping with the E and C domains (Fig. 2A). Both the E and C domains adopt their canonical V-shaped structures.^[41, 46]^ Regions distant from the *cis*-COM domains are less well defined and show high b-factors, indicating flexibility of these parts. Also, several loop regions are not resolved in the electron density map. However, the *cis*-COM domain is well defined and makes extensive interactions with both E and C domains that result in the formation of rigid interfaces.

**Figure 2.**
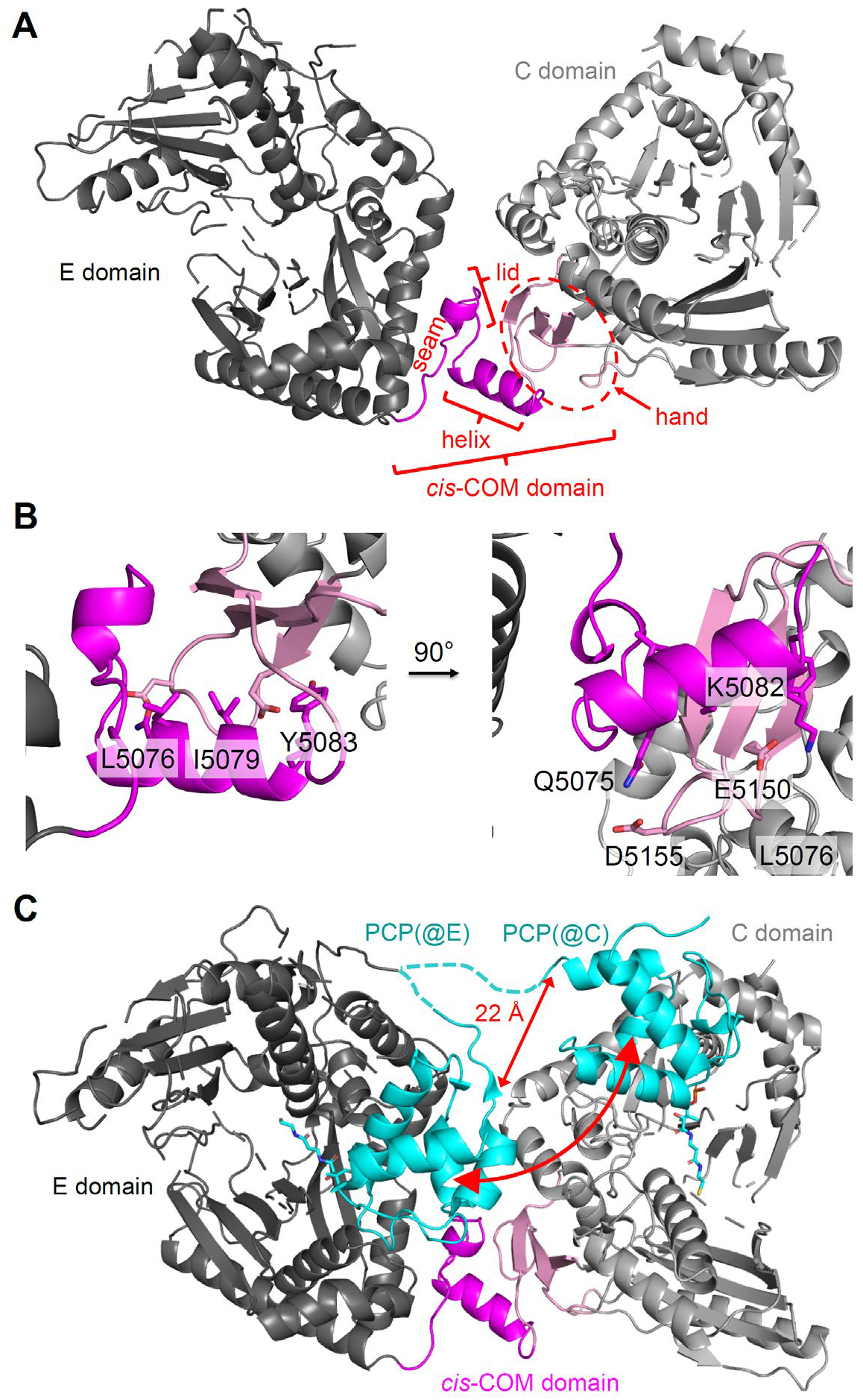
Crystal structure of the *cis*-COM domain. A) Entire structure of the E-COM-C ensemble of BacC4-C5. B) Zoom into the COM domain with some key residues highlighted. C) Modeling of the BacC4 PCP into the structure at the entry positions to the active sites of both the E domain and the donor site of the C domain, using pdb files 5isx and 6mfz, respectively. The red arrows indicate the PCP movement between the domains and the distance between the PCP’s C-terminal ends in each conformation.

The *cis*-COM domain shows the helix-hand arrangement with a helix up conformation, in agreement with the revised upside-down helix-hand model.^[35]^ The helix packs with three hydrophobic residues (L5076, I5079 and Y5083), which are conserved across COM domains,^[35]^ against the palm of the hand motif, e.g. the mostly hydrophobic central residues of three β-strands (I5091, V5160, S5148; Fig. 2B). This part of the structure is consistent with previous work on *trans*-COM domains.^[10, 33, 35-36]^ The back side of the central helix contains more hydrophilic and charged residues, some of which interact with counterparts on the side of the palm and the finger motifs (e.g., Q5075/D5155, K5082/E5150 & E5152; Fig. 2B). Importantly, the structure also reveals the folding of previously uncharacterized parts of the COM sequence that pertain to the stretch upstream of the central helix. Here, a seam segment (aa5056-5061) follows the last helix of the E domain as a mostly extended polypeptide chain, which then turns to fold into a lid region (aa5062-5072) that closes off the COM domain structure between the N-terminal end of the central helix and the first β-strand of the palm region. Together, seam, lid and helix represent what was initially defined as the COM^D^ domain in *trans*-COM domains,^[33]^ and all these segments make extensive contacts with the E domain. The lid residue D5068 interacts with R5158. Y5066 packs against residues L5076, I5091 and I5088, for example, making up a hydrophobic core with participation from the lid, helix palm and thumb. Finally, a loop element (L5200-E5203) in the C domain caps off the finger, palm and C-terminal end of the helix (Fig. 2B). In the contiguous polypeptide chain of the *cis*-COM domain, the C-terminal end of the helix (part of COM^D^) and the N-terminal end of the thumb (part of COM^A^) are connected by a peptide bond, similar to a helix-thumb-palm arrangement in a PCP-C didomain of the fungal TgaA NRPS,^[47]^ which, however, has no further homology to COM domains. Overall, the COM domain appears to act rather as a rigid connector than as a flexible hinge between its neighboring E and C domains.

The E-COM-C crystal structure can be superimposed well with the recently reported *trans*-COM cryo-EM structure^[17]^ (r.s.m.d. = 3.5 Å over 632 C-α of the entire structure and 1.3 Å over 42 C-α of the COM domain). Both structures show the same relative orientations of the neighboring E and C domains (Fig. S2), supporting a common structural role,^[35]^ with only slight differences in the binding modes of the central helix (Fig. S3).

### The COM Domain Favorably Positions E and C Domains for Interaction with BacC4 PCP

In the E-COM-C structure, the E and C domains are oriented such that the PCP binding sites on both domains face each other (donor position on the C domain). Such an orientation appears functionally relevant because in the NRPS synthesis logic the BacC4 PCP has to visit both domains consecutively and probably dynamically.^[22, 48-49]^ While the PCP is tethered via a linker to the E domain, ensuring the proper interaction, it has to reach the donor position of the C domain through the interdomain space resulting from the orientation of the two domains. To examine the possible path of the PCP when shuttling between the two catalytic domains, we modeled it into both binding sites using overlays with a PCP-E structure showing the epimerization conformation and a PCP-C structure showing the donor condensation conformation.^[16, 41]^ The model in Figure 2C shows that the PCP’s C-terminal residue as a pivot point has to translocate by about 22 Å and additionally the PCP body has to undergo rotation towards the other catalytic site by about 180°. While representing a significant conformational change, it is important to note that this PCP movement is a necessity to direct the Ppant group to two different domain entry sites. In fact, the path of the PCP appears to be nearly perfectly *minimized* by the observed positioning of E and C domains relative to one another, thereby minimizing entropic penalties and facilitating efficient PCP visits. Together, these considerations provide a rationale for why the COM domain acts as a rigid connector between the E and C domains.

We thus hypothesized a role for the COM domain to properly position the C domain relative to the PCP-E unit, likely as a rigid connector. A different COM structure resulting in a wider opening of the cleft between E and C domains or in a larger twist relative would necessitate additional movements of the catalytic domain bodies.

### A functional dipeptide formation assay reveals sensitivity of the *cis*-COM domain towards sequence changes

To test our hypothesis of a structural role of the COM domain we aimed for a mutagenesis study as outlined in Figure 3A and B. To this end, we developed a functional assay involving the crystalized *cis*-COM domain (Fig. 1C and 3C). To shortcut the bacitracin biosynthesis leading up to undecapeptidyl intermediate we aimed to convert the BacC4 module into an initiation module through deletion of its C domain and upstream modules.^[21, 49]^ Assuming sufficient substrate tolerances of the catalytic domains with the altered intermediates, the dimodular BacC4-C5 protein (domain composition A-PCP-E-COM-C-A-PCP-TE; construct **1**; 2362 aa) should activate and covalently bind L-Asp, racemize it to the D-aspartyl thioester and process this moiety along to the BacC5 module, where the resulting D-Asp-L-Asn dipeptide should be cleaved off by the TE domain (Fig. 3C).

**Figure 3.**
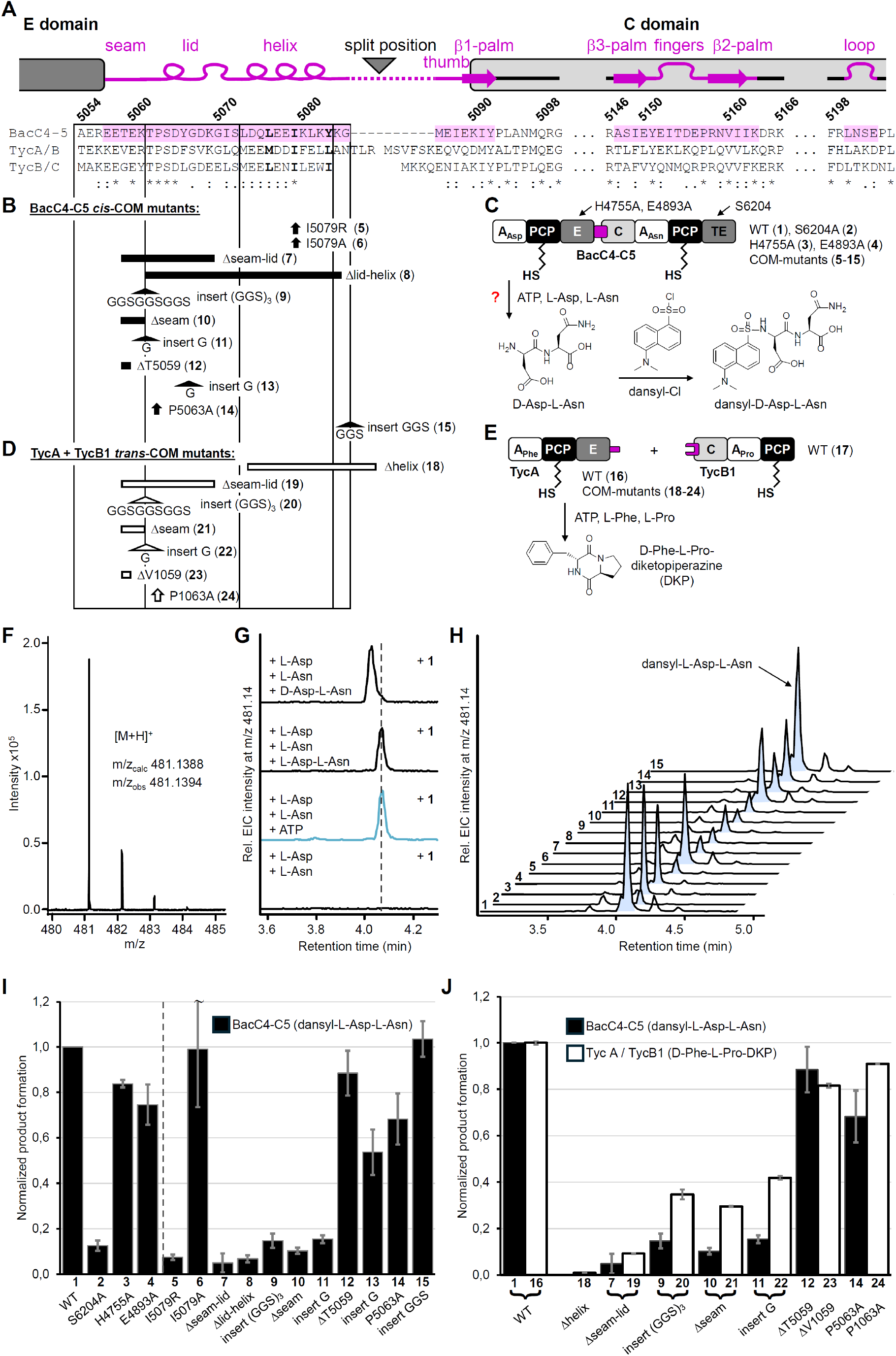
Mutational analysis of *cis*- and *trans*-COM domains. A) Structural features and sequence of the BacC4-C5 *cis*-COM domain with sequences of the *trans*-COM sequences of the TycA/B and TycB/C pairs. B) Schematic representation of investigated BacC4-C5 as well as its catalytic domain and *cis*-COM mutants. C) Scheme of the product formation assay with BacC4-C5. D) Schematic representation of investigated TycA/TycB1 system and its COM_D_ mutants. E) Scheme of the product formation assay with TycA and TycB1. F) Mass spectrum of the dansylated product of BacC4-C5 (**1**) fitting dansyl-Asp-Asn. G) EIC spectra of BacC4-C5 (**1**) assays including the indicated substrates and chemical standards. H) EIC spectra of product assays with all BacC4-C5 mutants. I) Diagram of product levels as analyzed in H, normalized to the wildtype (WT) level. J) Product formation of equivalent mutants of the *cis*-COM and *trans*-COM systems, each normalized to the wildtype (WT) level. Error bars represent standard deviations calculated from mean values.

We prepared recombinant holo-BacC4-C5 (**1**) by protein expression in *E. coli* and treatment of the purified protein with Ppant-transferase Sfp and CoA (Fig. S4).^[50]^ We confirmed its specific covalent and ATP-dependent binding of the substrate amino acids by ESI-MS (Fig. S4B). To assay for dipeptide formation, we added ATP (5 mM), L-Asp (2 mM) and L-Asn (2 mM). To increase the column retention time of the expected D-Asp-L-Asn product during LC-MS analysis we used dansyl chloride to convert all potential peptide products into the dansylated form (Fig. 3C). Indeed, we could detect the expected product peptide mass, and its formation was dependent on the holo-form of the protein as well as on the presence of all substrates, as expected (Fig. 3F and G). Dipeptide formation remained linear over time for about 1 h (Fig. S5) at a rate of about 1 molecule per 3 min (Fig. S6). A control construct in which the catalytic serine residue in the TE domain was mutated (S6204A; construct **2**) showed only 13% dipeptide production compared to the WT construct **1** (Fig. 3H and I; see Fig. S4A for all purified Bac4-C5 constructs), indicating that dipeptide formation occurred along the thiotemplate mechanism, as expected.

However, additional control experiments with two mutant proteins with an inactivated E domain (by mutation of the key catalytic residues H4755A and E4893A in constructs **3** and **4**, respectively) showed only a moderate reduction in product yields to about 80% (Fig. 3H and I), suggesting that the participation of the E domain was not crucial. Indeed, comparison with synthetic dipeptide standards showed that the BacC4-C5 constructs had formed the diastereomer L-Asp-L-Asn instead of D-Asp-L-Asn (Fig. 3G). Based on precedent in the literature, a likely explanation for this observation is that the E domain exhibits a strong substrate preference for its native undecapeptidyl substrate and was not capable to convert the L-aminoacyl moiety into the D-stereoisomer.^[49]^ In contrast, the BacC5 C domain appeared to have a high tolerance towards the opposite substrate stereochemistry at the donor site^[51-53]^ as it accepted the L-Aspartyl substrate for elongation to the L-Asp-L-Asn dipeptidyl moiety.

We reasoned that, despite these limitations, the dimodular BacC4-C5 model protein should still be valuable to study our hypothesis. The question whether the COM domain has a structural role to present the C domain in an optimized distance and orientation relative to the PCP-E unit should be independent of which aminoacyl thioester epimer is presented to the BacC5 C domain and hence independent of the catalytic participation of the E domain.

We next probed the functional outcome of structural perturbations of the COM domain by insertions, deletions and point mutations of varying severity (Fig. 3B). Our rationale was that structural changes were likely to be tolerated if the *cis*-COM domain mostly serves as a linker to ensure covalent connectivity. If, on the other hand, the *cis*-COM domain is critical in the fine-tuned positioning of the C domain, dipeptide formation should be sensitive against structural perturbations.

A destabilizing mutation I5079R in the central and hydrophobic helix-hand interaction interface (construct **5**) resulted in a drastic product decrease down to 7% compared to the unmutated construct **1**. In contrast, the mutation I5079A (**6**) had no significant effect, likely because it still supported the hydrophobic packing of the helix in the hand (Fig. 3B, H and I). In construct **7**, we left the helix-hand region untouched to ascertain its folding but deleted a part of the seam and most of the lid, to perturb the positioning of E and C domains. This structural change led to a massive drop in product formation to 5% (Fig. 3B, H and I). These findings clearly showed that the COM domain is more than a simple covalent linker.

We then introduced carefully designed alterations in the COM sequence that had the potential to still support the overall positioning of the C domain relative to the E domain. We reasoned that by deleting the lid and the helix in construct **8**, a similar distance between the E and C domains could be maintained, when assuming that the seam region can act as an extended bridge towards G5085 at the N-terminal end of the thumb region. Similarly, inserting a (GGS)_3_ linker before the lid (**9**) should increase the possible distance of E and C domains but still allow the native positioning when the flexible linker is oriented away from the COM (Fig. 3B). Both mutants **8** and **9** showed dipeptide product yields of only 7 to 15%, suggesting the changes were too drastic for a proper E and C domain positioning Fig. 3H and I. Even mutants with more subtle alterations to the amino acid sequence were heavily impaired. Deletion of three amino acids T5059 to K5061 (**10**) within the seam abrogated dipeptide formation to ≤15%, just like the insertion of a single glycine residue (**11**), indicating a high overall sensitivity of the E-COM-C interface towards structural perturbations (Fig. 3B, H and I).

We also observed a less pronounced but still significant impact with much more subtle pertubations: Deletion of a single residue in the seam (ΔT5059; **12**) or insertion of a single glycine residue between lid and helix (**13**), resulted in 88% and 54% product yield, respectively. Furthermore, mutation of the reasonably well conserved proline in the lid only reduced yields to 68% (P5063A; **14**). Only the insertion of the flexible linker sequence GGS at the end of the helix (**15**) showed no discernible loss of product yield (Fig. 3B, H and I). This position corresponds to the split site into *trans*-COM domains according to multiple sequence alignments (Fig. 3A and S1). The observed tolerance is therefore not surprising and further supports the idea that *trans*-COM domains can evolve from *cis*-COM domains. Together, the mutational analysis supported our model of an important structural role of the COM domain.

### Dipeptide formation and proximity-dependent chemical crosslinking in *trans*-COM assay confirms scaffolding role of COM fold

We next asked whether the structural role of *cis*-COM domains extends to *trans*-COM domains. We chose the first two modules from the tyrocidine biosynthesis (Fig. 1A), namely L-Phe-activating TycA (module composition A-PCP-E-COM^D^) and the L-Pro activating TycB1 (COM^A^-C-A-PCP) module taken from a truncated TycB enzyme (constructs **16** and **17**, Fig. 3A and E).^[25]^ The D-Phe-L-Pro dipeptide formed as thioester on the TycB1 autocatalytically cleaves itself off the template by cyclization to the D-Phe-L-Pro-diketopiperazine (DKP), thereby leading to product turnover and enabling sensitive readout of the enzymatic activity.^[35, 54]^ A negative control construct of TycA with the COM^D^ helix deleted (**18**) showed completely abrogated product formation, consistent with previous reports (Fig. 3D, H and I and Fig. S7).^[33,35, 55]^

We introduced most of the analogous deletions, insertions and mutations into seam and lid regions of the TycA COM^D^ domain as described above for the BacC4-5 *cis*-COM domain (Fig. 3D). The mutant enzymes were unaffected in their aminoacylation activity (Fig. S8) and were then assayed for dipeptide formation with TycB1 (Fig. 3E and Fig. S7). Notably, the structural changes overall had very similar dramatic impact on the *trans*-COM system. Specifically, the deletion of a seam-lid part (ΔV1059 to K1069; **19**), the insertion of a flexible (GGS)_3_ linker (between R1061 and T1062; **20**) or of a single glycine (**22**) between seam and lid, as well as the deletion of the last three residues of the seam (V1059 to R1061) (**21**) led to significant reductions in the D-Phe-L-Pro-DKP production down to 9, 35, 42, and 30 %, respectively (Fig.3D, H and I). Better tolerated were only the deletion of the single V1059 in the seam region (**23**) as well as the P1063A mutation in the lid (**24**), again consistent with the relative trends observed for the *cis*-COM domain of BacC4-C5 (Fig. 3D, H and I). Collectively, these findings corroborated the idea of a high structural similarity between *cis*- and *trans*-COM domains and of a common scaffolding role.

To collect direct evidence for the hypothesis that the COM domain positions the C domain relative to the PCP-E unit, we turned to conformationally sensitive chemical crosslinking. Proximity-dependent photo-activated crosslinking is useful to probe the interaction between two molecules or proteins.^[56-57]^ We previously reported the incorporation of the genetically encoded photo-crosslinking amino acid Bpa^[56, 58-59]^ into the GrsA/TycB1 and TycA/TycB1 pairs to investigate proximity parameters in the *trans*-COM domains.^[35, 55]^ Importantly, Bpa incorporated at Ser5 in the COM^A^ thumb region of TycB1 (TycB1(S5Bpa)) (**25**) results in two crosslink types into different regions of TycA (**16**), one reflecting a direct COM interface interaction, the other a conformation-dependent domain encounter (Fig. 4A and Fig. S9).^[55]^ The protein band above the 200 kDa marker on an SDS-PAGE gel corresponds to a crosslink to TycA residue(s) within the COM^D^ helix (calculated weight of a TycA-TycB1 crosslink is 246808 kDa; Fig 4A&B and Fig. S9 and S10). This crosslink, termed L-crosslink, results from the helix-thumb contact of the COM^D^/COM^A^ interaction, which is confirmed by our new E-COM-C structure. The second crosslink, termed T-crosslink, migrates at the upper end of the SDS gel (6% polyacrylamide; apparent molecular weight of 300-400 kDa; Fig. 4A&B)^[55]^ and connects Ser5Bpa of the thumb with a region of the functionally unrelated small A domain subunit (A^C^) of TycA (Fig. S10). This crosslink captures a close proximity brought about by the overall 3-D multi-domain arrangement (Fig. S9B).^[55]^ The aberrantly slow migration in the SDS-PAGE gel stems from the branched connectivity of the two polypeptide chains in T-shape (Fig. 4A).^[55]^ Here, we reasoned that the conformationally dependent T-crosslink should also report on the COM-mediated positioning of the C domain relative to the PCP-E unit, because the A^C^ domain is directly linked to the PCP, while the thumb region of the COM domain with Ser5Bpa sits on the body of the C domain (Fig. S9). A perturbed COM structure should thus affect the proximity between A^C^ and the thumb region. Notably, our crystal structure is consistent with this possible structural arrangement, because the residue corresponding to Ser5 in the COM^A^ of TycB1 faces the domain interspace accessible to the PCP and A^C^ domains.

**Figure 4.**
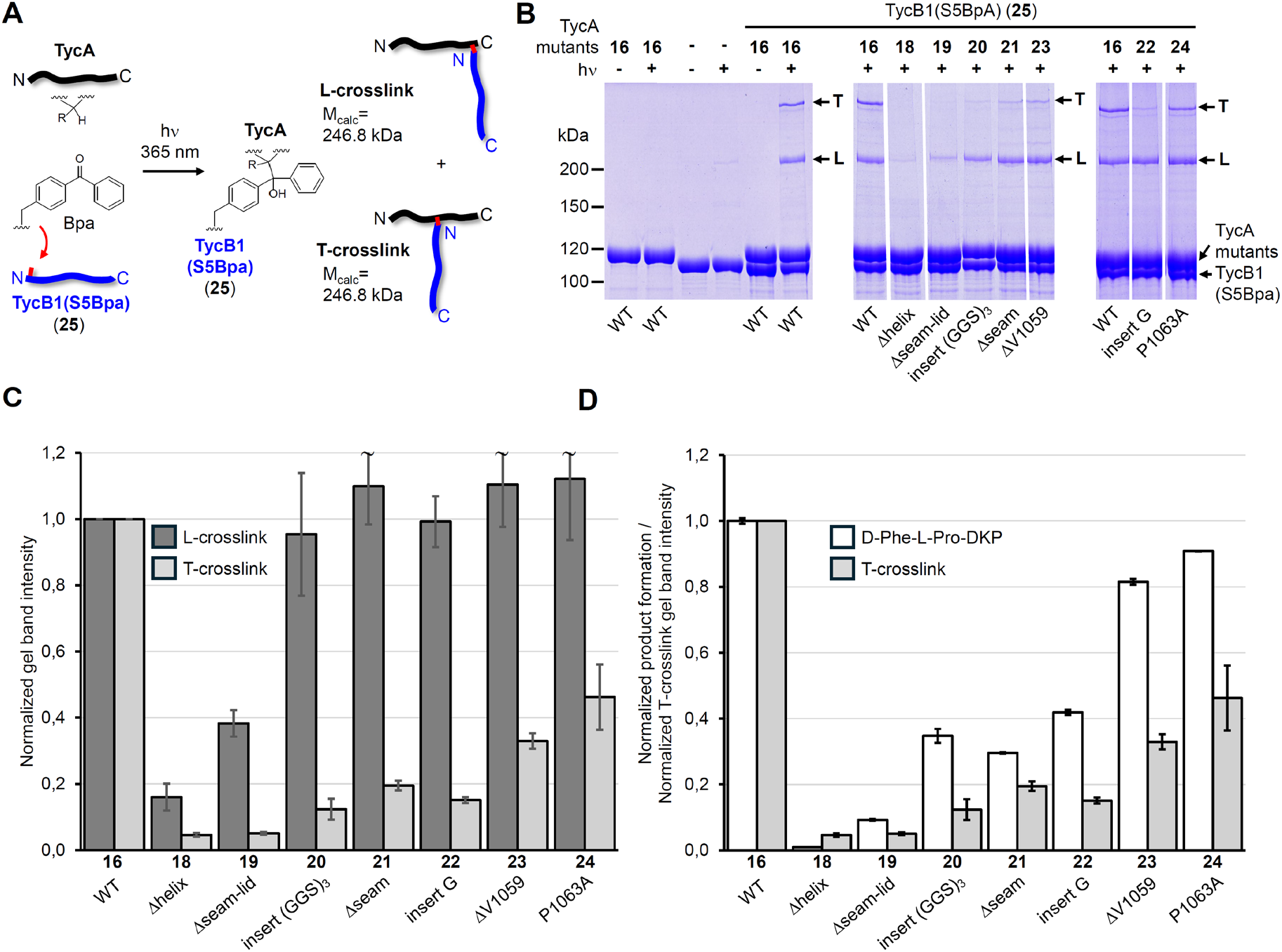
Photo-activated crosslinking analysis. The indicated COM_D_ of TycA were combined with TycB1(S5Bpa). A) Scheme of the crosslinking reaction resulting in nearly linear (L-form) and branched (T-form) polypeptide backbone connectivities of identical mass but different SDS-PAGE migration behavior. B) Analysis of photo-crosslink products on coomassie brilliant blue stained SDS-PAGE gels. Shown are lanes from 3 different gels, which were analyzed in relation to the wildtype TycA control (with **16**) present on each gel. C) Diagram of L- and T-crosslink intensities from experiments shown in B) analyzed by densitometry, normalized to the wildtype TycA control. D) Comparison of T-crosslink intensities shown in C) with D-Phe-L-Pro-DKP yields as shown in Figure 3J, each normalized to the wildtype (WT) levels.

The negative control mutant of TycA with the COM^D^ helix deleted (**18**) showed a nearly complete loss of both L- and T-crosslinks with TycB1(S5Bpa), consistent with a lack of association between the two proteins (Fig. 4B and C). The seam and lid deletion mutant **19** showed a significant reduction in the L-crosslink (down to 38% compared to WT TycA), together with a virtually complete disappearance of the T-crosslink (5%) (Fig. 4B and C). Since the diminished L-crosslink is a clear indication of an impaired intermolecular association with TycB1 it is possible that the observed reduction of this mutant in the dipeptide product formation assay to 9% is also a result of the reduced affinity towards TycB1, prohibiting further structural conclusions. In contrast, analysis of the remaining internal TycA mutants in the COM^D^ region (**20, 21, 22, 23** and **24**) showed that they all formed nearly unchanged L-crosslink levels compared to TycA WT (Fig. 4B and C). These observations suggested that the structural alterations in their COM^D^ domains did not strongly affect the association with the COM^A^ domain of TycB1, likely because their helix and lid regions were left basically intact. However, the formation of their T-crosslinks was strongly reduced (Fig. 4B and C). Interestingly, remaining levels of these T-crosslinks showed a correlation with the levels of dipeptide formation observed for the same mutants. In the first group of mutants (**20, 21** and **22**) T-crosslink levels dropped to 12-19% relative to WT TycA (Fig. 4B and C) and these proteins exhibited only 30-42% of dipeptide formation with TycB1 (Fig. 4D). On the other hand, a second group of mutants (**23** and **24**) retained relatively higher T-crosslink yields of 33% and 46% (Fig. 4B and C), which correlated with significantly higher dipeptide formation levels of 82 and 91%, respectively (Fig. 4D). Thus, these observations established a link between the domain arrangement in 3-D space and the dipeptide formation activity. Since the latter requires PCP interaction with the downstream C domain across the COM interface, these data provide direct evidence to support our model of the COM domain’s structural role for the conformational alignment in the multi-domain ensemble. Perturbing the structure of the COM domain has the potential to misposition the connected C domain relative to the PCP-E unit, resulting in impaired PCP interaction with the C domain and reduced peptide elongation activity (Fig. 5).

**Figure 5.**
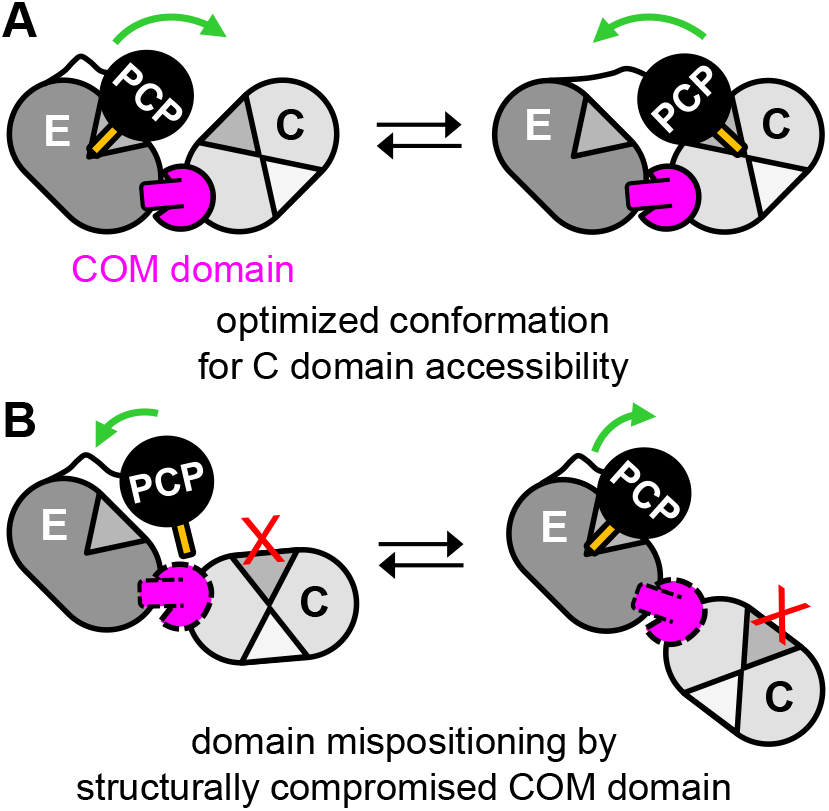
Model for structural role of the COM domain. A) Proper positioning of the C domain relative to the PCP-E unit. B) Consequences of a structurally altered COM domain.

## Conclusions

Multi-functional NRPSs and other related multi-domain enzymes like polyketide and fatty acid synthases^[60]^ have to coordinate the interaction of the central carrier protein with various catalytic domains to allow for a stepwise and directional product assembly.^[5-8]^ Recent work on NRPS conformations in solution suggest they can adopt a dynamic mixture of interconverting conformations with the PCP shuttling between catalytic domains in finely graded fashion.^[19-23]^ The A domain is one driving force for conformational changes with its ATP-driven cycle of amino acid adenylation and thiolation reactions.^[61]^ Furthermore, the chemical state of the Ppant business end (e.g., free thiol group, aminoacyl or peptidyl thioester) influences the relative population of different possible conformations as it determines for which catalytic domain(s) it serves as the fitting substrate or acts by product inhibition.^[19]^ Preferred adoption of a particular conformation helps to prime the enzyme for a productive PCP encounter with a catalytic domain, which together with the catalytic properties of the latter will decide on the rate of catalyzing the next step.

We here show that the COM domain, both as *cis*-COM and as split *trans*-COM domain, plays an important scaffolding role in addition to its previously described function as a *trans*-docking domain. The resulting domain positioning appears as another factor affecting the proper conformations of an NRPS assembly line to optimize product throughput. The importance of this aspect of COM domains is reflected by the observation that, to the best of our knowledge, a *cis*-COM or *trans*-COM domain is always found when an E domain is integrated into the NRPS assembly line (note that another type of epimerization domain exists that is inserted differently).^[62]^ In contrast, other docking domain types are only found as *trans*-forms in split NRPS chains, where they reside as additional tag sequences at the C- and N-termini. We propose that all COM domains properly position the C domain relative to the preceding PCP-E unit and have corroborated this model by a mutational analysis and conformationally dependent crosslinking. The COM-mediated domain positioning facilitates a shortened and entropically optimized trajectory of the PCP between the two domains. This arrangement allows the PCP to efficiently visit the donor site of the C domain and probably is important for a balanced population of all necessary conformations along a catalytic cycle, according to the PCP transitions to all its domain partners (Fig. 5 and Fig. S9B). PCP trajectories are likely also optimized when shuttling between other catalytic domains. For example, the trajectories between C (acceptor position) and A domains^[12]^ as well as between formylation (F) and A domains^[11]^ are structurally well investigated. The distances the PCP must travel in these two latter cases are much larger than the distance between the E and C domains as investigated here. The associated conformational changes also rely on differently stretched conformations of the involved A domain with its two subdomains, which are also ATP-driven.^[61]^ Notably, the *cis*-COM and *trans*-COM^[17]^ domain mediated positioning of E and C domains provides the first structurally investigated multi-domain ensemble to deal with PCP shuttling that does not involve the mobile A domain.

Extrapolating from the model of the COM domain’s scaffolding role, we further propose that several other structural factors in NRPS assembly lines will have important effects on the relative population of different conformations, for example, domain size and shape as well as their positioning mediated through contact interfaces and linkers^[63]^ of different length and flexibility. The impact of these parameters on the domain interplay has been hardly studied so far.

Finally, our *cis*-COM structure, together with the recently reported *trans*-COM structure,^[17]^ provide the long-awaited structural blueprints to identify specificity-determining molecular interactions in split *trans*-COM domains that could be exploited in improved COM-swap engineering experiments. However, the extended interaction interface, which includes substantial contact areas with the bodies of the E and C domains, suggests a limited matchmaker potential of such COM-swaps compared to other docking domain architectures.

Intriguingly, the high sensitivity of the NRPS templates investigated in this study towards structural perturbations in the *cis*-or *trans*-COM domains suggests that collateral sequence changes resulting from COM-swap experiments might cause impairment in productivity. Further research may help rationalize the success of COM-based combinatorial engineering.

## Materials and Methods

### DNA cloning of expression plasmids

The gene fragment encoding BacC4-C5[E-COM-C] was PCR-amplified from chromosomal DNA of *Bacillus licheniformis* ATCC 10716 (DSM 603; DSMZ-German Collection of Microorganisms and Cell Cultures GmbH) using oligonucleotides 5′-GTTACCATGGACTCAGATGAAGAAATGAAC-3′ and 5′-CATGGATCCTATTTTGACAGTCTTATTGATG-3′. The fragment was digested using *Nco* I and *Bam* HI and ligated into pET28a to give pAF24.

The gene fragment encoding BacC4-C5[A-PCP-E-COM-C-A-PCP-TE] was PCR-amplified from the same chromosomal DNA using oligonucleotides 5′-GCTGACCATGGGTGATAAATCAGCATCATATG-3′ and 5′-GCTCTGGATCCGGAAAGGACTGTGATATCG-3′ and ligated into pET28a using restriction sites *Nco* I and *Bam* HI to give pJD01. Derivatives of pJD01 coding for mutations, deletions and insertions in the COM domain were introduced using site-directed mutagenesis. All resulting amino acid sequences are given below.

pJR89 encoding SBP-TycA[A-PCP-E-COM^D^] was created from pJR88 (Ref^[55]^) by introducing a stop-codon following the gene encoding TycA to remove the following His-tag from the open reading frame and create the native C-terminal end in the encoded SBP-TycA. Derivatives of pJR89 to encode for mutations, deletions and insertion in the COM domain were introduced using site-directed mutagenesis. All resulting amino acid sequences are given below.

pBH96 was created by introducing an amber stop-codon into pJR95 (Ref^[21]^) using site-directed mutagenesis at the position coding for S5 in TycB1[COM^A^-C-A-PCP].

See Table S2 for all plasmids encoding the recombinant proteins.

### Protein production

Proteins were produced in *E. coli* LOBSTR BL21(DE3) cells carrying the encoding plasmid and the plasmid pLacIRARE2 that decodes rare codons. LB medium containing the antibiotics ampicillin at 100 µg/mL or kanamycin at 50 µg/mL and chloramphenicol at 34 µg/mL was inoculated 1:100 from an overnight culture. The culture was incubated at 37°C until the optical density of OD_600_ ∼0.8 was reached. Gene expression was induced by adding IPTG to a final concentration of 400 µM and conducted at 18°C for 16 hours and 180 rpm.

Cells expressing the genes encoding BacC4-C5-His_6_ wild type or mutant proteins thereof were harvested by centrifugation and resuspended in 10-15 mL ice-cold HEPES A buffer (50 mM HEPES, 100 mM NaCl; pH 8.0) for the purification via Ni^2+^-NTA affinity chromatography.

Cells expressing the genes encoding SBP-TycA wild type or mutant proteins thereof were harvested by centrifugation and resuspended in 15 mL ice-cold Strep-Tactin buffer (100 mM Tris/HCl, 150 mM NaCl; 1 mM EDTA, pH 8.0) for the purification via Strep-Tactin affinity chromatography.

Cells expressing the gene encoding TycB1-His_6_ were harvested by centrifugation and resuspended in 15 mL ice-cold HEPES A buffer (50 mM HEPES, 150 mM NaCl, pH 8.0) for the purification via Ni^2+^-NTA affinity chromatography.

TycB1(S5Bpa) containing p-benzoylphenylalanine (Bpa) was produced in LOBSTR BL21(DE3) cells using nonsense suppression. The cells carried the plasmid encoding TycB1(S5*tag*)-His_6_ as well as the plasmid pEVOL-Bpa encoding the tRNA-synthetase /tRNA_CUA_ pair.^[56]^ LB medium containing the antibiotics ampicillin at 100 µg/mL and chloramphenicol at 34 µg/mL was inoculated 1:100 from an overnight culture. The culture was incubated at 37°C until the optical density of OD_600_ ∼0.7 was reached. Before gene expression was induced by the addition of IPTG at a final concentration of 400 µM as well as 0.2 % (w/v) L-arabinose, Bpa was added at a final concentration of 1 mM (dissolved in 1 M NaOH). Gene expression was conducted at 28°C for 4 hours and 180 rpm.

BacC4-C5[E-COM-C]-His_6_ was produced in *E. coli* LOBSTR BL21(DE3) cells carrying the encoding plasmid. LB medium containing kanamycin at 50 µg/mL was inoculated 1:100 from an overnight culture and incubated at 37°C and 180 rpm until the optical density of OD_600_ ∼0.7 was reached. Gene expression was induced by the addition of IPTG at a final concentration of 400 µM and conducted at 28°C for 4 hours and 180 rpm.

The gene encoding BacC4-C5[E-COM-C]-His_6_ was also expressed as selenomethionine derivative. The protein was produced in LOBSTR BL21(DE3) cells carrying the encoding plasmid. M9 medium (Na_2_HPO_4_ 6,8 g/L, KH_2_PO_4_ 3 g/L, NaCl 0.59 g/L, NH_4_Cl 1 g/L, glucose 2 g/L, MgSO_4_ 0.241 g/L, CaCl_2_ 0.011 g/L, thiamine 0.5 mg/L) containing all necessary amino acids, selenomethionine and the corresponding antibiotic kanamycin at 50 µg/mL was inoculated 2:100 with harvested cells from an overnight culture. The culture was incubated at 37°C until the optical density of OD_600_ ∼0.8 was reached. Gene expression was induced by the addition of IPTG to a final concentration of 400 µM and conducted at 16°C for 16 hours and 180 rpm.

### Cell lysis and protein purification

To purifiy proteins via their C-terminal His_6_-tags, the cells were disrupted by a homogenizer (EmulsiFlex-C5, Avestin). The suspension was centrifugated 30 min at 25,000 x g and 4°C to separate soluble proteins from insoluble material and cell debris. The soluble protein fraction was adjusted to 20 mM imidazole. A Ni^2+^-NTA gravity flow column was equilibrated with HEPES A containing 20 mM imidazole (HEPES A-20) and the soluble fraction was loaded to the column twice. The column was then washed with 15 column volumes (cv) of HEPES A-20 and 10 cv of HEPES A-35 containing 35 mM imidazole. Proteins were eluted with HEPES B containing 250 mM imidazole. In the case of selenomethionine, all buffers were degassed and DTT was added at a final concentration of 1 mM to prevent oxidation.

The pooled fractions (selected by concentration and purify check using SDS-PAGE and Coomassie staining) were dialyzed first against assay buffer (50 mM HEPES, 100 mM NaCl, 10 mM MgCl_2_, 1 mM EDTA, pH 7.0) + 2 mM DTT, followed by buffer without DTT, and finally against buffer with 10 % glycerol. Proteins were concentrated when concentrations were below 25 µM using spin columns with a molecular cut-off of 30 kDa (TycA and TycB1 constructs) or 100 kDa (BacC4-C5 constructs) (Thermo Scientific™ Pierce™ Protein Concentrators PES 100 K MWCO).

After affinity chromatography of BacC4-C5[E-COM-C]-His_6_ and its selenomethionine derivative, the pooled fractions were dialyzed against crystallization buffer X (50 mM HEPES, pH 7.5, 300 mM NaCl,). In the case of the selenomethionine derivate, buffer X was degassed and 2 mM TCEP was added. Size exclusion chromatography (Superdex 200 10/300 GL, Cytiva) with buffer X was carried out to remove protein aggregates. The selected pooled fractions were concentrated to a final concentration of 15 mg/mL using spin columns with a molecular cut-off of 30 kDa (Thermo Scientific™ Pierce™ Protein Concentrators PES 30 K MWCO).

All steps were performed at 4°C. Protein concentrations were determined using calculated extinction coefficients at 280 nm.

### Protein crystallization

Elongated, rod-shaped crystals were grown at 12°C in a hanging drop vapor diffusion approach from an optimized initial hit derived from a MORPHEUS sparse matrix screen.^[64]^ The final crystallization condition contained 0.09 M halogens mix (Molecular Dimensions, MD2-100(250)-71; 0.3 M, sodium fluoride, 0.3 M sodium bromide, 0.3 M sodium iodide), 0.1 M buffer mix 3 pH 8.7 (Tris-base, BICINE), 36% precipitant mix 4 (Molecular Dimensions MD2-100-84; 25% MPD, 25% PEG 1000, 25% PEG 3350). The crystals were mounted on nylon loops and cooled in liquid nitrogen.

Diffraction data were collected at beam line P13 (PETRA III) at 0.979510 Å and 100 K allowing collection of anomalous scattering from the selenium atoms. The data were processed to 3.29 Å using XDSAPP2 (Ref^[65]^) with Friedel’s pairs as individual reflections. Single-wavelength anomalous diffraction phasing was used to determine the selenium substructure and generate an initial structural model with AutoSol^[66-67]^ that was improved with AutoBuild.^[68]^ The model was further refined in Coot^[69]^ and phenix_refine.^[70]^ The final structure was deposited at Protein Data Bank under accession number PDB 9RDE.

### 4’-phosphopantetheinylation

Apo-BacC4-C5-His_6_ wild type and mutant proteins were converted into their holo-forms by a 40 min incubation at 25°C with 0.05 eq. of 4’-phosphopantetheinyl transferase Sfp (*B. subtilis*) and 50 eq. coenzyme A in the presence of 10 mM MgCl_2_ and 1 mM TCEP. Apo-SBP-TycA and apo-TycB1-His_6_ wild type and mutant proteins were converted into their holo-forms at a concentration of 10 µM by a 3 hrs incubation at room temperature with 0.02 eq. of Sfp and 50 eq. coenzyme A in the presence of 10 mM MgCl_2_ and 2 mM TCEP.

### Asp-Asn dipeptide formation assay

In the following section, activity means ATP-dependent Asp-Asn dipeptide formation. For each holo-dimodule of interest as well as for each wild-type control, two aliquots with one reaction volume containing 7 µM of the respective protein were prepared with 2 mM of L-Asp, 2 mM of L-Asn and 20 mM MgCl_2_. ATP was added at a final concentration of 5 mM to one aliquot, whereas the other was complemented with buffer. After a 50 min incubation at 37°C, an aliquot was withdrawn from each reaction mixture for SDS PAGE analysis. The reaction was quenched and peptides and amino acids were dansylated by adding one reaction volume of a 75 mM dansyl chloride solution in dry *N, N*-dimethylformamide and 1 reaction volume of a 1 M Na_2_CO_3_ solution to the remaining reaction mixture. After a 45 min incubation at 60°C in the dark, the reaction was stopped by acidification with formic acid to a final concentration of 17% (v/v). Precipitates were removed by centrifugation (5 min, 18400 x *g*). The supernatant was used for MS analysis.

To quantify the dansylated products, 3 µL of the supernatant were subjected to LC-MS analysis on a maXis II UHR qTof spectrometer hyphenated to an Ultimate 3000 RS HPLC, which was equipped with a C18 reversed phase column (ZORBAX SB-C18 RR HT, 80 Å, 1.8 µm, 50 mm x 3 mm, Agilent Technologies, Waldbronn, Germany). The analyte solution was injected at 0.6 mL min^-1^ and 1% eluent B (eluent A: 0.1% formic acid in water, eluent B: 0.1% formic acid in acetonitrile) and exposed to a linear gradient reaching 40% B at 6 min and 95% B at 7 min. MS settings: range m/z 120 to m/z 1300, spectra rate 1 Hz, capillary voltage 4500 V, endplate offset -500 V, nebulizer pressure 3.0 bar, dry gas flowrate 8 L min^-1^, dry gas temperature 250°C. The system was controlled using Bruker Compass HyStar 6.0 and otofControl 5.2 (Bruker Daltonik GmbH, Bremen, Germany). After each run, a processing method was automatically carried out, which encompassed a lock mass calibration and generated an extracted ion chromatogram (EIC) for [M+H]^+^ of C_20_H_24_N_4_O_8_S (positive polarity, width ± 0.02, monoisotopic peak only). All signals within 3.9 min to 4.2 min were integrated and results exported to csv files. Data from csv files were manually imported into Excel and used for further analysis. Chemically synthesized standards of L-Asp-L-Asn-OH and D-Asp-L-Asn-OH were purchased from Davids Biotechnologie GmbH (Regensburg, Germany) and used for calibration of product yields by spiking defined amounts into the reaction mixture lacking ATP before the dansylation step.

Wild-type activity was repetitively measured, as well as the background in the absence of ATP. After subtraction of the background values, linear regression of the wild-type values provided the basis for normalization of the mutant values. In a final step, mutant activities were corrected for the actual protein concentrations of each BacC4-C5 construct. To this end, the withdrawn aliquots from each reaction mixture were run on SDS PAGE gels (8%). Gels were digitized and the intensities of the protein bands corresponding to the BacC4-C5 constructs were densitometrically determined using ImageJ 1.54i (Wayne Rasband and contributors, NIH, USA) and used to calculate concentration-based correction factors relative to the wildtype construct (**1**). All mutant activities were measured in triplicates (three samples, three gel lanes). Columns in the respective graphs show the averaged values, error bars indicate the standard error.

### D-Phe-L-Pro-diketopiperazine assay (DKP assay)

DKP assays were performed using 5 µM of holo-proteins SBP-TycA and TycB1-His_6_ in a total volume of 100 µL. For the reaction, L-Phe (1 mM), L-Pro (1 mM), ATP (5 mM) and MgCl_2_ (20 mM) were added, and the reaction mixture was performed for 30 min at 37°C. The reaction was stopped using 400 µL of an nBuOH/CHCl_3_ mixture (4:1) and 200 µL of ddH_2_O. Separation of the product was performed via 20 seconds of vortexing, followed by 2 minutes of centrifuging (15870 x *g*, rt) for phase separation. The organic layer was collected. 400 µL of the nBuOH/CHCl_3_ mixture were added two additional times together with 300 µL of ddH_2_O and the procedure repeated. The combined organic solvents were removed in vacuum and the residue was resuspended in 30 to 600 µL of HPLC buffer A (ddH_2_O with 0.1% trifluoroacetic acid and 5% acetonitrile (v/v)). For HPLC analysis, 20 µL were loaded on a C18 reversed phase column (ZORBAX SB-C18 RR HT, 80 Å, 1.8 µm, 50 mm x 3 mm, Agilent Technologies, Waldbronn, Germany) at a flow rate of 0.4 mL min^-1^ and subjected to a gradient starting from 5% eluent B (eluent A: 0.1% trifluoroacetic acid in water; eluent B: 0.1% trifluoroacetic acid in acetonitrile) to 80% B at 20 min, 80% B at 22 min, 5% B at 25 min and 5% B at 30 min. The absorbance of the eluate was monitored at 210 nm.

### NRPS aminoacylation assays

In a total volume of 30 µL, 7 µM holo-BacC4-C5-His_6_ wild type and mutant proteins were incubated with 2 mM of the respective amino acid and 1 mM ATP for 10 min at 25°C to trigger aminoacyl thioester formation. The reaction was quenched by adding formic acid. The supernatant of the subsequent centrifugation (5 min, 18400 x *g*) was used for MS analysis.

10 µM holo-SBP-TycA and holo-TycB1-His_6_ proteins were incubated with 1 mM of the respective amino acid and 5 mM ATP in the presence of 20 mM MgCl_2_ at 37°C. Aliquots were removed after 30 s, 60 s and 300 s. Each aliquot was quenched by adding formic acid. The supernatant of the subsequent centrifugation (5 min, 18400 x *g*) was used for MS analysis.

### Photo-activated crosslink reaction with TycB1(S5Bpa)

Photo-activated crosslink reactions were performed using 5 µM of holo-SBP-TycA (5 µM) and holo-TycB1(S5Bpa)-His_6_ in a total volume of 20 µL. The reaction mixture was preincubated for 30 min at 37°C and subsequently the crosslink reaction was performed for 60 min at 25°C using UV-light of 365 nm (Herolab UV-16L, 2×8 W). Subsequently, 9 µL of the reaction mixture were mixed with 3 µL of 4x SDS-PAGE buffer, of which 10 µL were analyzed via SDS-PAGE (6% acrylamide gel) and Coomassie staining.

### Densitometric analysis of crosslink bands

Analysis of crosslink bands was performed using Coomassie-stained SDS gels and *GelAnalyzer* software (Istvan Lazar Jr.), basically as previously described.^[35]^ L- and T-form crosslink band intensities were calculated using a rolling ball method and baseline correction in combination with manual integration.

### Tryptic digestion and MSMS analysis of crosslink bands

For tryptic protein in-gel digestions, proteins or mixtures were mixed with 4x SDS loading dye (250 mM Tris/HCl, pH 6.8, 8% (w/v) SDS, 40% (v/v) glycerine, 20% (v/v) β-mercaptoethanol, 0.2% (w/v) bromophenol blue) and heated to 95°C for 5 min. Following separation by SDS-polyacrylamide gel electrophoresis and Coomassie brilliant blue staining, the gel was then destained using 50% (v/v) EtOH and 10% (v/v) acetic acid in H2O. The gel pieces containing the crosslink bands were excised and washed in H_2_O, followed by a further destaining step using 50% (v/v) MeOH in 50 mM ammonium carbonate. Gel pieces were washed with acetonitrile and dried in a vacuum concentrator. Once dried, the gel pieces were reduced by a 25 min incubation with 25 mM DTT in 50 mM ammonium carbonate and subsequently alkylated by addition of iodoacetamide to a final concentration of 55 mM. After another wash and drying step, the gel pieces were rehydrated in 20 µL trypsin solution, consisting of 400 ng pre-warmed trypsin (Promega) in 50 mM ammonium carbonate supplemented with 0.01% ProteaseMax (Promega). After 10 min 30 µL buffer were added and then digested for 1 h at 50°C. After the digestion process the supernatant (50 µL) was collected and acidified with formic acid at a final concentration of 0.1% and analyzed by LC-MS without further dilution.

Tandem MS analysis of tryptic peptides (LC-MS^2^) was performed using an UltiMate™ 3000 RS LC nano system (Thermo Fisher Scientific GmbH) connected to a maXis II UHR-qTOF mass spectrometer with a nano-ESI source (CaptiveSpray with nanoBooster, Bruker Daltonik GmbH). 3.5 µL of each sample were loaded on a C18 trapping column (Acclaim PepMap™ 100, 5 μm, 100 Å, ID 100 μm x L 20 mm, Thermo Fisher Scientific GmbH) at a flow rate of 20 μL/min in 2% eluent B (eluent A: 0.1% formic acid in water; eluent B: 0.1% formic acid in acetonitrile). After 10 min of washing at 2% B, a 180-minute gradient (2 to 40% B, flow rate 500 nL/min) was applied for the separation on a C18 nano column (PepSep TWENTY-FIVE C18, ID 150 µm x L 250 mm, 1,5 µm, Bruker Daltonik GmbH). MS settings: capillary voltage 1600 V, mass range: m/z 150-2200. MS survey scans were performed with a cycle time of 2.5s. After each survey scan, the 10 to 20 most abundant precursor ions with z > 1 were selected for fragmentation using collision-induced dissociation. MS/MS summation time was adjusted depending on the precursor intensity, the precursor isolation window and the collision energy were depending on the precursor m/z and charge. DataAnalysis (v5.3) (Bruker Daltonik GmbH) was used for chromatogram processing and fragment spectra isolation. The resulting mgf files were analyzed using StavroX (version 3.6.6).^[71]^

### Mass analysis of intact proteins

Mass analysis of intact proteins was performed using an UltiMate™ 3000 RS system (Thermo Fisher Scientific GmbH) connected to a maXis II UHR-qTOF mass spectrometer (Bruker Daltonik GmbH) with a standard ESI source (Apollo, Bruker Daltonik GmbH). When necessary, proteins were reduced with 2 mM TCEP at 4°C for 10 min to avoid inhomogeneity issues. Then, samples were acidified using a 10% formic acid solution to reach a pH 2-3 and centrifuged (20,000 x g, 3 min). According to the protein concentration, an appropriate volume of the supernatant was loaded on a C4 column (Advance Bio RP-mAb C4, 2.1 mm x 50 mm, 3.5 µm, Agilent Technologies) at a flow rate of 0.6 mL/min in 5% eluent B (eluent A: 0.1% formic acid in water; eluent B: 0.1% formic acid in acetonitrile). After a desalting period of 7 minutes at 5% B, a steep gradient was applied (5-60% B in 2 min). MS settings: capillary voltage 4500 V, endplate offset 500 V, nebulizer 5.0 bar, dry gas 9.0 L/min, dry T = 200°C, mass range m/z 300-3000. Data were analyzed with DataAnalysis (version 5.2) (Bruker Daltonik GmbH) and deconvolution was performed using the MaxEnt algorithm implemented in the software.

## Acknowledgements

We thank Peter G. Schultz for providing a pEVOL plasmid for the incorporation of Bpa. We thank Anna-Lena Feldberg for help in the initial phase of the crystallization project. We gratefully acknowledge financial support by the DFG (grant MO1073/6-2 to H.D.M.).

## References

[1] R. D. Süssmuth, A. Mainz, Angew Chem Int Ed Engl, 2017, 56, 3770–3821.

[2] M. A. Marahiel, T. Stachelhaus, H. D. Mootz, Chem Rev, 1997, 97, 2651–2674.

[3] K.A.J. Bozhüyük, L. Prave, C. Kegler, L. Schenk, S. Kaiser, C. Schelhas, Y. N. Shi, W. Kuttenlochner, M. Schreiber, J. Kandler, M. Alanjary, T. M. Mohiuddin, M. Groll, G.K. A. Hochberg, H. B. Bode, Science, 2024, 383, eadg4320.

[4] T. Stachelhaus, A. Schneider, M. A. Marahiel, Science, 1995, 269, 69–72.

[5] T. Izore, M. J. Cryle, Nat Prod Rep, 2018, 35, 1120–1139.

[6] J. M. Reimer, A. S. Haque, M. J. Tarry, T. M. Schmeing, Curr Opin Struct Biol, 2018, 49, 104–113.

[7] K. D. Patel, M. R. MacDonald, S. F. Ahmed, J. Singh, A. M. Gulick, Nat Prod Rep, 2023, 40, 1550–1582.

[8] J. C. Corpuz, J. O. Sanlley, M. D. Burkart, Synth Syst Biotechnol, 2022, 7, 677–688.

[9] Y. Katsuyama, A. Miyanaga, Curr Opin Chem Biol, 2022, 71, 102223.

[10] A. Tanovic, S. A. Samel, L. O. Essen, M. A. Marahiel, Science, 2008, 321, 659–663.

[11] J. M. Reimer, M. N. Aloise, P. M. Harrison, T. M. Schmeing, Nature, 2016, 529, 239–242.

[12] E. J. Drake, B. R. Miller, C. Shi, J. T. Tarrasch, J. A. Sundlov, C. L. Allen, G. Skiniotis, C. C. Aldrich, A. M. Gulick, Nature, 2016, 529, 235–238.

[13] M. J. Tarry, A. S. Haque, K. H. Bui, T. M. Schmeing, Structure, 2017, 25, 783–793 e784.

[14] C. M. Fortinez, K. Bloudoff, C. Harrigan, I. Sharon, M. Strauss, T. M. Schmeing, Nat Commun, 2022, 13, 548.

[15] Y. J. Peng, Y. Chen, C. Z. Zhou, W. Miao, Y. L. Jiang, X. Zeng, C. C. Zhang, Structure, 2024, 32, 440–452 e444.

[16] J. M. Reimer, M. Eivaskhani, I. Harb, A. Guarne, M. Weigt, T. M. Schmeing, Science, 2019, 366, eaaw4388.

[17] G. W. Heberlig, J. J. La Clair, M. D. Burkart, Nature, 2025, 638, 261–269.

[18] A. Pistofidis, P. Ma, Z. Li, K. Munro, K. N. Houk, T. M. Schmeing, Nature, 2025, 638, 270–278.

[19] J. Alfermann, X. Sun, F. Mayerthaler, T. E. Morrell, E. Dehling, G. Volkmann, T. Komatsuzaki, H. Yang, H. D. Mootz, Nat Chem Biol, 2017, 13, 1009–1015.

[20] F. Mayerthaler, A. L. Feldberg, J. Alfermann, X. Sun, W. Steinchen, H. Yang, H. D. Mootz, RSC Chem Biol, 2021, 2, 843–854.

[21] J. Rüschenbaum, W. Steinchen, F. Mayerthaler, A. L. Feldberg, H. D. Mootz, Angew Chem Int Ed Engl, 2022, 61, e202212994.

[22] A. L. Feldberg, F. Mayerthaler, J. Rüschenbaum, J. Kröger, H. D. Mootz, Angew Chem Int Ed Engl, 2024, 63, e202317753.

[23] X. Sun, J. Alfermann, H. Li, M. B. Watkins, Y. T. Chen, T. E. Morrell, F. Mayerthaler, C. Y. Wang, T. Komatsuzaki, J. W. Chu, N. Ando, H. D. Mootz, H. Yang, Nat Chem, 2024, 16, 259–268.

[24] D. P. Frueh, H. Arthanari, A. Koglin, D. A. Vosburg, A. E. Bennett, C. T. Walsh, G. Wagner, Nature, 2008, 454, 903–906.

[25] H. D. Mootz, M. A. Marahiel, J Bacteriol, 1997, 179, 6843–6850.

[26] D. Konz, A. Klens, K. Schörgendorfer, M. A. Marahiel, Chem Biol, 1997, 4, 927–937.

[27] C. D. Richter, D. Nietlispach, R. W. Broadhurst, K. J. Weissman, Nat Chem Biol, 2008, 4, 75–81.

[28] C. Hacker, X. Cai, C. Kegler, L. Zhao, A. K. Weickhmann, J. P. Wurm, H. B. Bode, J. Wohnert, Nat Commun, 2018, 9, 4366.

[29] J. Watzel, C. Hacker, E. Duchardt-Ferner, H. B. Bode, J. Wohnert, ACS Chem Biol, 2020, 15, 982–989.

[30] J. Watzel, E. Duchardt-Ferner, S. Sarawi, H. B. Bode, J. Wohnert, Angew Chem Int Ed Engl, 2021, 60, 14171–14178.

[31] D. P. Dowling, Y. Kung, A. K. Croft, K. Taghizadeh, W. L. Kelly, C. T. Walsh, C. L. Drennan, Proc Natl Acad Sci U S A, 2016, 113, 12432–12437.

[32] S. Kosol, A. Gallo, D. Griffiths, T. R. Valentic, J. Masschelein, M. Jenner, E. L. C. de Los Santos, L. Manzi, P. K. Sydor, D. Rea, S. Zhou, V. Fulop, N. J. Oldham, S. C. Tsai, G. L. Challis, J. R. Lewandowski, Nat Chem, 2019, 11, 913–923.

[33] M. Hahn, T. Stachelhaus, Proc Natl Acad Sci U S A, 2004, 101, 15585–15590.

[34] M. Hahn, T. Stachelhaus, Proc Natl Acad Sci U S A, 2006, 103, 275–280.

[35] E. Dehling, G. Volkmann, J. C. Matern, W. Dörner, J. Alfermann, J. Diecker, H. D. Mootz, J Mol Biol, 2016, 428, 4345–4360.

[36] C. D. Fage, S. Kosol, M. Jenner, C. Öster, A. Gallo, M. Kaniusaite, R. Steinbach, M. Staniforth, V. G. Stavros, M. A. Marahiel, M. J. Cryle, J. R. Lewandowski, ACS Catalysis, 2021, 11, 10802–10813.

[37] Z. Lv, W. Ma, P. Zhang, Z. Lu, L. Zhou, F. Meng, Z. Wang, X. Bie, Synth Syst Biotechnol, 2022, 7, 989–1001.

[38] M. Kaniusaite, R. J. A. Goode, J. Tailhades, R. B. Schittenhelm, M. J. Cryle, Chem Sci, 2020, 11, 9443–9458.

[39] C. Chiocchini, U. Linne, T. Stachelhaus, Chem Biol, 2006, 13, 899–908.

[40] S. A. Samel, P. Czodrowski, L. O. Essen, Acta Crystallogr D Biol Crystallogr, 2014, 70, 1442–1452.

[41] W. H. Chen, K. Li, N. S. Guntaka, S. D. Bruner, ACS Chem Biol, 2016, 11, 2293–2303.

[42] N. Abbood, J. Effert, K. A. J. Bozhueyuek, H. B. Bode, ACS Synth Biol, 2023.

[43] L. Gao, W. Ma, Z. Lu, J. Han, Z. Ma, H. Liu, X. Bie, Synth Syst Biotechnol, 2022, 7, 1173–1180.

[44] H. Liu, L. Gao, J. Han, Z. Ma, Z. Lu, C. Dai, C. Zhang, X. Bie, Front Microbiol, 2016, 7, 1801.

[45] Y. C. Cheng, W. J. Ke, S. T. Liu, J Microbiol Immunol Infect, 2017, 50, 755–762.

[46] T. A. Keating, C. G. Marshall, C. T. Walsh, A. E. Keating, Nat Struct Biol, 2002, 9, 522–526.

[47] J. Zhang, N. Liu, R. A. Cacho, Z. Gong, Z. Liu, W. Qin, C. Tang, Y. Tang, J. Zhou, Nat Chem Biol, 2016, 12, 1001–1003.

[48] T. Stachelhaus, C. T. Walsh, Biochemistry, 2000, 39, 5775–5787.

[49] U. Linne, M. A. Marahiel, Biochemistry, 2000, 39, 10439–10447.

[50] R. H. Lambalot, A. M. Gehring, R. S. Flugel, P. Zuber, M. LaCelle, M. A. Marahiel, R. Reid, C. Khosla, C. T. Walsh, Chem Biol, 1996, 3, 923–936.

[51] P. J. Belshaw, C. T. Walsh, T. Stachelhaus, Science, 1999, 284, 486–489.

[52] M. J. Calcott, J. G. Owen, D. F. Ackerley, Nat Commun, 2020, 11, 4554.

[53] H. Peng, J. Schmiederer, X. Chen, G. Panagiotou, H. Kries, ACS Chem Biol, 2024, 19, 599–606.

[54] T. Stachelhaus, H. D. Mootz, V. Bergendahl, M. A. Marahiel, J Biol Chem, 1998, 273, 22773–22781.

[55] E. Dehling, J. Rüschenbaum, J. Diecker, W. Dörner, H. D. Mootz, Chem Sci, 2020, 11, 8945–8954.

[56] J. W. Chin, A. B. Martin, D. S. King, L. Wang, P. G. Schultz, Proc Natl Acad Sci U S A, 2002, 99, 11020–11024.

[57] F. Ishikawa, S. Konno, Y. Uchiyama, H. Kakeya, G. Tanabe, Philos Trans R Soc Lond B Biol Sci, 2023, 378, 20220026.

[58] G. Dorman, H. Nakamura, A. Pulsipher, G. D. Prestwich, Chem Rev, 2016, 116, 15284–15398.

[59] J. C. Kauer, S. Erickson-Viitanen, H. R. Wolfe, Jr., W. F. DeGrado, J Biol Chem, 1986, 261, 10695–10700.

[60] M. Grininger, Nat Chem Biol, 2023, 19, 401–415.

[61] A. M. Gulick, ACS Chem Biol, 2009, 4, 811–827.

[62] J. Wang, D. Li, L. Chen, W. Cao, L. Kong, W. Zhang, T. Croll, Z. Deng, J. Liang, Z. Wang, Nat Commun, 2022, 13, 592.

[63] B. R. Miller, J. A. Sundlov, E. J. Drake, T. A. Makin, A. M. Gulick, Proteins, 2014, 82, 2691–2702.

[64] F. Gorrec, J Appl Crystallogr, 2009, 42, 1035–1042.

[65] K. M. Sparta, M. Krug, U. Heinemann, U. Mueller, M. S. Weiss, J. Appl. Cryst., 2016, 49, 1085–1092.

[66] D. Liebschner, P. V. Afonine, M. L. Baker, G. Bunkoczi, V. B. Chen, T. I. Croll, B. Hintze, L. W. Hung, S. Jain, A. J. McCoy, N. W. Moriarty, R. D. Oeffner, B. K. Poon, M.G. Prisant, R. J. Read, J. S. Richardson, D. C. Richardson, M. D. Sammito, O. V. Sobolev, D. H. Stockwell, T. C. Terwilliger, A. G. Urzhumtsev, L. L. Videau, C. J. Williams, P. D. Adams, Acta Crystallogr D Struct Biol, 2019, 75, 861–877.

[67] T. C. Terwilliger, P. D. Adams, R. J. Read, A. J. McCoy, N. W. Moriarty, R. W. Grosse-Kunstleve, P. V. Afonine, P. H. Zwart, L. W. Hung, Acta Crystallogr D Biol Crystallogr, 2009, 65, 582–601.

[68] T. C. Terwilliger, R. W. Grosse-Kunstleve, P. V. Afonine, N. W. Moriarty, P. H. Zwart, L. W. Hung, R. J. Read, P. D. Adams, Acta Crystallogr D Biol Crystallogr, 2008, 64, 61–69.

[69] P. Emsley, B. Lohkamp, W. G. Scott, K. Cowtan, Acta Crystallogr D Biol Crystallogr, 2010, 66, 486–501.

[70] P. V. Afonine, R. W. Grosse-Kunstleve, N. Echols, J. J. Headd, N. W. Moriarty, M. Mustyakimov, T. C. Terwilliger, A. Urzhumtsev, P. H. Zwart, P. D. Adams, Acta Crystallogr D Biol Crystallogr, 2012, 68, 352–367.

[71] M. Götze, J. Pettelkau, S. Schaks, K. Bosse, C. H. Ihling, F. Krauth, R. Fritzsche, U. Kuhn, A. Sinz, J Am Soc Mass Spectrom, 2012, 23, 76–87.

