## Supporting Figures and Tables for "Structural Basis for a Scaffolding Role of the COM Domain in Nonribosomal Peptide Synthetases"

### Supporting Information

Julia Diecker, Benedikt Hermanns, Jennifer Rüschenbaum, René Rasche, Wolfgang Dörner, Alexander Schröder, Daniel Kümmel and Henning D. Mootz*

Institut of Biochemistry, Department of Chemistry and Pharmacy

University of Münster

Correnstraße 36, 48149 Münster, Germany

### Supporting Figures

***cis-COM domains***

```
                  E-domain                                                                                    C-domain
                  -------                                                                 ----------------------------------------------------
BacC4-5           KAVIRHCAE------REETEKTPSDYGDKGISLDQLEEIKLKYKG-------------------MEIEKIYPLANMQRGMLFHALEDKESQ-AYFEQMAINMKGLIDERLFAETFNDIMERHE
BacA4-5           EKIVDHCVD------KEGSDMTPSDYGDVSLGLEELELIKDKYS-------------------AFQIEKIYPLANMQKGMLFHNAMDQTSG-AYFQQIVIKLKGRVHPDILEESFHEIVKRHE
BacC2-3           EQIIKHCTQ------QTESERTPSDYGDTNISLAELEEIKGKY--------------------RSAIEKIYPLANMQKGMLFHAIEDHTSD-AYFQQTVMDIEGYVDPAILEASFNDIMKRHE
LgrB2-3           RAIIAHCRT------EQAGGYTPSDFPLAELDQNSLDKFIGHNR---------------------LIENVYTLTPLQEGMLFHSLYEQAGG-DYVVQLALKLEH-VNVEAFSAAWQKVVERHA
LgrC2-3           REIMEHCRS------EEAGGYTPSDFPLARLDQRAIDNYVGRDR---------------------SIENVYPLTPLQEGMLFHSLYEHAGG-DYVVQFSMTMHH-VEVDVFQQAWQKVVDRHS
LgrC4-5           REIIAHGQS------EAAGGYTPSDFPLARMDQRALDKYLGQNR---------------------SIENVYPLSPLQGGMLFHSLYEQEGG-DYVVQLAMTVEG-LDVEAFEQAWQKVVDRHS
LgrD2-3           REIVAHCTL------PEAGGYSPSDFPLAVLEQKQIDKHIGFDR---------------------QIEDVYTLSPLQQGMLFHSLYNQDSG-DYVVQFAVTFQN-LDVSVLEKAWQNVLDRHS
FegR2-3           TGLVTHVSR------PGSGGRTPSDFPLVALDQARIEQLEAEV---------------------PGLAEVLPVSPLQEGLLFHALFDEQGTDVYVEQMVLDLEGPLDSVALRASWQALLDRHA
FegR4-5           SGLAAHVAG------PGSGGRTPSDFPLVALDQARIEQLEADV---------------------PGLAEVLPVSPLQEGMLFHALYDEQGTDVYVEQMSTGLEGELDAVALRASWQALLDRHA
FegQ2-3           TGLATHVSR------PGSGGRTPSDFPLVELDQPQIEELETQH---------------------PGLADVLPVSPLQEGLLFHALFDESDTDVYVEQMDLGLDGPLDARALRGAWQALTARHA
FegQ1-2           TGLTVHTSR------PGGGGRTPSDLPLLALTQPQVEELEAGE---------------------PEFVDILPVTPLQEGLLFHAQFDENTRDVYVEQMLLDLRGPLDVRALRASWAAMLDRHP
DptA2-3           EGLVAHARR------PDAGGLTPSDLPLVALDHAELEALQADVTG--------------------GVHDILPVSPLQEGLLFHSSFAADGVDVYVGQLTFDLTGPVDADHLHAVVESLVTRHD
Tcp11_4-5         TGLATHAGD------PRAGGHTPADFDLVEVTQRDVTALEAAAPGLT---------------------DIWPLSPLQRGMLFERALDEDGVDVYQSQRILDLDGPLDADRLRAAWRTLVARHE
Tcp11_5-6         AGLAVHAGD------VRAGGHTPSDFPLTALSGQEVAQIEAAVPDLL---------------------DIWPLSPLQEGLLFHAVD-ERGPDVYASMRTLAVDGPLDVARFRASWQAVLDRHA
```

***trans-COM domains***

```
                            COM^D                      COM^A
                ----------------------------------    -------------------
TycA-B            LAIIAHCTE------KKEVERTPSDFSVKGLQMEEMDDIFELLANTLR          MSVFSKEQVQDMYALTPMQEGMLFHALLDQEHN-SHLVQMSISLQGDLDVGLFTDSLHVLVERYD
TycB-C            LGIIEHCMA------KEEGEYTPSDLGDEELSMEELENILEWI                 MKKQENIAKIYPLTPLQEGMLFHAVTDTGSS-AYCLQMSATIEGDFHLPLFEKSLNKLVENYE
GrsA-B            LAIIEHCVQ------KEDTELTPSDFSFKELELEEMDDIFDLLADSLT          MSTFKKEHVQDMYRLSPMQEGMLFHALLDKDKN-AHLVQMSIAIEGIVDVELLSESLNILIDRYD
SrfAA-AB          LRIIEHCLS------QDGTELTPSDLGDDDLTLDELDKL                  MEIFMSKKSIQKVYALTPMQEGMLYHAMLDPHSS-SYFTQLELGIHGAFDLEIFEKSVNELIRSYD
SrfAB-AC          LMLIRHCTE------KEDKEFTPSDFSAEDLEMDEMGDIFDMLEENLK          MSQFSKDQVQDMYYLSPMQEGMLFHAILNPGQS-FYLEQITMKVKGSLNIKCLEESMNVIMDRYD
LicA-B            LQLIDHCLE------RDGGELTPSDLGDDELTLEELDKL                  MEIFMGKQKIQKVYPLTPMQEGMLYHAMLDPESS-SYFTQLELFIDGEFDLGIFEKSVNQLVRSYD
LicB-C            TALIDHCTA------KEEREFTPSDFSAGDLEMEEMGDLFDVLEENLK    MKGDTILSQFKKEHVQDIYYLSPMQEGMLFHTLLHPGQS-FYIEQISMQVKGSFQKDILEKSMNVIIGRYD
BacB-C            QMVISHCTG------KHETEKTSSDYGYDKLSLEDLEELLNEYESVDS              MKTKVEKIYPLSNMQKGMLFHAMKDEASH-AYFEQFIIELKGDVDERMFEESLNEVMKRHE
LgrA-B            NALLAHCLQ------KTETELTPSDFVDKNLSRSELDDIMDLISDL               MSRKKVDNIYPLTPMQEGMLFHSLLDEGSE-SYFEQMRFTIKGLIDPAILEQSLNALIERHD
LgrB-C            QSLIQAEKQSYRAEDFEDADLSQSALNKVLARLKNRKGNELHGGSH              MNSMGDLTDLYTLTPLQEGMLFHSLYS-EGS-AYMIQTTAILTGELDIVPFEKAWKKVIQRHS
LgrC-D            RQLLAHAKT------DKAAAPAQSAASNLSEFGWDDEEIADLLDLIQDK              MNNIETYYPVTPLQQGLIFHSLLEPESG-AYIVQMGLKLQGPLNIPLFEQAWQCLVDRHA
FengA-B           EDIIAHCSG------KKEREKTLTDYSHTELTAQALSSIEDLVKGL              MTQATEIQDIYPLSYMQEGMLFHSLLDSGSN-AYVEQISFTVSGDLCIDTFQKSLDMLVSRYD
FengB-C           KQIISHCTG------KEEREWSAADFSDEELTLEDLSDIMGAVNKL              MGQQPEIQDIYPLSFMQEGMLFHSLLDHDSR-AYFEQAVFTINGQLDRERFQKSIDAVFERFD
FengC-D           RDIMAHCTG------KEEAEKTVSDFSSKTLTSDDLDSIASFVEEL              MTKKNAIQDIYPLSYMQEGMLFHSLLQKESQ-AYAEQASFSITGKVDTRVFEESVHALFDRHD
FengD-E           LKLSEHCLS------KTDTEKTVSDFDDRELTEEALQDIADLLSFH              MDKTKNIQNIYPLSRMQEGMLFHSFLQKEGA-AYIEQSVFTIKGRLRPELFEQSVQSVISRHD
PpsA—B            IETIEHCSQ------KKAREKTLSDFSNKELTLSALSSIEDLVKDL              MAQSAQIQDIYPLSHMQEGMLFHSLMDFSSK-AYIEQTSFTITGNLCVDSFQKSLNLLVSRYD
PpsB-C            ENIIEHCTG------KENQEWSASDFTDEDLTLDELSEIMGAVNKL              MPQQPEIQDIYPLSFMQEGMLFHSLYDEQSR-AYFEQASFTIHGQLDLERFQKSMDAVFDRYD
PpsC—D            LDIMKHCAG------QQKAEKTLSDFSSQSLTAEDLDSISSLVEEL              MTKANSIQDIYPLSYMQEGMLFHSLLQKDSQ-AYVEQASFTIEGKVNPQFFQNSINALVERHD
PpsD—E            LQLIEHCQN------KSETEKTISDFDDQELTEDALQEIADMLSFH       MKKGADTMNTIKKIKNIYPLSHMQEGMLFHSFLRKEEG-AYVEQSLFTIKGSLSYDWFQRSIQAIIDRHD
FegR-S            TGLVTHTSD------AGSGGYTPSDFPLVDISQDELDEFEAMAKQIDEGA           MSDSALVGVWPLSPLQEGMLFHAVYDEHGTDVYVEQISTGLEGRLDTVVLRASWQAVLDRHE
Tcp9-10           SGLAGQADD------PSAGGHTASDFDLLDLDQDEIENFEAIAAEFGGGQES   MTTPTTSPGSALAEVWPLSPMQEGMLYHASFDDAAPDIYVIQQSQIIDGPLDTERFRRSWEVLLHRHA
                                       : :           .  .                                 :: :* *:::.          :       .   .    :      :   .
```

>>> Figure S1 continued on next page

```
                                                                    C-domain                                          >>> (seq. truncated)
                  ------------------------------------------------------------------------------------------------------------------------
                            COM^A                                               COM^A
                     ------------------                                           ----
BacC4-5           ILR--ASIEYEITDEPRNVIIKDRKINLDYHDLRKQSPAEREQVIQAYRKADREKGFRLNSEPLIRAALMRTEDDSYTFIWTNHHILLDGWSRGIIMGELFHMYHMKEARQKHRLEEARP
BacA4-5           ILR--ASFEYEITAEPRQIIARDRKTPFTSIDLTGENRTRQHRFIETYLKEDQEKGFDLSSEALMRVCLIKMSDESYRLIWSHHHILLDGWCLGIVLSELFSLYGKIMKGESRRLKEPKP
BacC2-3           ILR--ASYEYEIVEEPRQIIIENRSIDFTYFNIAKSSAQQQEMFIERLLNEDRKKGFDLSKDVLMRAYLLKTAERSYRLVWSHHHILLDGWCLGIIMRELFVIYENRMNGKASPLKETKP
LgrB2-3           ILRTSFLWSG--LEKPHQVVHAKVKTFVERLDWRHLTAAEQEAGLQTYLEQDRKRGFDLARPPLMRWTLIRLDASTFQFVWSFHHMLLDGWSTPIVFQDWQAFYAAASHGKEASLPAIPP
LgrC2-3           ILRTSFIWDG--VSTPHQIVRKHVQVVVDEQDWRHVPADQQKAEWDAFLEEDRKRSFAITEPPLMRWTLLRISDTAYRFIWSFHHVLLDGWSVPLVMKDWFAAYMALADGKDIQFGAVHP
LgrC4-5           ILRTSFIWEG--LTEPHQVVRKQVKACVEKIDLRHLTPDQQKAELSEYLAADRRRSFEIAVAPLMRWTLFRLSESAYRFTWSFHHVLLDGWSIPIVLKDWFSAYLSLAEGKEVAHSFVQP
LgrD2-3           ILRTHFVWEG--LSEPHQVVRKDVKVTLTKEDWRHLQADVQDEMLAAFLEEDRRRSFDIAQAPLSRWVVFQTKDEEYRFVWSFHHVLLDGWSVPIVLNELLAHYAAISEGREGKLVPSQP
FegR2-3           SLRAGFRQLDG-LDEPVQVIARSVTLPWREVDLSALDGEAASAAAERLAGEDQARRFDVAVPPLLKVMLVKVGPDRYRMAVTLHHILLDGWSLPILMQELWSAY--EAGGSVAALPRVTP
FegR4-5           SLRAGFRQLDG-VDQPVQVIAQGVTLPWREVDLSALAEDEALAEAERLSAEERDLGFRMTVPPLLKVLLVKVGADRYRMSVTLHHILLDGWSLPIMMRELWDAY--AAGGSAAGLPAVTP
FegQ2-3           SLRAGFRQLPG-VEQPVQIVAREVTLPWREVDLSALPENEALAEAARLGVEERARRYDLSAPPLLRILLVKVGETRHRMMVTLHHILLDGWSLPILTRDLWAAY--AAGGSTTGLPPVIP
FegQ1-2           ALRAGFRRIPG-LKQPVQTITARATLPWREEDLSALGRDAAHAEARRLEDEERARRFDVQQPPLVRVLLTALGDGHHRMVVTLHHLVLDGWSLPVLVRELWSAY--DAGGDAGALAPVTP
DptA2-3           VLRTGYRQAQ--SGEWIAVVARQVHTPWQYIHTLDT-------DADTLTNDERWRPFDMTQGPLARFTLARINDTHFRFIVTYHHVILDGWSVAVLIRELFTTYRDTALGRRPEVPYSPP
Tcp11_4-5         SLRTSFHQLE--SGETVQVVVDQADIGWRVADVSHRAEADAAAEVGRLLAEDQAQRFDVTRAPLLRLLLVRLGADRHRLVVTSHHIVLDGWSTPIILGEMSAAY--AGAPSTATAPSP--
Tcp11_5-6         ALRASFHQLE--SGAAVQAIAREVTLPWQETDLSDLPEDVALAEFDRLAAQLRDERFDLTRAPLLRLHLVRLGERRHRLAFASHHISCDGWSLPIVSTEVMAAY--EGR----GLPAPTS
TycA-B            VFRTLFLYEK--LKQPLQVVLKQRPIPIEFYDLSACDESEKQLRYTQYKRADQERTFHLAKDPLMRVALFQMSQHDYQVIWSFHHILMDGWCFSIIFDDLLAIYLSLQNKTALSLEPVQP
TycB-C            VLRTAFVYQN--MQRPRQVVFKERKVTVPCENIAHLPSAEQDAYIQAYTK--QHHAFDLTKDNLMKAAIFQTAENKYRLVWAFHHIIVDGWTLGVLLHKLLTYYAALRKGEPIPREATKP
GrsA-B            VFRTTFLHEK--IKQPLQVVLKERPVQLQFKDISSLDEEKREQAIEQYKYQDGETVFDLTRDPLMRVAIFQTGKVNYQMIWSFHHILMDGWCFNIIFNDLFNIYLSLKEKKPLQLEAVQP
SrfAA-AB          ILRTVFVHQQ--LQKPRQVVLAERKTKVHYEDISHADENRQKEHIERYKQDVQRQGFNLAKDILFKVAVFRLAADQLYLVWSNHHIMMDGWSMGVLMKSLFQNYEALRAGRTPANGQGKP
SrfAB-AC          VFRTVFIHEK--VKRPVQVVLKKRQFQIEEIDLTHLTGSEQTAKINEYKEQDKIRGFDLTRDIPMRAAIFKKAEESFEWVWSYHHIILDGWCFGIVVQDLFKVYNALREQKPYSLPPVKP
LicA-B            ILRTVFVHQQ--LQKPRQVVLAERQAIVEFEDLADLDEEKQNNRIDRYKQEVQAAGFNLAKDMLFKTAVFRLDRNKYLLVWSNHHIVMDGWSMGILMKRLFQNYEAFRANRTVPLDQGKP
LicB-C            IFRTVFVHEK--MKRPVQVVLKERSFQAEEIDLSGLSEAEQNERIEDYKRKDKEKGFNLSKDIPMRTAVFKKGQDRYEWVWSYHHILLDGWCFGIVVQELFEVYNALRENRLYSLGPVKP
BacB-C            ILR--ASFHHRLD-EPLHVIIKDRHMKFDYLDIRG--RHDQDGVLERYLAEDRQKGFDLAKDTLMRACLIRMSDDSYQFVWTYHHILLDGWCLGIILDELLTIYEMKRKGQNHQLEDPRP
LgrA-B            ILRTVFLLEK--VQKPRQIVLRERKTKVQVLDITHLSEGEQAAYLEDFAQKDRQASFDLAKDVLIRLTLVRTSADTHTLFWSHHHILLDGWCIPIVLNDFFQIYQQRKGGLPVELGPVYP
LgrB-C            ILRTGFIWEE--TEKPLQAVFESVPFSIRQKDWSSYGSDEQESMLAAFLQNEKAAGFDLSEAPLMRVTIIKLGEAVHRLIWSFHHLLLDGWSSPIVFQEVLDFYEAYRQGKDLRLPQARP
LgrC-D            IFRTRFVGGK--VKEYVQVVLKDLKISLVEHDLIHLSSSEQEAFLHHFAKEDRKRGFDIEQAPLMRLNVFHLNSETVHFLWTLHHVLIDGWSMPLVFGEVFAAYEMLSKGQPLSLPPVRA
FengA-B           IFRTIFIKEVPDLEGPQQVVLSRRDTAVRTEDISDYSEERQQSIIEEFKETDRHKGFDLQKGPLMRLTLFQTGENRHTCVWTHHHIMMDGWSLGIVLKDFFSMYYANRNGRPVNLGSPAP
FengB-C           IFRTTFIHKN--VAKPRQVVLKNRPSRVQFHDISHLDEKAQDKYTDRFRKEDKDKGFDLQSDPLMRVSILKKAPERYVCIWSHHHIVMDGWCFGIVMKEFLMIYQSLGDGRLPSLEPVQP
FengC-D           IFRTIFISQN--VSVPQQVVLKERNVSIIEENLTNLNKADQIKHIEEAKRRDRKKGFHLQKDMLMRVTLLQTGECEYTCIWSFHHIIMDGWCLGIVLKEFFQIYASRLRRTPLTLEPAVP
FengD-E           IFRTVFLPHVAQLNGPRQVVLREREFRLHREDLTHLDEAEQSIYLQRFKERDRLKGFDLQKDMLMRVSLFKTADEKYICVWSHHHILMDGWCLGIVLQEFMHMYRAIESGSPVTLAPAKP
PpsA—B            IFRTIFIKEVPDLTGPQQVVLSNRELTVYREDISRLADQEQQTLIDAFMTKDREKGFDLQKDPLMRLALFDRGDSQYTCVWTHHHIIMDGWCLGIILKEFFSMYDSLKNNSPVQLGSTVP
PpsB-C            IFRTAFIYKN--VAKPRQVVLKQRHCPIHIEDISHLNERDKEHCTEAFKEQDKSKGFDLQTDVLMRISILKWAPDHYVCIWSHHHILMDGWCLGIVIKDFLHIYQALGKGQLPDLPPVQP
PpsC—D            IFRTIFISQN--VSSPQQVVLRERNVIVLEEDITHLNEAEQSQFIEQWKEKDRDRGFHLQKDVLMRIALIQTGESQYSCIWTFHHIMMDGWCLSIVLKEFLHIYASYVNASPITLEPVQP
PpsD—E            IFRTVFLPHVPHLSGPRQVVMTEREFHLNSEDISHLPTNDQNEYIERFKEKDKQKGFDLQKDMLMRISLFKTAKDEHVCIWSHHHILMDGWCLGIVMQEFMQIYQSIHAGKPLSLDPVRP
FegR-S            SLRAGFQRRS--SGDPVQLIRRRVVLPWREEDLSALPEEEALGEAERLSTEEQAQGFDMTVPPLLKVLLVKVGQDRYRMSVTLHHILLDGWSLPILMRELWTCY--EAGGSAEGLPAVTP
Tcp9-10           ALRASFHRRK--SGETVQLIPREVRLPWAERDLSGLPEKAALAEVGEIAAKERAERFDLTKPPLLRLMLIRLGPQRHCLVTTSHHLLMDGWSRAILESELLHVY--ASGGTVSGLPPAGS
                   :*                :           .                        : :      :  :            : **:  ***   ::       *
```

Figure S1. Multiple sequence alignment of *cis*- and *trans*-COM domains. Shown are selected sequences aligned with CLUSTALW. Structural elements of the *cis*- and *trans*-COM domains of BacC4-5 and TycA/TycB, respectively, are highlighted in bold and shaded letters.

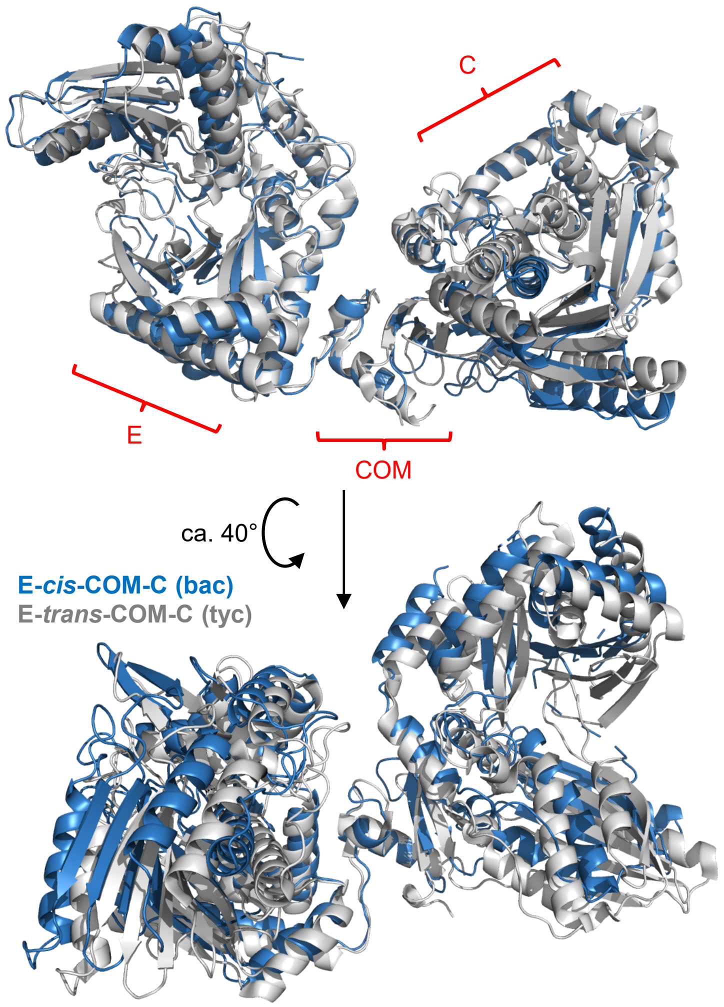

Figure S2. Comparison von *cis*-COM and *trans*-COM ensembles with neighboring E and C domains. Crystal and cryo-EM structures depicting the E-*cis*-COM-C (this study) and E-*trans*-COM-C structures (pdb-code: 9BFD, Ref[1]), respectively, were overlayed using the $COM^D$ parts. Note that the positioning of the E and C domains relative to one another is highly similar for both domain ensembles, further underlining the conserved structural role of the COM domain in orienting its neighboring domains.

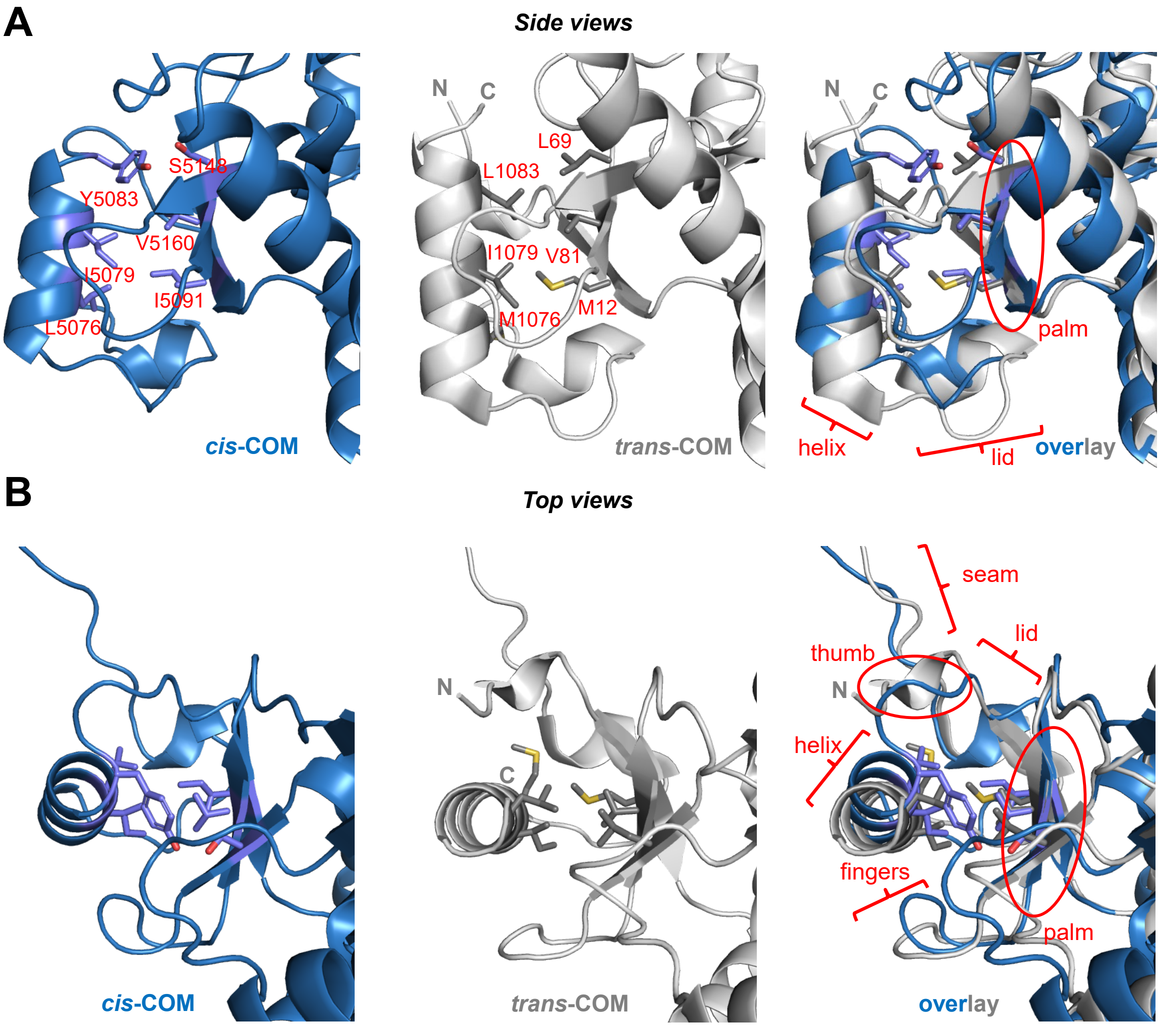

Figure S3. Binding modes in helix-hand interaction. Shown are parts of the crystal and cryo-EM structures depicting the *cis*-COM (this study) and *trans*-COM structures (pdb-code: 9BFD, Ref[1]), respectively. In the illustrations the COM domains are shown starting at the beginning of the seam region (the N-terminally located E domains are not shown) and with most of the body of the C-terminally located C domains being cut off at the right-hand sides of the panels. The central three residues from both the helices and the palm regions of each helix-palm interface are highlighted as stick representations of the side chains with the corresponding residue numbers given in the two upper left panels. The overlayed representations (right panels) were created by overlaying all residues of the three palm β-sheets from the two structures to reveal how the helices are oriented relative to the palm region. Note that the two helices are in differently staggered arrangement relative to their palm interaction interface and show a slightly different tilt of their helix axis relative to the palm region. Together with the additional helix turn found for the central helix of TycA (*trans*-COM), this arrangement results in slight differences of the positioning of the N-terminal ends of the two helices and slight differences in the folding of the lid regions. These differences might represent minor folding differences within COM domains, in particular when comparing COM domains with varying numbers of amino acids (see Figure S1). A) Side views with angle perpendicular to the central α-helices. B) Top views along the helix axis with the N-terminal end of the helices facing the viewer. Images were created using PyMol.

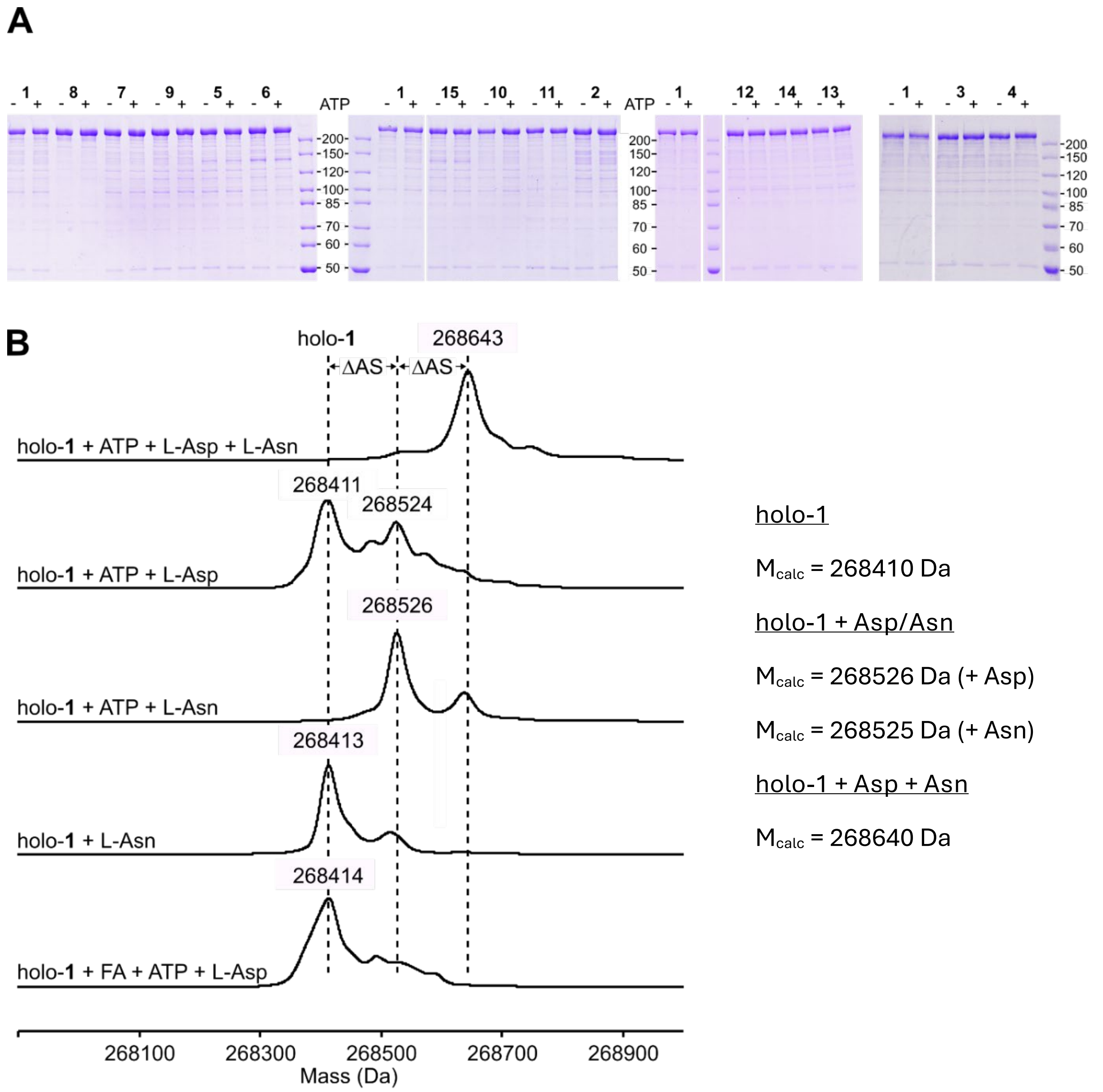

Figure S4: Dimodular BacC4-C5 constructs. A) Coomassie-stained 8% SDS PAGE gels of purified and Ppantylated holo-BacC4-C5 proteins **1** to **15**, with amino acids L-Asp and L-Asn and with or without ATP as indicated. These gels were run from removed aliquots of the dipeptide assays and also used to correct the amounts of formed dipeptide to the actual concentrations of the holo-BacC4-C5 proteins. B) Aminoacylation assay of holo-BacC4-C5 (**1**). Shown are mass spectra of the intact proteins (ESI-Q-TOF). Protein samples were ppantylated prior to the incubation with the respective substrates. Holo-**1** (7 µM) was then mixed with amino acids (2 mM each) and ATP (1 mM), as indicated, and incubated for 10 min. The protein concentration during the 10 min incubation period was 7 µM. Negative controls either lacked ATP or were acidified prior to the addition of ATP by addition of formic acid (quench control).

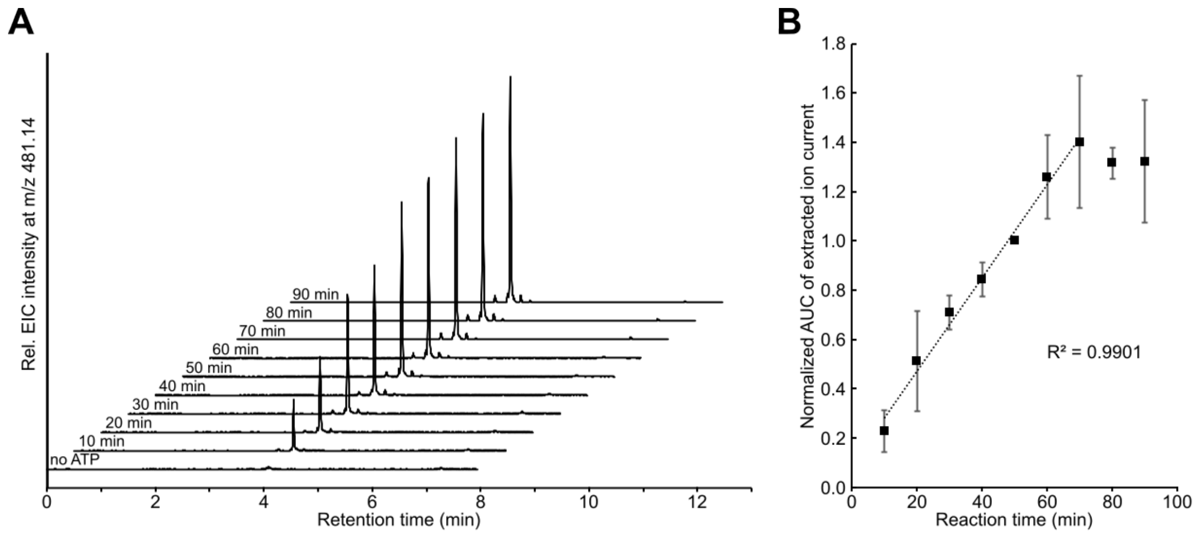

Figure S5: Time-course of product formation of holo-BacC4-C5 (**1**). A) EICs of $[M+H]^+$ of isobaric compounds with $C_{20}H_{24}N_4O_8S$ are shown, corresponding to the dansylated di-peptides D-Asp-L-Asn and L-Asp-L-Asn. Assay conditions: Incubation times were varied from 10 min to 90 min in the presence of 2 mM L-Asp, 2 mM L-Asn, 5 mM ATP (0 mM ATP in control) and 7 µM holo BacC4-C5 protein (**1**). After incubation, samples were quenched and dansylated for 45 min at 60°C. B) Averaged areas under the curve of three repeats of the experiment shown in A. AUCs show linearity with $R^2$ = 0.9901 for points between 10 min and 70 min. EIC = extracted ion count.

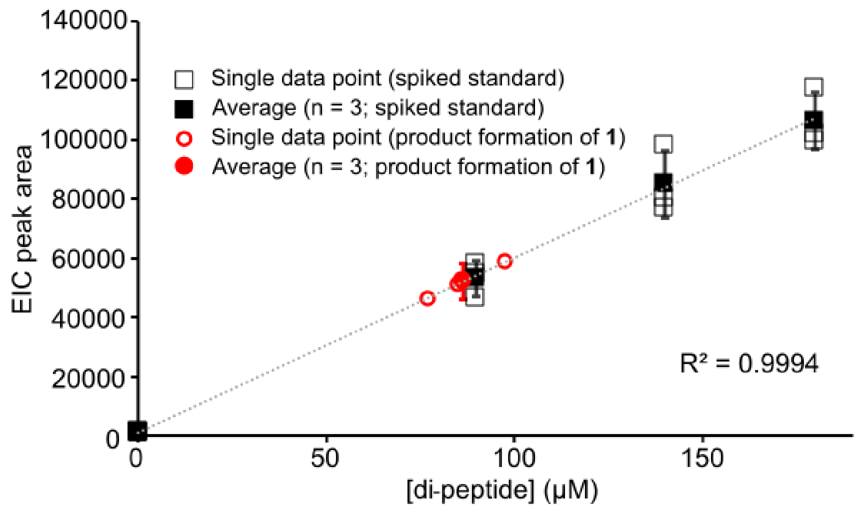

Figure S6: L-Asp-L-Asn standard calibration and product formation measurement. The dipeptide standard was spiked at 0 µM, 90 µM, 140 µM and 180 µM (open rectangles: single data points; filled black rectangles: average of of three technical repeats; error bars: standard error) into assay samples with 7 µM holo-BacC4-C5 WT, 2 mM L-Asp, 2 mM L-Asn lacking ATP. Enzyme activity samples contained no standard but 5 mM ATP (open circles: single data points; filled red circle: average of three technical repeats; error bar: standard error). Dansylation was carried out after 50 min incubation at 37°C. The whole calibration and product formation procedure was carried out three times. Within the incubation period holo-BacC4-C5 (**1**) produced 101.7 ± 22.3 µM L-Asp-L-Asn (n = 3), corresponding to about 1 molecule in 3 min per enzyme (0.29 $min^{-1}$), when calculating with the assumed enzyme concentration of 7 μM.

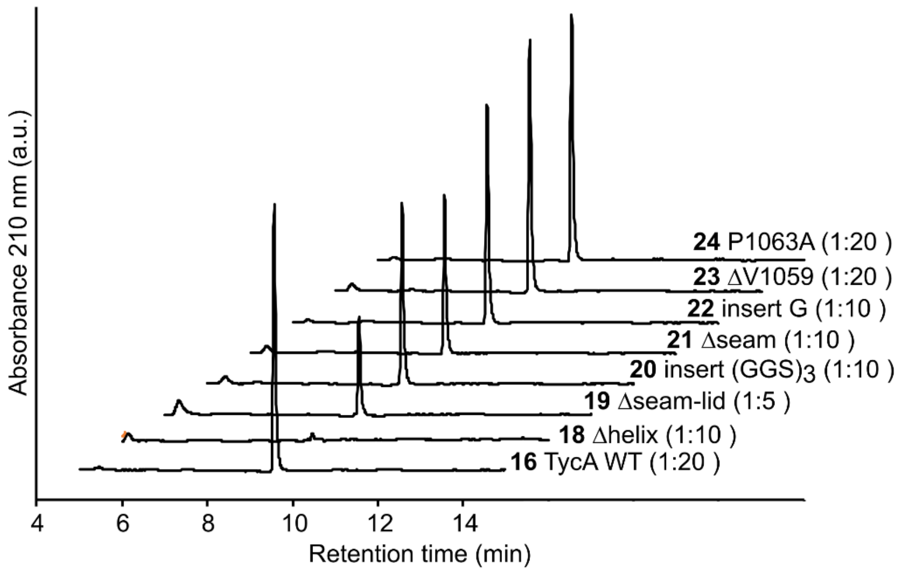

Figure S7. HPLC chromatograms of DKP formation of TycA WT and indicated mutants with TycB1. Numbers in brackets indicate the dilutions (in HPLC buffer), with which the samples were analyzed by HPLC.

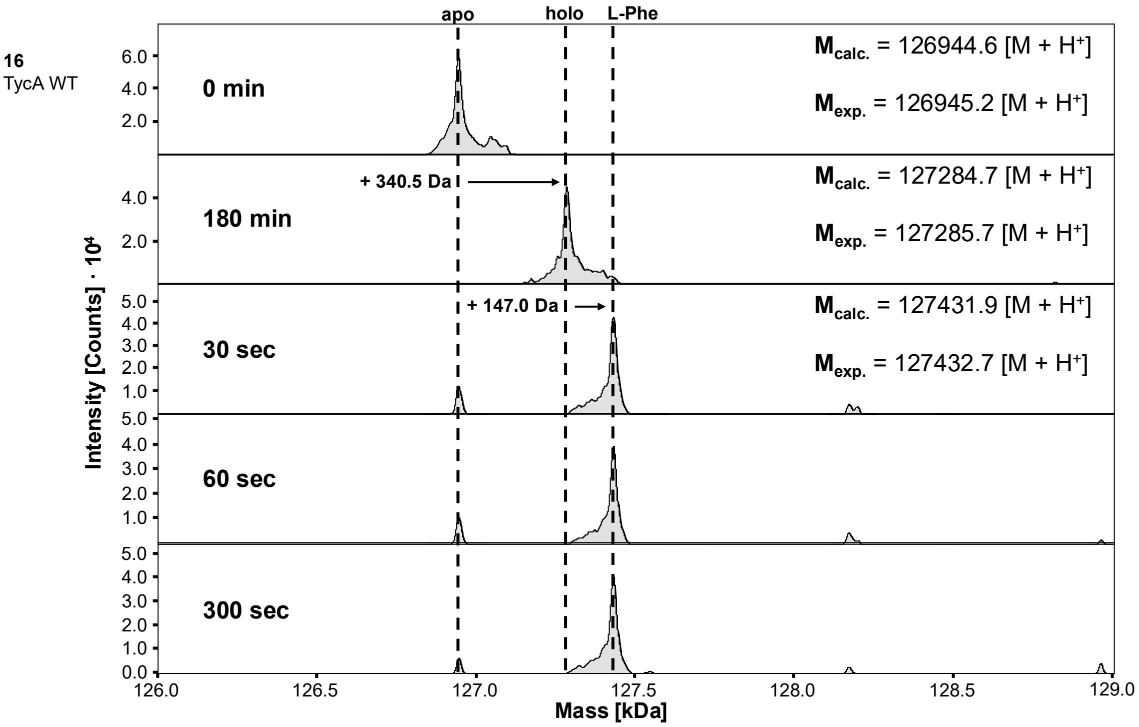

Figure S8 >>> continued

**18**
Δhelix

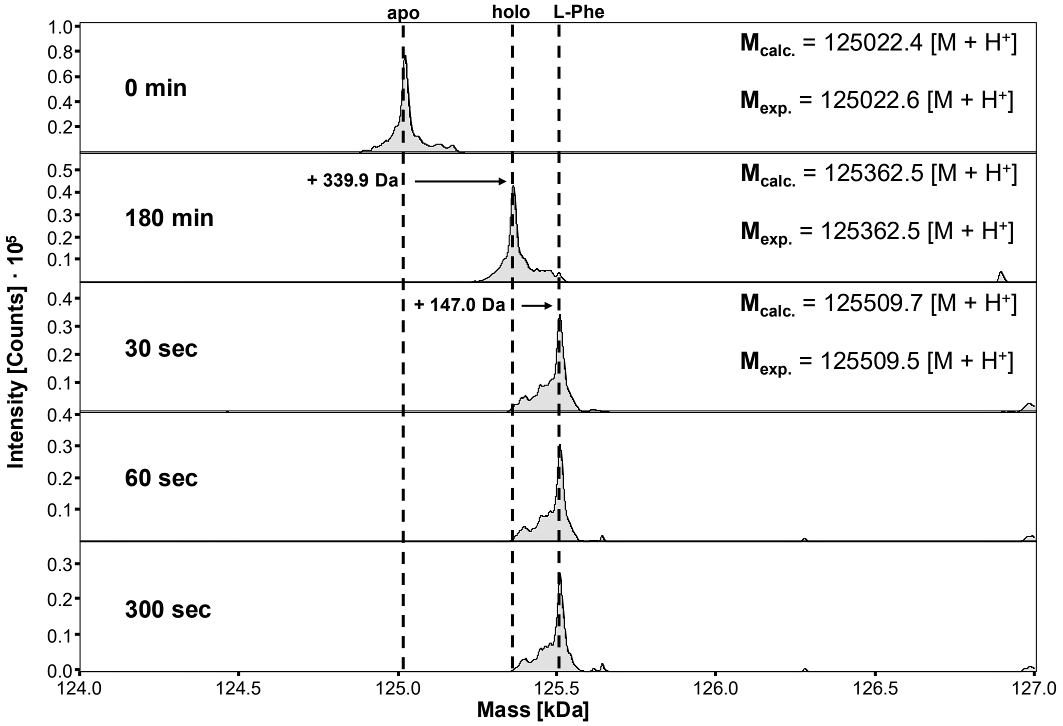

**19**
Δseam-lid

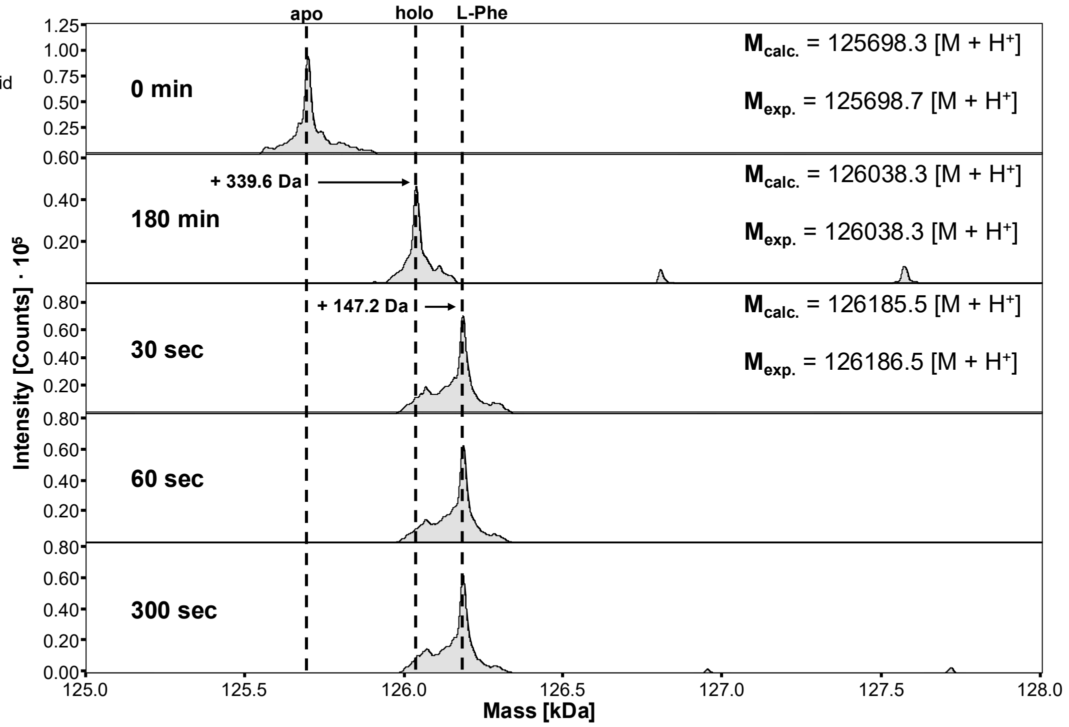

Figure S8 >>> continued

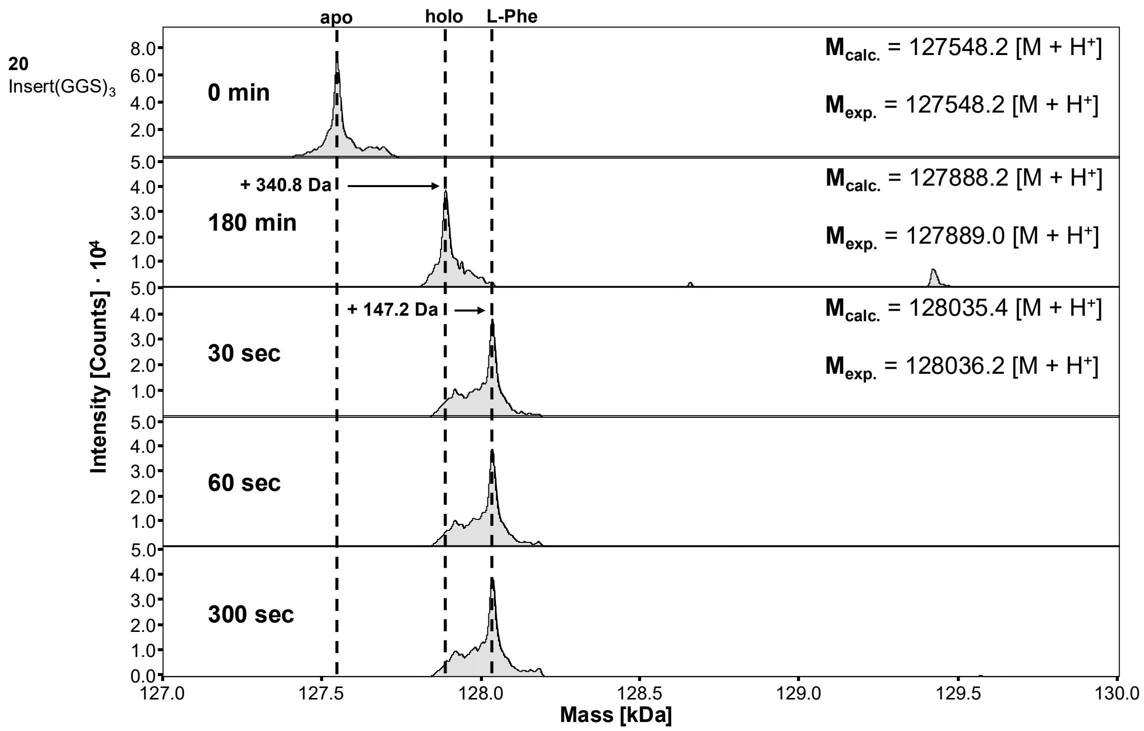

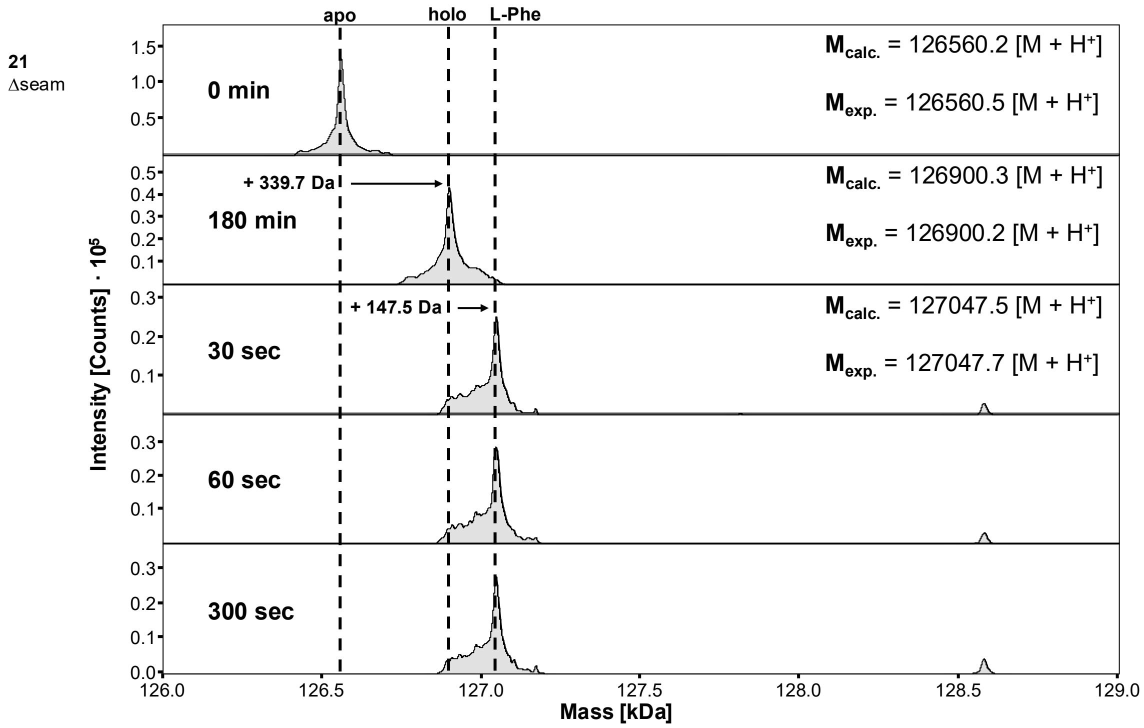

Figure S8 >>> continued

22
Insert G

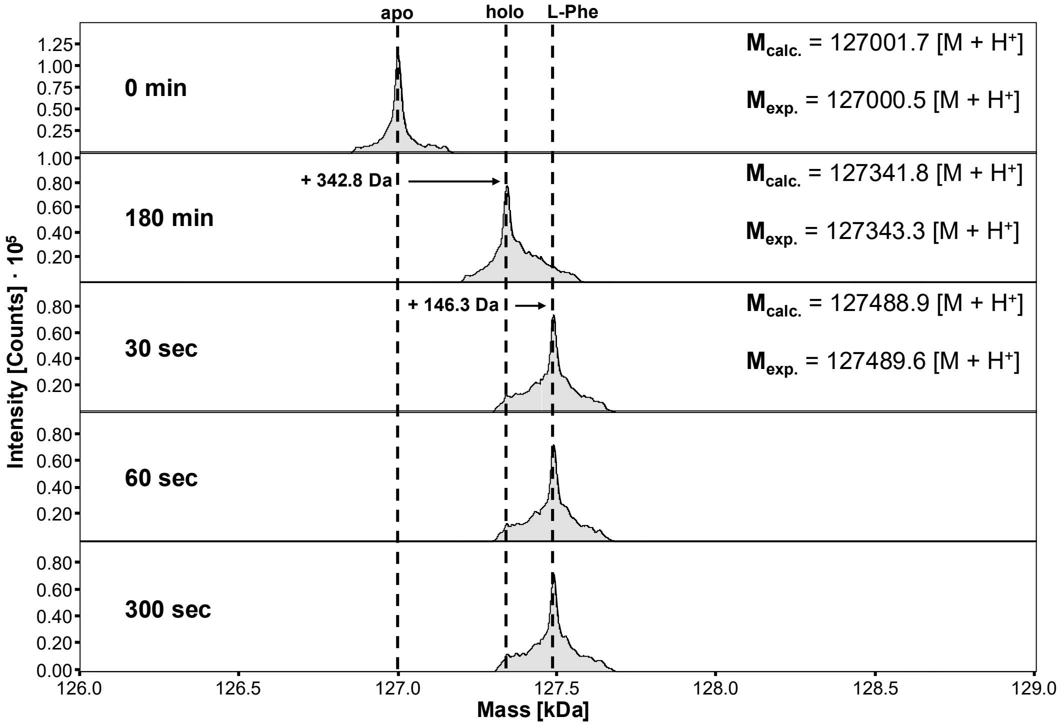

23
ΔV1059

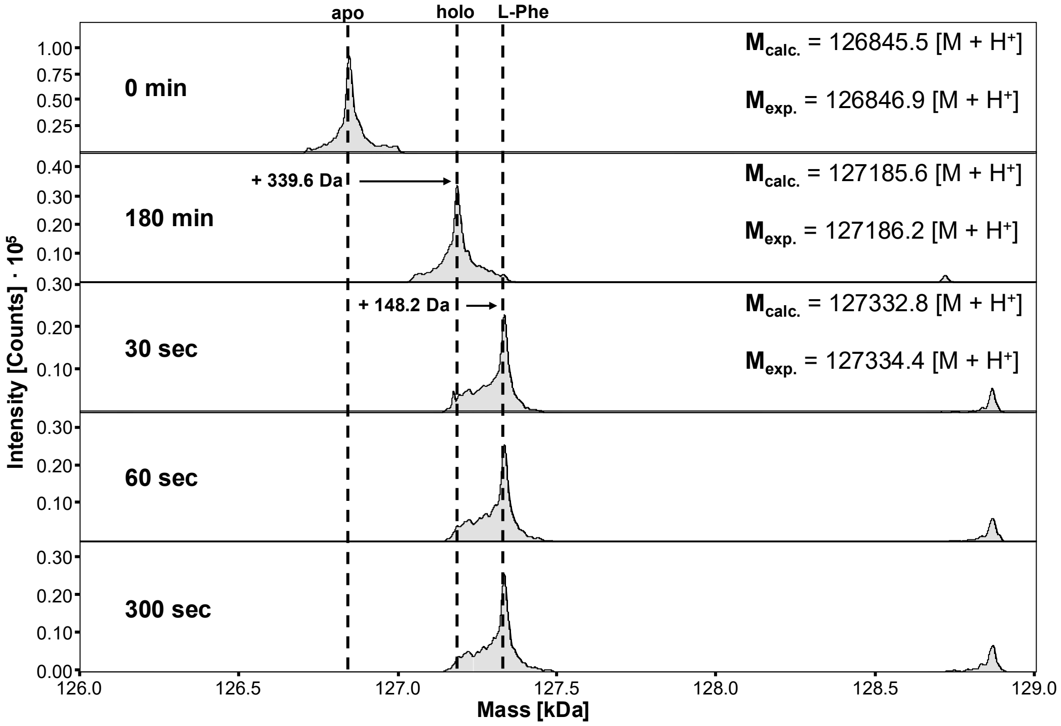

Figure S8 >>> continued

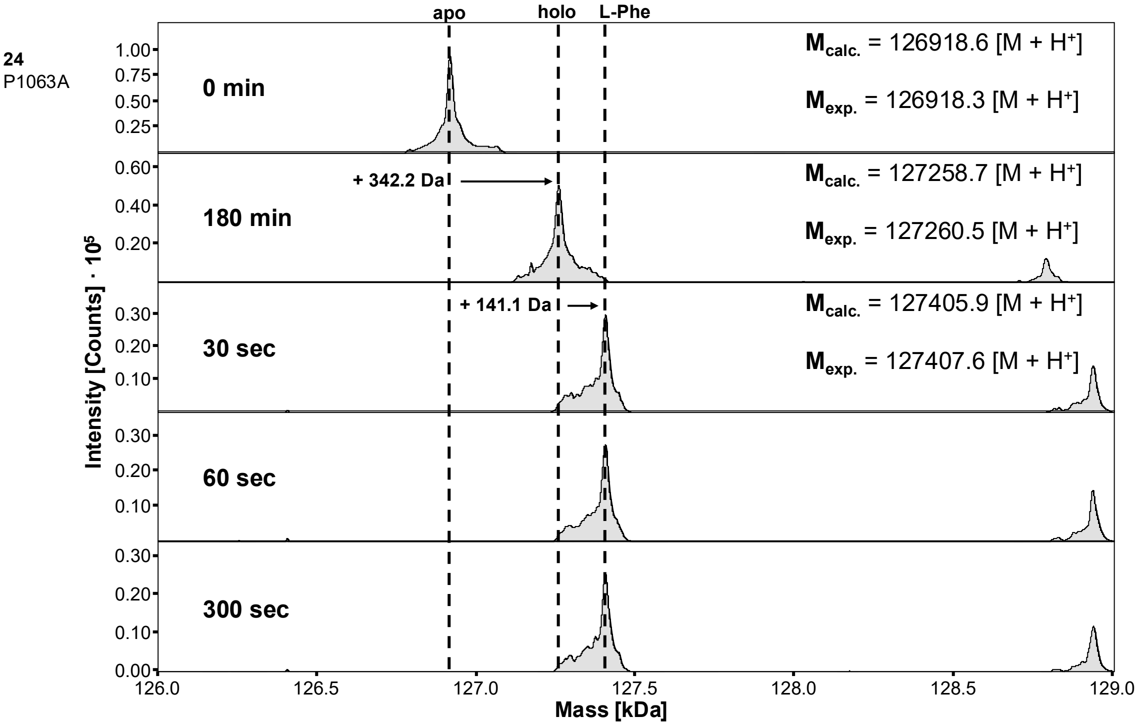

Figure S8. Aminoacylation activity of SBP-TycA (**16**) and its COM mutants **18-24**. The indicated proteins were analyzed by ESI-MS as purified apo-proteins (top spectra; 0 min), following 3 h of 4'-Ppantylation with Sfp and CoASH to convert them into the holo-forms (second spectra from top; 180 min), and at 3 time points after addition of ATP and L-Phe to following the formation of the aminoacyl thioester (three spectra from bottom; at 30 sec, 60 sec, and 300 sec). These data show that all mutants retained the virtually full aminoacylation activity of the wild-type protein **16** after 30 sec.

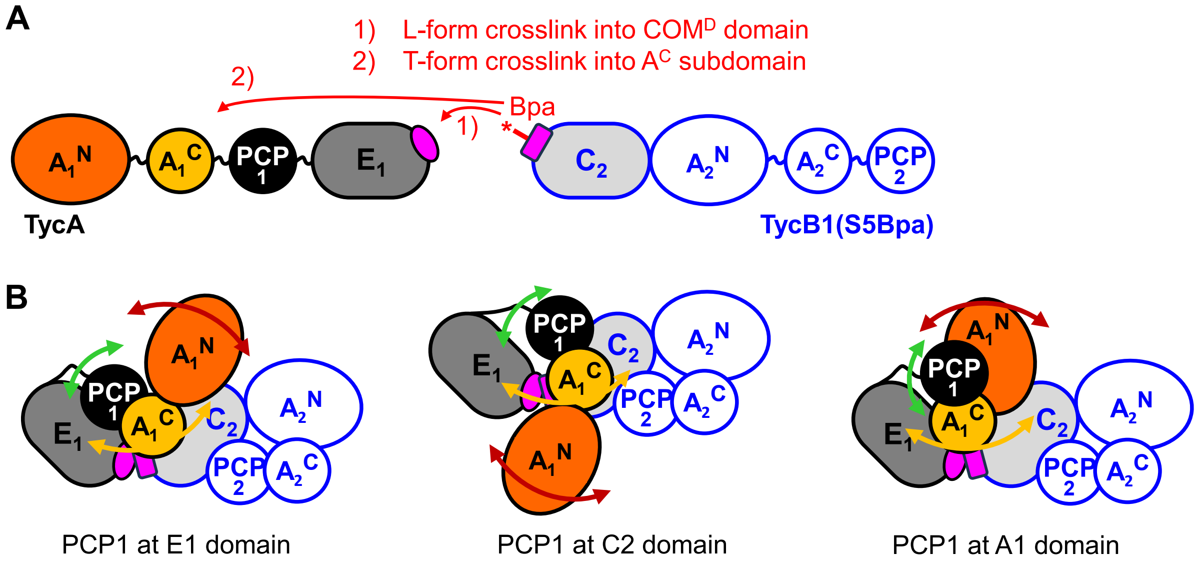

Figure S9. L- and T-crosslink formation in photo-crosslinking between TycA and TycB1. A) Linear representation of the multi-domain TycA and TycB1 module with the photo-crosslinking amino acid Bpa inserted at position S5 of TycB1, corresponding to the thumb region of the COM$^{A}$ domain. Photo-crosslinking results in proximity-driven covalent bond formation with 1) either the COM$^{D}$ domain of TycA or 2) the A$^{C}$ subdomain of TycA. B) Illustration of theoretically possible 3-D domain arrangements in the associated TycA/TycB1 protein complex to explain the spatial proximity between the A$^{C}$ subdomain of TycA and the COM$^{A}$ of TycB1 domain, giving rise to the T-form crosslink. Three different complexes depict the interaction of PCP1 with each of its catalytic domain partners. With a PCP1 positioning in the interspace between E1 and C2 domain, as supported by the E-COM-C structure, the spatial proximity of the $A^{C}1$ domain to S5Bpa, which would explain the T-form crosslink, is plausible in each case given the dimensions of the domains and the flexibility of the linkers between them. Note that the preferred exact location, if any, of the $A^{N}1$ and $A^{C}1$ domains in general is not known. Likewise, the positioning of the PCP1 domain relative to the E domain cannot be extrapolated from known structures in case of the PCP1-$A^{N}1$/$A^{C}1$ interaction (right panel).

A

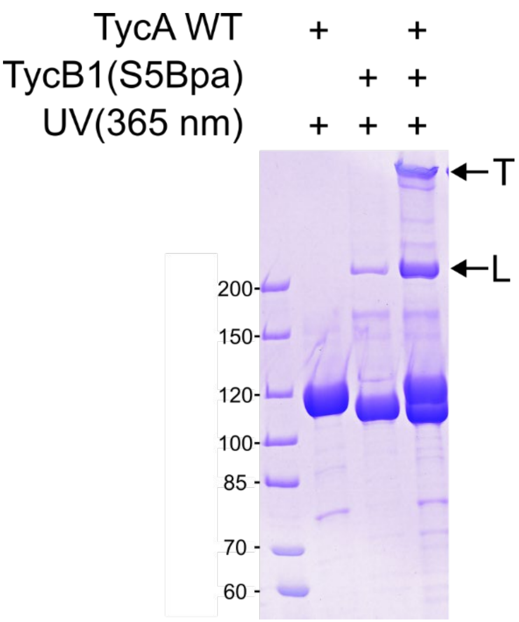

B

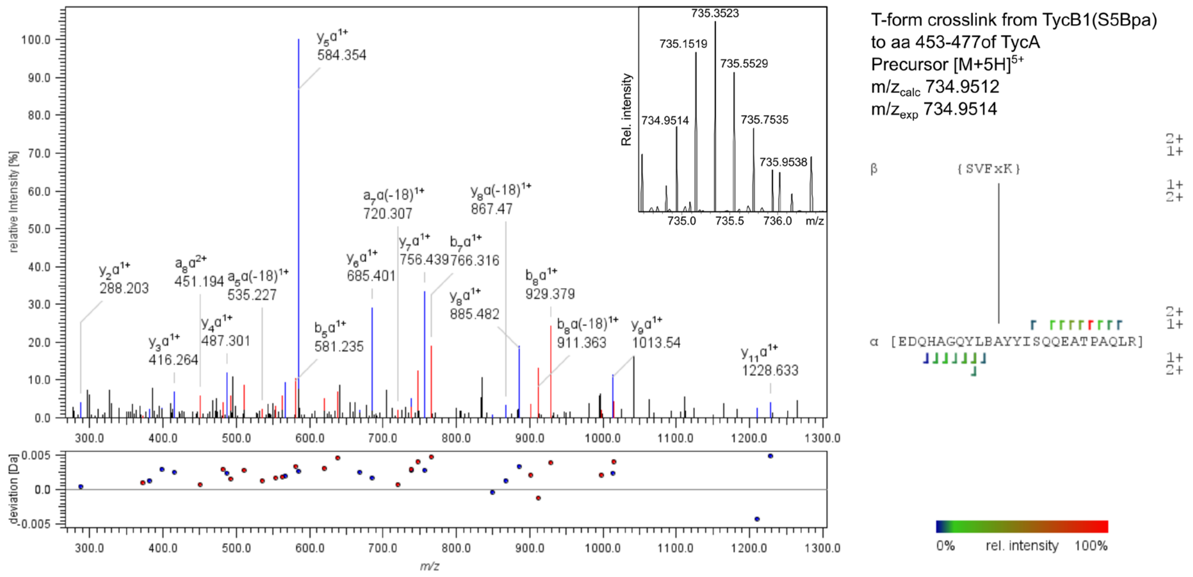

Figure S10 >>> continued

C

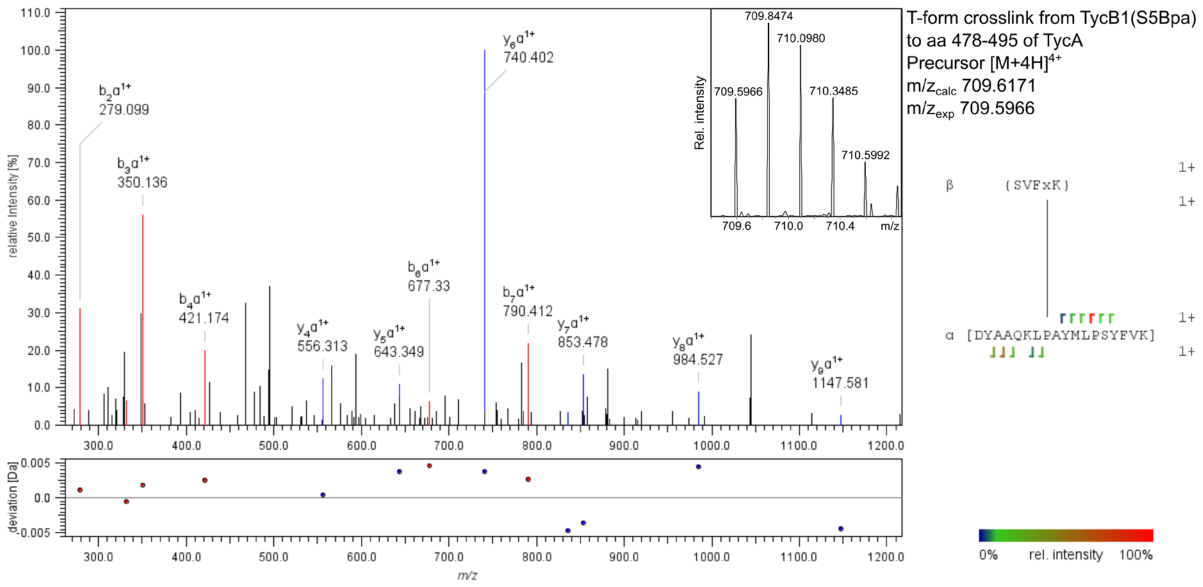

D

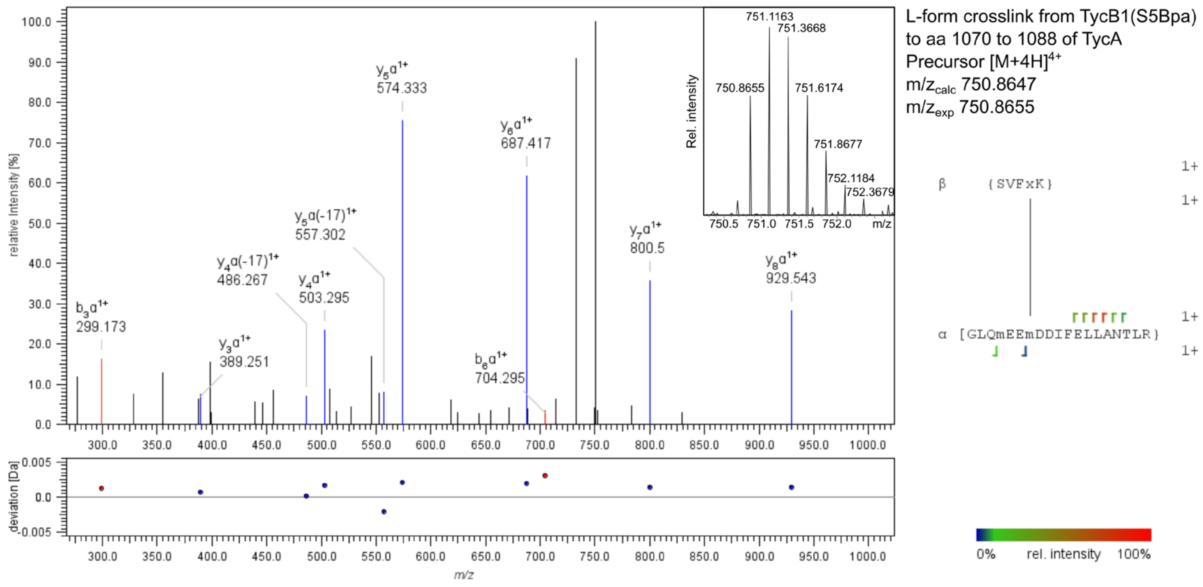

Figure S10. Mapping of photocrosslinks between TycB1 and TycA. A) SDS-PAGE gel of UV-irradiated Tyc module proteins and mixtures thereof. B-D) MSMS spectra of mapped crosslink peptides from tryptic digestion of TycA / TycB1 S5Bpa crosslink gel bands as indicated by arrows in A). General remarks: The corresponding precursors are shown in the upper-right insets of each MSMS spectrum. Crosslink peptides are schematically depicted next to each MSMS spectrum. In the amino acid sequences, “x” stands for the crosslinker Bpa, “B” denotes carbamidomethyl cysteine and “m” indicates oxidized methionine. Fragment ions flanking amino acids are indicated by half-square brackets. Due to the incomplete coverage of detectable fragment ions, there is a range of potential binding partners delimited by brackets rather than one amino acid position. Panel B) shows an identified mapped crosslink from the excised T-crosslink band into the peptide aa453-477 (EDQHAGQYLBAYYISQQEATPAQLR) of TycA, which corresponds to a part of the $A^C$ subdomain of TycA. Panel C) shows an identified crosslink from the excised T-crosslink band into the peptide aa478-495 (DYAAQKLPAYMLPSYFVK) of TycA, which also corresponds to a part of the $A^C$ subdomain of TycA. Panel D) shows an identified crosslink from the excised L-crosslink band into peptide aa1070-1089 (GLQmEEmDDIFELLANTLR) of TycA, which corresponds to the seam and helix region of the COM$^D$ domain of TycA.

### Supporting Tables

**Table S1**. Diffraction data collection and refinement statistics.

| | |
|---|---|
| **Wavelength (Å)** | 0.979510 |
| **Space group** | $P4_1$ |
| **Unit cell a, b, c (Å)**<br>**α, β,γ (°)** | 147.749, 147.749, 66.5<br>90 , 90, 90 |
| **Resolution range** | 49.42 – 3.29 (3.48-3.29) |
| **Total reflections** | 298,579 (45,532) |
| **Unique reflections** | 42,422 (6,884) |
| **Multiplicity** | 7.04 (6.61) |
| **Anomal Corr** | 53 (2) |
| **SigAno** | 1.31 (0.67) |
| **Completeness (%)** | 99.7 (98.3) |
| **I / sigma** | 9.55 (1.43) |
| **R-meas (%)** | 12.5 (116.4) |
| **R-pim (%)** | 4.71 (45.3) |
| **CC ½** | 0.996 (0.626) |
| **CC*** | 0.999 (0.877) |
| **Wilson B factor** | 129.44 |
| **Refinement** | 49.42 – 3.29 |
| **Space group** | P4 (1) |
| **Reflections in refinement** | 40297 |
| **Reflections in free set** | 2110 |
| **Rwork** | 0.2435 |
| **Rfree** | 0.3013 |
| **RMSD bonds** | 0.0013 |
| **RMSD angles** | 0.49 |
| **Ramachandran favoured (%)** | 97.28 |
| **Ramachandran allowed (%)** | 2.72 |
| **Ramachandran outliers (%)** | 0.0 |
| **Rotamer outliers** | 0.0 |
| **Clash score** | 2.76 |
| **B-factor (min/max/mean)** | 51.77/352.73/140.57 |
| **MolProbity score** | 1.20 |

Statistics for the last shell is given in parentheses. Refinement statistics were calculated with MolProbity and phenix_refine.

Table S2. List of purified recombinant proteins and the respective expression plasmids.

| No. | Protein | MW (kDa) | Encoding plasmid | Vector backbone |
|---|---|---|---|---|
| | BacC4-C5[E-COM-C]-His$_6$ | 106.366 | pAF24 | pET28a |
| 1 | BacC4-C5[A-PCP-E-COM-C-A-PCP-TE]-His$_6$ | 267.730 | pJD01 | pET28a |
| 2 | BacC4-C5[A-PCP-E-COM-C-A-PCP-TE(S6204A)]-His$_6$ | 267.714 | pJD16 | pET28a |
| 3 | BacC4-C5[A-PCP-E(H4755A)-COM-C-A-PCP-TE]-His$_6$ | 267.664 | pJD22 | pET28a |
| 4 | BacC4-C5[A-PCP-E(E4893A)-COM-C-A-PCP-TE]-His$_6$ | 267.672 | pJD23 | pET28a |
| 5 | BacC4-C5[A-PCP-E-COM(I5079R)-C-A-PCP-TE]-His$_6$ | 267.773 | pJD10 | pET28a |
| 6 | BacC4-C5[A-PCP-E-COM(I5079A)-C-A-PCP-TE]-His$_6$ | 267.688 | pJD11 | pET28a |
| 7 | BacC4-C5[A-PCP-E-COM(Δ5059-5069)-C-A-PCP-TE]-His$_6$ | 266.508 | pJD08 | pET28a |
| 8 | BacC4-C5[A-PCP-E-COM(Δ5062-5084)-C-A-PCP-TE]-His$_6$ | 265.107 | pJD07 | pET28a |
| 9 | BacC4-C5[A-PCP-E-COM(Ins5062(GGS)$_3$)-C-A-PCP-TE]-His$_6$ | 268.334 | pJD09 | pET28a |
| 10 | BacC4-C5[A-PCP-E-COM(Δ5059-5061)-C-A-PCP-TE]-His$_6$ | 267.372 | pJD14 | pET28a |
| 11 | BacC4-C5[A-PCP-E-COM(Ins5062G)-C-A-PCP-TE]-His$_6$ | 267.787 | pJD15 | pET28a |
| 12 | BacC4-C5[A-PCP-E-COM(Δ5059)-C-A-PCP-TE]-His$_6$ | 267.629 | pJD19 | pET28a |
| 13 | BacC4-C5[A-PCP-E-COM(Ins5068G)-C-A-PCP-TE]-His$_6$ | 267.787 | pJD21 | pET28a |
| 14 | BacC4-C5[A-PCP-E-COM(P5063A)-C-A-PCP-TE]-His$_6$ | 267.704 | pJD20 | pET28a |
| 15 | BacC4-C5[A-PCP-E-COM(Ins5086GGS)-C-A-PCP-TE]-His$_6$ | 267.931 | pJD13 | pET28a |
| 16 | SBP-TycA[A-PCP-E-COM$^D$] | 126.945 | pJR89 | pET28a |
| 17 | TycB1[COM$^A$-C-A-PCP]-His$_6$ | 119.696 (excluding Met1) | pJR95 (Ref[2]) | pTrc99a |
| 18 | SBP-TycA[A-PCP-E-COM$^D$(ΔHlx)] | 125.022 | pBH86 | pET28a |
| 19 | SBP-TycA[A-PCP-E-COM$^D$(ΔLinker)] | 125.698 | pBH81 | pET28a |
| 20 | SBP-TycA[A-PCP-E-COM$^D$(VER(GGS)$_3$TPSD)] | 127.548 | pBH82 | pET28a |
| 21 | SBP-TycA[A-PCP-E-COM$^D$(ΔLinker_sm)] | 126.560 | pBH83 | pET28a |
| 22 | SBP-TycA[A-PCP-E-COM$^D$(VERgTPSD)] | 127.002 | pBH88 | pET28a |
| 23 | SBP-TycA[A-PCP-E-COM$^D$(ΔV1059)] | 126.845 | pBH85 | pET28a |
| 24 | SBP-TycA[A-PCP-E-COM$^D$(P1063A)] | 126.919 | pBH89 | pET28a |
| 25 | TycB1[COM$^A$-C(S5X)-A-PCP]-His$_6$ (with S5Bpa) | 119.860 (excluding Met1) | pBH96 | pTrc99a |

### Amino Acid Sequences

Sequence sections differing from wild-type sequence are highlighted in yellow.

#### BacC4-C5[E-COM-C]-H$_6$

MGDKSASYETVEGEVLLTPIQQEYFSLNKTDRNHYNHAVMLYRKNGFDESIVKRVFKEIIKHHDALRTVFTEEDGKIIQYNRGPDKQLFDLFVYDVSSENDQPQKVYQLATELQ
QSIDIETGPLVKLALFKTNNGDHLLIIIHHLVVDGISWRILFEDLAIGYSQLANGEKVEFYPKTASYQAYARHIAEYAKSVKLLSEKQYWLKAIAEGVEFLDMNENAGAFKVEDSRT
FSTELEKEETKRLLRETNRAYHTEINDILITALLVAARDMNGQNQLRITLEGHGREQVADGIDISRTVGWFTSKYPVFIDLGQETDMSRTIKMVKEHLRNVPNKGIGYGILKYLTR
DSEIAKGAASPILFNYLGQLDEDINSGEFSSSHLSPGEAAGKGITREHPLEINAVVFRGKLAIQTTYNTRAYSEDVVRAFAQNYKEALKAVIRHCAEREETEKTPSDYGDKGISLD
QLEEIKLKYKGMEIEKIYPLANMQRGMLFHALEDKESQAYFEQMAINMKGLIDERLFAETFNDIMERHEILRASIEYEITDEPRNVIIKDRKINLDYHDLRKQSPAEREQVIQAYRK
ADREKGFRLNSEPLIRAALMRTEDDSYTFIWTNHHILLDGWSRGIIMGELFHMYHMKEARQKHRLEEARPYSDYIGWLQQQDKEAAKAYWRNYLSGFTEKSPISVLAGSSGHA
KYKRKEAVIEFPEQLTGRITELASRNNVTFHTVLQCIWGMLLARYNQTDDVVFGTVISGRDAQVTGIEKMVGLFINTVPTRIRLDKSQSFKELIKSVQEQALEGRTYHDMNLSEV
QSLSELKRELLDHILIFENYAVDQSAFETSGKRGAGFVFEEIHAEEQTNYGFNIVAVPGERLVIKLTYDGNIYHDHIIAGIKGHLQQVMEQVVQHEDQSLNDITVLSGSRSHHHHH
H

#### 1) BacC4-C5[A-PCP-E-COM-C-A-PCP-TE]-H$_6$

MDSDEEMNALLDQNQGKQADYPQDQTVHQLFEQQADKTPEQTAVVYADEKLTYRELNERANQLARLLRDKGADADQPVAIMIEPSLEMIISMLAVLKAGAAYVPIEPEQLAKR
TNEILSDSRAAILLVKGSVKENVAFAGEIVNVADGLIDAKVASNLSASGSADQNAYIIYTSGSTGKPKGVFVRHGNVVNYTTWFMKEAGLTENDKAMLVSSYAFDLGYTSIFSAL
LSGSELHIARKECYTNAHRALKYIKENGITYIKLTPSLFNIFVNDPGFSAEKPCATLRLVVLGGEMINTRDVETFYNQYPDHVVMNHYGPTETTIGSVFKVIDPEHLDSFKECPVIG
TPIHNTNAYVLDENMKLLPEGVYGELCIAGAGVTGGYVNRPDETKEKFIENPFAPHTKMYRTGDLARRLSDGNIELAGRIDTQVKVRGYRIEPEEIKNRLLAHDDIKEAFIAARED
HKGAKQLCAYFTADAELPFEDIRTYLMHELPEYMIPSSFVQIEKMPLSANGKIDTAALPEPQPGKETEYEPPRNETEEKLVQIWEEVLGIDKIGITHHFFAAGGDSIKALQMISRLS
REGLSLEMKDLFANPQIKSLSRYVKAESDKSASYETVEGEVLLTPIQQEYFSLNKTDRNHYNHAVMLYRKNGFDESIVKRVFKEIIKHHDALRTVFTEEDGKIIQYNRGPDKQLF
DLFVYDVSSENDQPQKVYQLATELQQSIDIETGPLVKLALFKTNNGDHLLIIIHHLVVDGISWRILFEDLAIGYSQLANGEKVEFYPKTASYQAYARHIAEYAKSVKLLSEKQYWLK
AIAEGVEFLDMNENAGAFKVEDSRTFSTELEKEETKRLLRETNRAYHTEINDILITALLVAARDMNGQNQLRITLEGHGREQVADGIDISRTVGWFTSKYPVFIDLGQETDMSRTI
KMVKEHLRNVPNKGIGYGILKYLTRDSEIAKGAASPILFNYLGQLDEDINSGEFSSSHLSPGEAAGKGITREHPLEINAVVFRGKLAIQTTYNTRAYSEDVVRAFAQNYKEALKAV
IRHCAEREETEKTPSDYGDKGISLDQLEEIKLKYKGMEIEKIYPLANMQRGMLFHALEDKESQAYFEQMAINMKGLIDERLFAETFNDIMERHEILRASIEYEITDEPRNVIIKDRKI
NLDYHDLRKQSPAEREQVIQAYRKADREKGFRLNSEPLIRAALMRTEDDSYTFIWTNHHILLDGWSRGIIMGELFHMYHMKEARQKHRLEEARPYSDYIGWLQQQDKEAAKAY
WRNYLSGFTEKSPISVLAGSSGHAKYKRKEAVIEFPEQLTGRITELASRNNVTFHTVLQCIWGMLLARYNQTDDVVFGTVISGRDAQVTGIEKMVGLFINTVPTRIRLDKSQSFK
ELIKSVQEQALEGRTYHDMNLSEVQSLSELKRELLDHILIFENYAVDQSAFETSGKRGAGFVFEEIHAEEQTNYGFNIVAVPGERLVIKLTYDGNIYHDHIIAGIKGHLQQVMEQV
VQHEDQSLNDITVLSEAERNRLLYEWNDTKAEYPNQTIHRLFEEQAEKTPELAAVVSGNDKLTYRELNEKSNQLARYLRDKGVKADTIVAIMAERSPEMVVGIMGILKAGGAYL
PIDPDYPEERIKYMLEDSGAAIILADHKQDLGTLHQEAVELTGDFSSYPADNLEPAGNADSLAYIIYTSGSTGKPKGVMIRQRGLVNYITWADRVYVQGEQLDFALYSSIAFDLTV
TSIFTPLISGNRVIVYRHSEDGEPLIRKVFRDQKAGIVKLTPSHLSLVKDMDASGSSIKRLIVGGEDLKTELAKEITERFHHNIEIYNEYGPTETVVGCMIYQYDAGWDRQVSVPIG
KPASNVQLYILDERQEVQPVGIAGELYISGDGVAKGYLNKPELTSEKFLPNPFLPGERMYRTGDLAKMRPDGHIEYLGRIDHQVKIRGYRIELGEIEHQLLRHSDIKEAAVAAKTD
QNNDQVLCAYVVSERDITQKDIKTFLAKELPEYMVPSYLLKLDELPLTPNGKVDLKALPEPDRSAGALLEYEPPRHELEEKMAAIWEDILNIEQIGINANIFDIGANSLNVMSFVSR
LYAELGFRVPFKDIFSKPTIKELSDFLKHAQDLLKDYTDDCMQLTRAEEGGKNLFCFPPAASMGIAYMGLAKHLKQHSVYSFNFIPSANRIRKYADIIKNIQGEGPYTLIGYSSGGI
LAFDVAKELNRQGYEVEDLIIIDSKYRTKAEKHQFTEEEYREEISKTFELEKYRDVEKLLSDYLVDLVMKSYVYIQNTVTTGAIDGHISYIKSSDNQRGENMMMWEKATSKTFTVV
QGAGTHMQMISKSHPDILERNARLIHDIINKTVKIGSRSHHHHHH

#### 2) BacC4-C5[A-PCP-E-COM-C-A-PCP-TE(S6204A)]-H$_6$

MDSDEEMNALLDQNQGKQADYPQDQTVHQLFEQQADKTPEQTAVVYADEKLTYRELNERANQLARLLRDKGADADQPVAIMIEPSLEMIISMLAVLKAGAAYVPIEPEQLAKR
TNEILSDSRAAILLVKGSVKENVAFAGEIVNVADGLIDAKVASNLSASGSADQNAYIIYTSGSTGKPKGVFVRHGNVVNYTTWFMKEAGLTENDKAMLVSSYAFDLGYTSIFSAL
LSGSELHIARKECYTNAHRALKYIKENGITYIKLTPSLFNIFVNDPGFSAEKPCATLRLVVLGGEMINTRDVETFYNQYPDHVVMNHYGPTETTIGSVFKVIDPEHLDSFKECPVIG
TPIHNTNAYVLDENMKLLPEGVYGELCIAGAGVTGGYVNRPDETKEKFIENPFAPHTKMYRTGDLARRLSDGNIELAGRIDTQVKVRGYRIEPEEIKNRLLAHDDIKEAFIAARED
HKGAKQLCAYFTADAELPFEDIRTYLMHELPEYMIPSSFVQIEKMPLSANGKIDTAALPEPQPGKETEYEPPRNETEEKLVQIWEEVLGIDKIGITHHFFAAGGDSIKALQMISRLS
REGLSLEMKDLFANPQIKSLSRYVKAESDKSASYETVEGEVLLTPIQQEYFSLNKTDRNHYNHAVMLYRKNGFDESIVKRVFKEIIKHHDALRTVFTEEDGKIIQYNRGPDKQLF
DLFVYDVSSENDQPQKVYQLATELQQSIDIETGPLVKLALFKTNNGDHLLIIIHHLVVDGISWRILFEDLAIGYSQLANGEKVEFYPKTASYQAYARHIAEYAKSVKLLSEKQYWLK
AIAEGVEFLDMNENAGAFKVEDSRTFSTELEKEETKRLLRETNRAYHTEINDILITALLVAARDMNGQNQLRITLEGHGREQVADGIDISRTVGWFTSKYPVFIDLGQETDMSRTI
KMVKEHLRNVPNKGIGYGILKYLTRDSEIAKGAASPILFNYLGQLDEDINSGEFSSSHLSPGEAAGKGITREHPLEINAVVFRGKLAIQTTYNTRAYSEDVVRAFAQNYKEALKAV
IRHCAEREETEKTPSDYGDKGISLDQLEEIKLKYKGMEIEKIYPLANMQRGMLFHALEDKESQAYFEQMAINMKGLIDERLFAETFNDIMERHEILRASIEYEITDEPRNVIIKDRKI
NLDYHDLRKQSPAEREQVIQAYRKADREKGFRLNSEPLIRAALMRTEDDSYTFIWTNHHILLDGWSRGIIMGELFHMYHMKEARQKHRLEEARPYSDYIGWLQQQDKEAAKAY
WRNYLSGFTEKSPISVLAGSSGHAKYKRKEAVIEFPEQLTGRITELASRNNVTFHTVLQCIWGMLLARYNQTDDVVFGTVISGRDAQVTGIEKMVGLFINTVPTRIRLDKSQSFK
ELIKSVQEQALEGRTYHDMNLSEVQSLSELKRELLDHILIFENYAVDQSAFETSGKRGAGFVFEEIHAEEQTNYGFNIVAVPGERLVIKLTYDGNIYHDHIIAGIKGHLQQVMEQV
VQHEDQSLNDITVLSEAERNRLLYEWNDTKAEYPNQTIHRLFEEQAEKTPELAAVVSGNDKLTYRELNEKSNQLARYLRDKGVKADTIVAIMAERSPEMVVGIMGILKAGGAYL
PIDPDYPEERIKYMLEDSGAAIILADHKQDLGTLHQEAVELTGDFSSYPADNLEPAGNADSLAYIIYTSGSTGKPKGVMIRQRGLVNYITWADRVYVQGEQLDFALYSSIAFDLTV
TSIFTPLISGNRVIVYRHSEDGEPLIRKVFRDQKAGIVKLTPSHLSLVKDMDASGSSIKRLIVGGEDLKTELAKEITERFHHNIEIYNEYGPTETVVGCMIYQYDAGWDRQVSVPIG
KPASNVQLYILDERQEVQPVGIAGELYISGDGVAKGYLNKPELTSEKFLPNPFLPGERMYRTGDLAKMRPDGHIEYLGRIDHQVKIRGYRIELGEIEHQLLRHSDIKEAAVAAKTD
QNNDQVLCAYVVSERDITQKDIKTFLAKELPEYMVPSYLLKLDELPLTPNGKVDLKALPEPDRSAGALLEYEPPRHELEEKMAAIWEDILNIEQIGINANIFDIGANSLNVMSFVSR
LYAELGFRVPFKDIFSKPTIKELSDFLKHAQDLLKDYTDDCMQLTRAEEGGKNLFCFPPAASMGIAYMGLAKHLKQHSVYSFNFIPSANRIRKYADIIKNIQGEGPYTLIGYASGGI
LAFDVAKELNRQGYEVEDLIIIDSKYRTKAEKHQFTEEEYREEISKTFELEKYRDVEKLLSDYLVDLVMKSYVYIQNTVTTGAIDGHISYIKSSDNQRGENMMMWEKATSKTFTVV
QGAGTHMQMISKSHPDILERNARLIHDIINKTVKIGSRSHHHHHH

#### 3) BacC4-C5[A-PCP-E(H4755A)-COM-C-A-PCP-TE]-H$_6$

MDSDEEMNALLDQNQGKQADYPQDQTVHQLFEQQADKTPEQTAVVYADEKLTYRELNERANQLARLLRDKGADADQPVAIMIEPSLEMIISMLAVLKAGAAYVPIEPEQLAKR
TNEILSDSRAAILLVKGSVKENVAFAGEIVNVADGLIDAKVASNLSASGSADQNAYIIYTSGSTGKPKGVFVRHGNVVNYTTWFMKEAGLTENDKAMLVSSYAFDLGYTSIFSAL
LSGSELHIARKECYTNAHRALKYIKENGITYIKLTPSLFNIFVNDPGFSAEKPCATLRLVVLGGEMINTRDVETFYNQYPDHVVMNHYGPTETTIGSVFKVIDPEHLDSFKECPVIG
TPIHNTNAYVLDENMKLLPEGVYGELCIAGAGVTGGYVNRPDETKEKFIENPFAPHTKMYRTGDLARRLSDGNIELAGRIDTQVKVRGYRIEPEEIKNRLLAHDDIKEAFIAARED
HKGAKQLCAYFTADAELPFEDIRTYLMHELPEYMIPSSFVQIEKMPLSANGKIDTAALPEPQPGKETEYEPPRNETEEKLVQIWEEVLGIDKIGITHHFFAAGGDSIKALQMISRLS
REGLSLEMKDLFANPQIKSLSRYVKAESDKSASYETVEGEVLLTPIQQEYFSLNKTDRNHYNHAVMLYRKNGFDESIVKRVFKEIIKHHDALRTVFTEEDGKIIQYNRGPDKQLF
DLFVYDVSSENDQPQKVYQLATELQQSIDIETGPLVKLALFKTNNGDHLLIIIHALVVDGISWRILFEDLAIGYSQLANGEKVEFYPKTASYQAYARHIAEYAKSVKLLSEKQYWLK
AIAEGVEFLDMNENAGAFKVEDSRTFSTELEKEETKRLLRETNRAYHTEINDILITALLVAARDMNGQNQLRITLEGHGREQVADGIDISRTVGWFTSKYPVFIDLGQETDMSRTI
KMVKEHLRNVPNKGIGYGILKYLTRDSEIAKGAASPILFNYLGQLDEDINSGEFSSSHLSPGEAAGKGITREHPLEINAVVFRGKLAIQTTYNTRAYSEDVVRAFAQNYKEALKAV
IRHCAEREETEKTPSDYGDKGISLDQLEEIKLKYKGMEIEKIYPLANMQRGMLFHALEDKESQAYFEQMAINMKGLIDERLFAETFNDIMERHEILRASIEYEITDEPRNVIIKDRKI
NLDYHDLRKQSPAEREQVIQAYRKADREKGFRLNSEPLIRAALMRTEDDSYTFIWTNHHILLDGWSRGIIMGELFHMYHMKEARQKHRLEEARPYSDYIGWLQQQDKEAAKAY
WRNYLSGFTEKSPISVLAGSSGHAKYKRKEAVIEFPEQLTGRITELASRNNVTFHTVLQCIWGMLLARYNQTDDVVFGTVISGRDAQVTGIEKMVGLFINTVPTRIRLDKSQSFK
ELIKSVQEQALEGRTYHDMNLSEVQSLSELKRELLDHILIFENYAVDQSAFETSGKRGAGFVFEEIHAEEQTNYGFNIVAVPGERLVIKLTYDGNIYHDHIIAGIKGHLQQVMEQV
VQHEDQSLNDITVLSEAERNRLLYEWNDTKAEYPNQTIHRLFEEQAEKTPELAAVVSGNDKLTYRELNEKSNQLARYLRDKGVKADTIVAIMAERSPEMVVGIMGILKAGGAYL
PIDPDYPEERIKYMLEDSGAAIILADHKQDLGTLHQEAVELTGDFSSYPADNLEPAGNADSLAYIIYTSGSTGKPKGVMIRQRGLVNYITWADRVYVQGEQLDFALYSSIAFDLTV
TSIFTPLISGNRVIVYRHSEDGEPLIRKVFRDQKAGIVKLTPSHLSLVKDMDASGSSIKRLIVGGEDLKTELAKEITERFHHNIEIYNEYGPTETVVGCMIYQYDAGWDRQVSVPIG
KPASNVQLYILDERQEVQPVGIAGELYISGDGVAKGYLNKPELTSEKFLPNPFLPGERMYRTGDLAKMRPDGHIEYLGRIDHQVKIRGYRIELGEIEHQLLRHSDIKEAAVAAKTD
QNNDQVLCAYVVSERDITQKDIKTFLAKELPEYMVPSYLLKLDELPLTPNGKVDLKALPEPDRSAGALLEYEPPRHELEEKMAAIWEDILNIEQIGINANIFDIGANSLNVMSFVSR
LYAELGFRVPFKDIFSKPTIKELSDFLKHAQDLLKDYTDDCMQLTRAEEGGKNLFCFPPAASMGIAYMGLAKHLKQHSVYSFNFIPSANRIRKYADIIKNIQGEGPYTLIGYSSGGI
LAFDVAKELNRQGYEVEDLIIIDSKYRTKAEKHQFTEEEYREEISKTFELEKYRDVEKLLSDYLVDLVMKSYVYIQNTVTTGAIDGHISYIKSSDNQRGENMMMWEKATSKTFTVV
QGAGTHMQMISKSHPDILERNARLIHDIINKTVKIGSRSHHHHHH

#### 4) BacC4-C5[A-PCP-E(E4893A)-COM-C-A-PCP-TE]-H$_6$

MDSDEEMNALLDQNQGKQADYPQDQTVHQLFEQQADKTPEQTAVVYADEKLTYRELNERANQLARLLRDKGADADQPVAIMIEPSLEMIISMLAVLKAGAAYVPIEPEQLAKR
TNEILSDSRAAILLVKGSVKENVAFAGEIVNVADGLIDAKVASNLSASGSADQNAYIIYTSGSTGKPKGVFVRHGNVVNYTTWFMKEAGLTENDKAMLVSSYAFDLGYTSIFSAL

LSGSELHIARKECYTNAHRALKYIKENGITYIKLTPSLFNIFVNDPGFSAEKPCATLRLVVLGGEMINTRDVETFYNQYPDHVVMNHYGPTETTIGSVFKVIDPEHLDSFKECPVIG
TPIHNTNAYVLDENMKLLPEGVYGELCIAGAGVTGGYVNRPDETKEKFIENPFAPHTKMYRTGDLARRLSDGNIELAGRIDTQVKVRGYRIEPEEIKNRLLAHDDIKEAFIAARED
HKGAKQLCAYFTADAELPFEDIRTYLMHELPEYMIPSSFVQIEKMPLSANGKIDTAALPEPQPGKETEYEPPRNETEEKLVQIWEEVLGIDKIGITHHFFAAGGDSIKALQMISRLS
REGLSLEMKDLFANPQIKSLSRYVKAESDKSASYETVEGEVLLTPIQQEYFSLNKTDRNHYNHAVMLYRKNGFDESIVKRVFKEIIKHHDALRTVFTEEDGKIIQYNRGPDKQLF
DLFVYDVSSENDQPQKVYQLATELQQSIDIETGPLVKLALFKTNNGDHLLIIIHHLVVDGISWRILFEDLAIGYSQLANGEKVEFYPKTASYQAYARHIAEYAKSVKLLSEKQYWLK
AIAEGVEFLDMNENAGAFKVEDSRTFSTELEKEETKRLLRETNRAYHTEINDILITALLVAARDMNGQNQLRITLAGHGREQVADGIDISRTVGWFTSKYPVFIDLGQETDMSRTI
KMVKEHLRNVPNKGIGYGILKYLTRDSEIAKGAASPILFNYLGQLDEDINSGEFSSSHLSPGEAAGKGITREHPLEINAVVFRGKLAIQTTYNTRAYSEDVVRAFAQNYKEALKAV
IRHCAEREETEKTPSDYGDKGISLDQLEEIKLKYKGMEIEKIYPLANMQRGMLFHALEDKESQAYFEQMAINMKGLIDERLFAETFNDIMERHEILRASIEYEITDEPRNVIIKDRKI
NLDYHDLRKQSPAEREQVIQAYRKADREKGFRLNSEPLIRAALMRTEDDSYTFIWTNHHILLDGWSRGIIMGELFHMYHMKEARQKHRLEEARPYSDYIGWLQQQDKEAAKAY
WRNYLSGFTEKSPISVLAGSSGHAKYKRKEAVIEFPEQLTGRITELASRNNVTFHTVLQCIWGMLLARYNQTDDVVFGTVISGRDAQVTGIEKMVGLFINTVPTRIRLDKSQSFK
ELIKSVQEQALEGRTYHDMNLSEVQSLSELKRELLDHILIFENYAVDQSAFETSGKRGAGFVFEEIHAEEQTNYGFNIVAVPGERLVIKLTYDGNIYHDHIIAGIKGHLQQVMEQV
VQHEDQSLNDITVLSEAERNRLLYEWNDTKAEYPNQTIHRLFEEQAEKTPELAAVVSGNDKLTYRELNEKSNQLARYLRDKGVKADTIVAIMAERSPEMVVGIMGILKAGGAYL
PIDPDYPEERIKYMLEDSGAAIILADHKQDLGTLHQEAVELTGDFSSYPADNLEPAGNADSLAYIIYTSGSTGKPKGVMIRQRGLVNYITWADRVYVQGEQLDFALYSSIAFDLTV
TSIFTPLISGNRVIVYRHSEDGEPLIRKVFRDQKAGIVKLTPSHLSLVKDMDASGSSIKRLIVGGEDLKTELAKEITERFHHNIEIYNEYGPTETVVGCMIYQYDAGWDRQVSVPIG
KPASNVQLYILDERQEVQPVGIAGELYISGDGVAKGYLNKPELTSEKFLPNPFLPGERMYRTGDLAKMRPDGHIEYLGRIDHQVKIRGYRIELGEIEHQLLRHSDIKEAAVAAKTD
QNNDQVLCAYVVSERDITQKDIKTFLAKELPEYMVPSYLLKLDELPLTPNGKVDLKALPEPDRSAGALLEYEPPRHELEEKMAAIWEDILNIEQIGINANIFDIGANSLNVMSFVSR
LYAELGFRVPFKDIFSKPTIKELSDFLKHAQDLLKDYTDDCMQLTRAEEGGKNLFCFPPAASMGIAYMGLAKHLKQHSVYSFNFIPSANRIRKYADIIKNIQGEGPYTLIGYSSGGI
LAFDVAKELNRQGYEVEDLIIIDSKYRTKAEKHQFTEEEYREEISKTFELEKYRDVEKLLSDYLVDLVMKSYVYIQNTVTTGAIDGHISYIKSSDNQRGENMMMWEKATSKTFTVV
QGAGTHMQMISKSHPDILERNARLIHDIINKTVKIGSRSHHHHHH

#### 5) BacC4-C5[A-PCP-E-COM(I5079R)-C-A-PCP-TE]-H$_6$

MDSDEEMNALLDQNQGKQADYPQDQTVHQLFEQQADKTPEQTAVVYADEKLTYRELNERANQLARLLRDKGADADQPVAIMIEPSLEMIISMLAVLKAGAAYVPIEPEQLAKR
TNEILSDSRAAILLVKGSVKENVAFAGEIVNVADGLIDAKVASNLSASGSADQNAYIIYTSGSTGKPKGVFVRHGNVVNYTTWFMKEAGLTENDKAMLVSSYAFDLGYTSIFSAL
LSGSELHIARKECYTNAHRALKYIKENGITYIKLTPSLFNIFVNDPGFSAEKPCATLRLVVLGGEMINTRDVETFYNQYPDHVVMNHYGPTETTIGSVFKVIDPEHLDSFKECPVIG
TPIHNTNAYVLDENMKLLPEGVYGELCIAGAGVTGGYVNRPDETKEKFIENPFAPHTKMYRTGDLARRLSDGNIELAGRIDTQVKVRGYRIEPEEIKNRLLAHDDIKEAFIAARED
HKGAKQLCAYFTADAELPFEDIRTYLMHELPEYMIPSSFVQIEKMPLSANGKIDTAALPEPQPGKETEYEPPRNETEEKLVQIWEEVLGIDKIGITHHFFAAGGDSIKALQMISRLS
REGLSLEMKDLFANPQIKSLSRYVKAESDKSASYETVEGEVLLTPIQQEYFSLNKTDRNHYNHAVMLYRKNGFDESIVKRVFKEIIKHHDALRTVFTEEDGKIIQYNRGPDKQLF
DLFVYDVSSENDQPQKVYQLATELQQSIDIETGPLVKLALFKTNNGDHLLIIIHHLVVDGISWRILFEDLAIGYSQLANGEKVEFYPKTASYQAYARHIAEYAKSVKLLSEKQYWLK
AIAEGVEFLDMNENAGAFKVEDSRTFSTELEKEETKRLLRETNRAYHTEINDILITALLVAARDMNGQNQLRITLEGHGREQVADGIDISRTVGWFTSKYPVFIDLGQETDMSRTI
KMVKEHLRNVPNKGIGYGILKYLTRDSEIAKGAASPILFNYLGQLDEDINSGEFSSSHLSPGEAAGKGITREHPLEINAVVFRGKLAIQTTYNTRAYSEDVVRAFAQNYKEALKAV
IRHCAEREETEKTPSDYGDKGISLDQLEERKLKYKGMEIEKIYPLANMQRGMLFHALEDKESQAYFEQMAINMKGLIDERLFAETFNDIMERHEILRASIEYEITDEPRNVIIKDRKI
NLDYHDLRKQSPAEREQVIQAYRKADREKGFRLNSEPLIRAALMRTEDDSYTFIWTNHHILLDGWSRGIIMGELFHMYHMKEARQKHRLEEARPYSDYIGWLQQQDKEAAKAY
WRNYLSGFTEKSPISVLAGSSGHAKYKRKEAVIEFPEQLTGRITELASRNNVTFHTVLQCIWGMLLARYNQTDDVVFGTVISGRDAQVTGIEKMVGLFINTVPTRIRLDKSQSFK
ELIKSVQEQALEGRTYHDMNLSEVQSLSELKRELLDHILIFENYAVDQSAFETSGKRGAGFVFEEIHAEEQTNYGFNIVAVPGERLVIKLTYDGNIYHDHIIAGIKGHLQQVMEQV
VQHEDQSLNDITVLSEAERNRLLYEWNDTKAEYPNQTIHRLFEEQAEKTPELAAVVSGNDKLTYRELNEKSNQLARYLRDKGVKADTIVAIMAERSPEMVVGIMGILKAGGAYL
PIDPDYPEERIKYMLEDSGAAIILADHKQDLGTLHQEAVELTGDFSSYPADNLEPAGNADSLAYIIYTSGSTGKPKGVMIRQRGLVNYITWADRVYVQGEQLDFALYSSIAFDLTV
TSIFTPLISGNRVIVYRHSEDGEPLIRKVFRDQKAGIVKLTPSHLSLVKDMDASGSSIKRLIVGGEDLKTELAKEITERFHHNIEIYNEYGPTETVVGCMIYQYDAGWDRQVSVPIG
KPASNVQLYILDERQEVQPVGIAGELYISGDGVAKGYLNKPELTSEKFLPNPFLPGERMYRTGDLAKMRPDGHIEYLGRIDHQVKIRGYRIELGEIEHQLLRHSDIKEAAVAAKTD
QNNDQVLCAYVVSERDITQKDIKTFLAKELPEYMVPSYLLKLDELPLTPNGKVDLKALPEPDRSAGALLEYEPPRHELEEKMAAIWEDILNIEQIGINANIFDIGANSLNVMSFVSR
LYAELGFRVPFKDIFSKPTIKELSDFLKHAQDLLKDYTDDCMQLTRAEEGGKNLFCFPPAASMGIAYMGLAKHLKQHSVYSFNFIPSANRIRKYADIIKNIQGEGPYTLIGYSSGGI
LAFDVAKELNRQGYEVEDLIIIDSKYRTKAEKHQFTEEEYREEISKTFELEKYRDVEKLLSDYLVDLVMKSYVYIQNTVTTGAIDGHISYIKSSDNQRGENMMMWEKATSKTFTVV
QGAGTHMQMISKSHPDILERNARLIHDIINKTVKIGSRSHHHHHH

#### 6) BacC4-C5[A-PCP-E-COM(I5079A)-C-A-PCP-TE]-H$_6$

MDSDEEMNALLDQNQGKQADYPQDQTVHQLFEQQADKTPEQTAVVYADEKLTYRELNERANQLARLLRDKGADADQPVAIMIEPSLEMIISMLAVLKAGAAYVPIEPEQLAKR
TNEILSDSRAAILLVKGSVKENVAFAGEIVNVADGLIDAKVASNLSASGSADQNAYIIYTSGSTGKPKGVFVRHGNVVNYTTWFMKEAGLTENDKAMLVSSYAFDLGYTSIFSAL
LSGSELHIARKECYTNAHRALKYIKENGITYIKLTPSLFNIFVNDPGFSAEKPCATLRLVVLGGEMINTRDVETFYNQYPDHVVMNHYGPTETTIGSVFKVIDPEHLDSFKECPVIG
TPIHNTNAYVLDENMKLLPEGVYGELCIAGAGVTGGYVNRPDETKEKFIENPFAPHTKMYRTGDLARRLSDGNIELAGRIDTQVKVRGYRIEPEEIKNRLLAHDDIKEAFIAARED
HKGAKQLCAYFTADAELPFEDIRTYLMHELPEYMIPSSFVQIEKMPLSANGKIDTAALPEPQPGKETEYEPPRNETEEKLVQIWEEVLGIDKIGITHHFFAAGGDSIKALQMISRLS
REGLSLEMKDLFANPQIKSLSRYVKAESDKSASYETVEGEVLLTPIQQEYFSLNKTDRNHYNHAVMLYRKNGFDESIVKRVFKEIIKHHDALRTVFTEEDGKIIQYNRGPDKQLF
DLFVYDVSSENDQPQKVYQLATELQQSIDIETGPLVKLALFKTNNGDHLLIIIHHLVVDGISWRILFEDLAIGYSQLANGEKVEFYPKTASYQAYARHIAEYAKSVKLLSEKQYWLK
AIAEGVEFLDMNENAGAFKVEDSRTFSTELEKEETKRLLRETNRAYHTEINDILITALLVAARDMNGQNQLRITLEGHGREQVADGIDISRTVGWFTSKYPVFIDLGQETDMSRTI
KMVKEHLRNVPNKGIGYGILKYLTRDSEIAKGAASPILFNYLGQLDEDINSGEFSSSHLSPGEAAGKGITREHPLEINAVVFRGKLAIQTTYNTRAYSEDVVRAFAQNYKEALKAV
IRHCAEREETEKTPSDYGDKGISLDQLEEAKLKYKGMEIEKIYPLANMQRGMLFHALEDKESQAYFEQMAINMKGLIDERLFAETFNDIMERHEILRASIEYEITDEPRNVIIKDRKI
NLDYHDLRKQSPAEREQVIQAYRKADREKGFRLNSEPLIRAALMRTEDDSYTFIWTNHHILLDGWSRGIIMGELFHMYHMKEARQKHRLEEARPYSDYIGWLQQQDKEAAKAY
WRNYLSGFTEKSPISVLAGSSGHAKYKRKEAVIEFPEQLTGRITELASRNNVTFHTVLQCIWGMLLARYNQTDDVVFGTVISGRDAQVTGIEKMVGLFINTVPTRIRLDKSQSFK
ELIKSVQEQALEGRTYHDMNLSEVQSLSELKRELLDHILIFENYAVDQSAFETSGKRGAGFVFEEIHAEEQTNYGFNIVAVPGERLVIKLTYDGNIYHDHIIAGIKGHLQQVMEQV
VQHEDQSLNDITVLSEAERNRLLYEWNDTKAEYPNQTIHRLFEEQAEKTPELAAVVSGNDKLTYRELNEKSNQLARYLRDKGVKADTIVAIMAERSPEMVVGIMGILKAGGAYL
PIDPDYPEERIKYMLEDSGAAIILADHKQDLGTLHQEAVELTGDFSSYPADNLEPAGNADSLAYIIYTSGSTGKPKGVMIRQRGLVNYITWADRVYVQGEQLDFALYSSIAFDLTV
TSIFTPLISGNRVIVYRHSEDGEPLIRKVFRDQKAGIVKLTPSHLSLVKDMDASGSSIKRLIVGGEDLKTELAKEITERFHHNIEIYNEYGPTETVVGCMIYQYDAGWDRQVSVPIG
KPASNVQLYILDERQEVQPVGIAGELYISGDGVAKGYLNKPELTSEKFLPNPFLPGERMYRTGDLAKMRPDGHIEYLGRIDHQVKIRGYRIELGEIEHQLLRHSDIKEAAVAAKTD
QNNDQVLCAYVVSERDITQKDIKTFLAKELPEYMVPSYLLKLDELPLTPNGKVDLKALPEPDRSAGALLEYEPPRHELEEKMAAIWEDILNIEQIGINANIFDIGANSLNVMSFVSR
LYAELGFRVPFKDIFSKPTIKELSDFLKHAQDLLKDYTDDCMQLTRAEEGGKNLFCFPPAASMGIAYMGLAKHLKQHSVYSFNFIPSANRIRKYADIIKNIQGEGPYTLIGYSSGGI
LAFDVAKELNRQGYEVEDLIIIDSKYRTKAEKHQFTEEEYREEISKTFELEKYRDVEKLLSDYLVDLVMKSYVYIQNTVTTGAIDGHISYIKSSDNQRGENMMMWEKATSKTFTVV
QGAGTHMQMISKSHPDILERNARLIHDIINKTVKIGSRSHHHHHH

#### 7) BacC4-C5[A-PCP-E-COM(Δ5059-5069)-C-A-PCP-TE]-H$_6$

MDSDEEMNALLDQNQGKQADYPQDQTVHQLFEQQADKTPEQTAVVYADEKLTYRELNERANQLARLLRDKGADADQPVAIMIEPSLEMIISMLAVLKAGAAYVPIEPEQLAKR
TNEILSDSRAAILLVKGSVKENVAFAGEIVNVADGLIDAKVASNLSASGSADQNAYIIYTSGSTGKPKGVFVRHGNVVNYTTWFMKEAGLTENDKAMLVSSYAFDLGYTSIFSAL
LSGSELHIARKECYTNAHRALKYIKENGITYIKLTPSLFNIFVNDPGFSAEKPCATLRLVVLGGEMINTRDVETFYNQYPDHVVMNHYGPTETTIGSVFKVIDPEHLDSFKECPVIG
TPIHNTNAYVLDENMKLLPEGVYGELCIAGAGVTGGYVNRPDETKEKFIENPFAPHTKMYRTGDLARRLSDGNIELAGRIDTQVKVRGYRIEPEEIKNRLLAHDDIKEAFIAARED
HKGAKQLCAYFTADAELPFEDIRTYLMHELPEYMIPSSFVQIEKMPLSANGKIDTAALPEPQPGKETEYEPPRNETEEKLVQIWEEVLGIDKIGITHHFFAAGGDSIKALQMISRLS
REGLSLEMKDLFANPQIKSLSRYVKAESDKSASYETVEGEVLLTPIQQEYFSLNKTDRNHYNHAVMLYRKNGFDESIVKRVFKEIIKHHDALRTVFTEEDGKIIQYNRGPDKQLF
DLFVYDVSSENDQPQKVYQLATELQQSIDIETGPLVKLALFKTNNGDHLLIIIHHLVVDGISWRILFEDLAIGYSQLANGEKVEFYPKTASYQAYARHIAEYAKSVKLLSEKQYWLK
AIAEGVEFLDMNENAGAFKVEDSRTFSTELEKEETKRLLRETNRAYHTEINDILITALLVAARDMNGQNQLRITLEGHGREQVADGIDISRTVGWFTSKYPVFIDLGQETDMSRTI
KMVKEHLRNVPNKGIGYGILKYLTRDSEIAKGAASPILFNYLGQLDEDINSGEFSSSHLSPGEAAGKGITREHPLEINAVVFRGKLAIQTTYNTRAYSEDVVRAFAQNYKEALKAV
IRHCAEREEGISLDQLEEIKLKYKGMEIEKIYPLANMQRGMLFHALEDKESQAYFEQMAINMKGLIDERLFAETFNDIMERHEILRASIEYEITDEPRNVIIKDRKINLDYHDLRKQS
PAEREQVIQAYRKADREKGFRLNSEPLIRAALMRTEDDSYTFIWTNHHILLDGWSRGIIMGELFHMYHMKEARQKHRLEEARPYSDYIGWLQQQDKEAAKAYWRNYLSGFTE
KSPISVLAGSSGHAKYKRKEAVIEFPEQLTGRITELASRNNVTFHTVLQCIWGMLLARYNQTDDVVFGTVISGRDAQVTGIEKMVGLFINTVPTRIRLDKSQSFKELIKSVQEQAL
EGRTYHDMNLSEVQSLSELKRELLDHILIFENYAVDQSAFETSGKRGAGFVFEEIHAEEQTNYGFNIVAVPGERLVIKLTYDGNIYHDHIIAGIKGHLQQVMEQVVQHEDQSLNDI
TVLSEAERNRLLYEWNDTKAEYPNQTIHRLFEEQAEKTPELAAVVSGNDKLTYRELNEKSNQLARYLRDKGVKADTIVAIMAERSPEMVVGIMGILKAGGAYLPIDPDYPEERIK
YMLEDSGAAIILADHKQDLGTLHQEAVELTGDFSSYPADNLEPAGNADSLAYIIYTSGSTGKPKGVMIRQRGLVNYITWADRVYVQGEQLDFALYSSIAFDLTVTSIFTPLISGNR
VIVYRHSEDGEPLIRKVFRDQKAGIVKLTPSHLSLVKDMDASGSSIKRLIVGGEDLKTELAKEITERFHHNIEIYNEYGPTETVVGCMIYQYDAGWDRQVSVPIGKPASNVQLYIL
DERQEVQPVGIAGELYISGDGVAKGYLNKPELTSEKFLPNPFLPGERMYRTGDLAKMRPDGHIEYLGRIDHQVKIRGYRIELGEIEHQLLRHSDIKEAAVAAKTDQNNDQVLCA
YVVSERDITQKDIKTFLAKELPEYMVPSYLLKLDELPLTPNGKVDLKALPEPDRSAGALLEYEPPRHELEEKMAAIWEDILNIEQIGINANIFDIGANSLNVMSFVSRLYAELGFRV
PFKDIFSKPTIKELSDFLKHAQDLLKDYTDDCMQLTRAEEGGKNLFCFPPAASMGIAYMGLAKHLKQHSVYSFNFIPSANRIRKYADIIKNIQGEGPYTLIGYSSGGILAFDVAKEL
NRQGYEVEDLIIIDSKYRTKAEKHQFTEEEYREEISKTFELEKYRDVEKLLSDYLVDLVMKSYVYIQNTVTTGAIDGHISYIKSSDNQRGENMMMWEKATSKTFTVVQGAGTHM
QMISKSHPDILERNARLIHDIINKTVKIGSRSHHHHHH

#### 8) BacC4-C5[A-PCP-E-COM(Δ5062-5084)-C-A-PCP-TE]-H$_6$

MDSDEEMNALLDQNQGKQADYPQDQTVHQLFEQQADKTPEQTAVVYADEKLTYRELNERANQLARLLRDKGADADQPVAIMIEPSLEMIISMLAVLKAGAAYVPIEPEQLAKR
TNEILSDSRAAILLVKGSVKENVAFAGEIVNVADGLIDAKVASNLSASGSADQNAYIIYTSGSTGKPKGVFVRHGNVVNYTTWFMKEAGLTENDKAMLVSSYAFDLGYTSIFSAL
LSGSELHIARKECYTNAHRALKYIKENGITYIKLTPSLFNIFVNDPGFSAEKPCATLRLVVLGGEMINTRDVETFYNQYPDHVVMNHYGPTETTIGSVFKVIDPEHLDSFKECPVIG
TPIHNTNAYVLDENMKLLPEGVYGELCIAGAGVTGGYVNRPDETKEKFIENPFAPHTKMYRTGDLARRLSDGNIELAGRIDTQVKVRGYRIEPEEIKNRLLAHDDIKEAFIAARED
HKGAKQLCAYFTADAELPFEDIRTYLMHELPEYMIPSSFVQIEKMPLSANGKIDTAALPEPQPGKETEYEPPRNETEEKLVQIWEEVLGIDKIGITHHFFAAGGDSIKALQMISRLS
REGLSLEMKDLFANPQIKSLSRYVKAESDKSASYETVEGEVLLTPIQQEYFSLNKTDRNHYNHAVMLYRKNGFDESIVKRVFKEIIKHHDALRTVFTEEDGKIIQYNRGPDKQLF
DLFVYDVSSENDQPQKVYQLATELQQSIDIETGPLVKLALFKTNNGDHLLIIIHHLVVDGISWRILFEDLAIGYSQLANGEKVEFYPKTASYQAYARHIAEYAKSVKLLSEKQYWLK
AIAEGVEFLDMNENAGAFKVEDSRTFSTELEKEETKRLLRETNRAYHTEINDILITALLVAARDMNGQNQLRITLEGHGREQVADGIDISRTVGWFTSKYPVFIDLGQETDMSRTI
KMVKEHLRNVPNKGIGYGILKYLTRDSEIAKGAASPILFNYLGQLDEDINSGEFSSSHLSPGEAAGKGITREHPLEINAVVFRGKLAIQTTYNTRAYSEDVVRAFAQNYKEALKAV
IRHCAEREETEKGMEIEKIYPLANMQRGMLFHALEDKESQAYFEQMAINMKGLIDERLFAETFNDIMERHEILRASIEYEITDEPRNVIIKDRKINLDYHDLRKQSPAEREQVIQAY
RKADREKGFRLNSEPLIRAALMRTEDDSYTFIWTNHHILLDGWSRGIIMGELFHMYHMKEARQKHRLEEARPYSDYIGWLQQQDKEAAKAYWRNYLSGFTEKSPISVLAGSSG
HAKYKRKEAVIEFPEQLTGRITELASRNNVTFHTVLQCIWGMLLARYNQTDDVVFGTVISGRDAQVTGIEKMVGLFINTVPTRIRLDKSQSFKELIKSVQEQALEGRTYHDMNLS
EVQSLSELKRELLDHILIFENYAVDQSAFETSGKRGAGFVFEEIHAEEQTNYGFNIVAVPGERLVIKLTYDGNIYHDHIIAGIKGHLQQVMEQVVQHEDQSLNDITVLSEAERNRLL
YEWNDTKAEYPNQTIHRLFEEQAEKTPELAAVVSGNDKLTYRELNEKSNQLARYLRDKGVKADTIVAIMAERSPEMVVGIMGILKAGGAYLPIDPDYPEERIKYMLEDSGAAIILA
DHKQDLGTLHQEAVELTGDFSSYPADNLEPAGNADSLAYIIYTSGSTGKPKGVMIRQRGLVNYITWADRVYVQGEQLDFALYSSIAFDLTVTSIFTPLISGNRVIVYRHSEDGEP
LIRKVFRDQKAGIVKLTPSHLSLVKDMDASGSSIKRLIVGGEDLKTELAKEITERFHHNIEIYNEYGPTETVVGCMIYQYDAGWDRQVSVPIGKPASNVQLYILDERQEVQPVGIA
GELYISGDGVAKGYLNKPELTSEKFLPNPFLPGERMYRTGDLAKMRPDGHIEYLGRIDHQVKIRGYRIELGEIEHQLLRHSDIKEAAVAAKTDQNNDQVLCAYVVSERDITQKDI
KTFLAKELPEYMVPSYLLKLDELPLTPNGKVDLKALPEPDRSAGALLEYEPPRHELEEKMAAIWEDILNIEQIGINANIFDIGANSLNVMSFVSRLYAELGFRVPFKDIFSKPTIKEL
SDFLKHAQDLLKDYTDDCMQLTRAEEGGKNLFCFPPAASMGIAYMGLAKHLKQHSVYSFNFIPSANRIRKYADIIKNIQGEGPYTLIGYSSGGILAFDVAKELNRQGYEVEDLIIID
SKYRTKAEKHQFTEEEYREEISKTFELEKYRDVEKLLSDYLVDLVMKSYVYIQNTVTTGAIDGHISYIKSSDNQRGENMMMWEKATSKTFTVVQGAGTHMQMISKSHPDILERN
ARLIHDIINKTVKIGSRSHHHHHH

### 9) BacC4-C5[A-PCP-E-COM(Ins5062(GGS)$_3$)-C-A-PCP-TE]-H$_6$

MDSDEEMNALLDQNQGKQADYPQDQTVHQLFEQQADKTPEQTAVVYADEKLTYRELNERANQLARLLRDKGADADQPVAIMIEPSLEMIISMLAVLKAGAAYVPIEPEQLAKR
TNEILSDSRAAILLVKGSVKENVAFAGEIVNVADGLIDAKVASNLSASGSADQNAYIIYTSGSTGKPKGVFVRHGNVVNYTTWFMKEAGLTENDKAMLVSSYAFDLGYTSIFSAL
LSGSELHIARKECYTNAHRALKYIKENGITYIKLTPSLFNIFVNDPGFSAEKPCATLRLVVLGGEMINTRDVETFYNQYPDHVVMNHYGPTETTIGSVFKVIDPEHLDSFKECPVIG
TPIHNTNAYVLDENMKLLPEGVYGELCIAGAGVTGGYVNRPDETKEKFIENPFAPHTKMYRTGDLARRLSDGNIELAGRIDTQVKVRGYRIEPEEIKNRLLAHDDIKEAFIAARED
HKGAKQLCAYFTADAELPFEDIRTYLMHELPEYMIPSSFVQIEKMPLSANGKIDTAALPEPQPGKETEYEPPRNETEEKLVQIWEEVLGIDKIGITHHFFAAGGDSIKALQMISRLS
REGLSLEMKDLFANPQIKSLSRYVKAESDKSASYETVEGEVLLTPIQQEYFSLNKTDRNHYNHAVMLYRKNGFDESIVKRVFKEIIKHHDALRTVFTEEDGKIIQYNRGPDKQLF
DLFVYDVSSENDQPQKVYQLATELQQSIDIETGPLVKLALFKTNNGDHLLIIIHHLVVDGISWRILFEDLAIGYSQLANGEKVEFYPKTASYQAYARHIAEYAKSVKLLSEKQYWLK
AIAEGVEFLDMNENAGAFKVEDSRTFSTELEKEETKRLLRETNRAYHTEINDILITALLVAARDMNGQNQLRITLEGHGREQVADGIDISRTVGWFTSKYPVFIDLGQETDMSRTI
KMVKEHLRNVPNKGIGYGILKYLTRDSEIAKGAASPILFNYLGQLDEDINSGEFSSSHLSPGEAAGKGITREHPLEINAVVFRGKLAIQTTYNTRAYSEDVVRAFAQNYKEALKAV
IRHCAEREETEKggsggsggsTPSDYGDKGISLDQLEEIKLKYKGMEIEKIYPLANMQRGMLFHALEDKESQAYFEQMAINMKGLIDERLFAETFNDIMERHEILRASIEYEITDEPR
NVIIKDRKINLDYHDLRKQSPAEREQVIQAYRKADREKGFRLNSEPLIRAALMRTEDDSYTFIWTNHHILLDGWSRGIIMGELFHMYHMKEARQKHRLEEARPYSDYIGWLQQQ
DKEAAKAYWRNYLSGFTEKSPISVLAGSSGHAKYKRKEAVIEFPEQLTGRITELASRNNVTFHTVLQCIWGMLLARYNQTDDVVFGTVISGRDAQVTGIEKMVGLFINTVPTRIR
LDKSQSFKELIKSVQEQALEGRTYHDMNLSEVQSLSELKRELLDHILIFENYAVDQSAFETSGKRGAGFVFEEIHAEEQTNYGFNIVAVPGERLVIKLTYDGNIYHDHIIAGIKGHL
QQVMEQVVQHEDQSLNDITVLSEAERNRLLYEWNDTKAEYPNQTIHRLFEEQAEKTPELAAVVSGNDKLTYRELNEKSNQLARYLRDKGVKADTIVAIMAERSPEMVVGIMGIL
KAGGAYLPIDPDYPEERIKYMLEDSGAAIILADHKQDLGTLHQEAVELTGDFSSYPADNLEPAGNADSLAYIIYTSGSTGKPKGVMIRQRGLVNYITWADRVYVQGEQLDFALYS
SIAFDLTVTSIFTPLISGNRVIVYRHSEDGEPLIRKVFRDQKAGIVKLTPSHLSLVKDMDASGSSIKRLIVGGEDLKTELAKEITERFHHNIEIYNEYGPTETVVGCMIYQYDAGWDR
QVSVPIGKPASNVQLYILDERQEVQPVGIAGELYISGDGVAKGYLNKPELTSEKFLPNPFLPGERMYRTGDLAKMRPDGHIEYLGRIDHQVKIRGYRIELGEIEHQLLRHSDIKEA
AVAAKTDQNNDQVLCAYVVSERDITQKDIKTFLAKELPEYMVPSYLLKLDELPLTPNGKVDLKALPEPDRSAGALLEYEPPRHELEEKMAAIWEDILNIEQIGINANIFDIGANSLN
VMSFVSRLYAELGFRVPFKDIFSKPTIKELSDFLKHAQDLLKDYTDDCMQLTRAEEGGKNLFCFPPAASMGIAYMGLAKHLKQHSVYSFNFIPSANRIRKYADIIKNIQGEGPYTL
IGYSSGGILAFDVAKELNRQGYEVEDLIIIDSKYRTKAEKHQFTEEEYREEISKTFELEKYRDVEKLLSDYLVDLVMKSYVYIQNTVTTGAIDGHISYIKSSDNQRGENMMMWEKA
TSKTFTVVQGAGTHMQMISKSHPDILERNARLIHDIINKTVKIGSRSHHHHHH

### 10) BacC4-C5[A-PCP-E-COM(Δ5059-5061)-C-A-PCP-TE]-H$_6$

MDSDEEMNALLDQNQGKQADYPQDQTVHQLFEQQADKTPEQTAVVYADEKLTYRELNERANQLARLLRDKGADADQPVAIMIEPSLEMIISMLAVLKAGAAYVPIEPEQLAKR
TNEILSDSRAAILLVKGSVKENVAFAGEIVNVADGLIDAKVASNLSASGSADQNAYIIYTSGSTGKPKGVFVRHGNVVNYTTWFMKEAGLTENDKAMLVSSYAFDLGYTSIFSAL
LSGSELHIARKECYTNAHRALKYIKENGITYIKLTPSLFNIFVNDPGFSAEKPCATLRLVVLGGEMINTRDVETFYNQYPDHVVMNHYGPTETTIGSVFKVIDPEHLDSFKECPVIG
TPIHNTNAYVLDENMKLLPEGVYGELCIAGAGVTGGYVNRPDETKEKFIENPFAPHTKMYRTGDLARRLSDGNIELAGRIDTQVKVRGYRIEPEEIKNRLLAHDDIKEAFIAARED
HKGAKQLCAYFTADAELPFEDIRTYLMHELPEYMIPSSFVQIEKMPLSANGKIDTAALPEPQPGKETEYEPPRNETEEKLVQIWEEVLGIDKIGITHHFFAAGGDSIKALQMISRLS
REGLSLEMKDLFANPQIKSLSRYVKAESDKSASYETVEGEVLLTPIQQEYFSLNKTDRNHYNHAVMLYRKNGFDESIVKRVFKEIIKHHDALRTVFTEEDGKIIQYNRGPDKQLF
DLFVYDVSSENDQPQKVYQLATELQQSIDIETGPLVKLALFKTNNGDHLLIIIHHLVVDGISWRILFEDLAIGYSQLANGEKVEFYPKTASYQAYARHIAEYAKSVKLLSEKQYWLK
AIAEGVEFLDMNENAGAFKVEDSRTFSTELEKEETKRLLRETNRAYHTEINDILITALLVAARDMNGQNQLRITLEGHGREQVADGIDISRTVGWFTSKYPVFIDLGQETDMSRTI
KMVKEHLRNVPNKGIGYGILKYLTRDSEIAKGAASPILFNYLGQLDEDINSGEFSSSHLSPGEAAGKGITREHPLEINAVVFRGKLAIQTTYNTRAYSEDVVRAFAQNYKEALKAV
IRHCAEREETPSDYGDKGISLDQLEEIKLKYKGMEIEKIYPLANMQRGMLFHALEDKESQAYFEQMAINMKGLIDERLFAETFNDIMERHEILRASIEYEITDEPRNVIIKDRKINLD
YHDLRKQSPAEREQVIQAYRKADREKGFRLNSEPLIRAALMRTEDDSYTFIWTNHHILLDGWSRGIIMGELFHMYHMKEARQKHRLEEARPYSDYIGWLQQQDKEAAKAYWR
NYLSGFTEKSPISVLAGSSGHAKYKRKEAVIEFPEQLTGRITELASRNNVTFHTVLQCIWGMLLARYNQTDDVVFGTVISGRDAQVTGIEKMVGLFINTVPTRIRLDKSQSFKELI
KSVQEQALEGRTYHDMNLSEVQSLSELKRELLDHILIFENYAVDQSAFETSGKRGAGFVFEEIHAEEQTNYGFNIVAVPGERLVIKLTYDGNIYHDHIIAGIKGHLQQVMEQVVQ
HEDQSLNDITVLSEAERNRLLYEWNDTKAEYPNQTIHRLFEEQAEKTPELAAVVSGNDKLTYRELNEKSNQLARYLRDKGVKADTIVAIMAERSPEMVVGIMGILKAGGAYLPID
PDYPEERIKYMLEDSGAAIILADHKQDLGTLHQEAVELTGDFSSYPADNLEPAGNADSLAYIIYTSGSTGKPKGVMIRQRGLVNYITWADRVYVQGEQLDFALYSSIAFDLTVTSI
FTPLISGNRVIVYRHSEDGEPLIRKVFRDQKAGIVKLTPSHLSLVKDMDASGSSIKRLIVGGEDLKTELAKEITERFHHNIEIYNEYGPTETVVGCMIYQYDAGWDRQVSVPIGKP
ASNVQLYILDERQEVQPVGIAGELYISGDGVAKGYLNKPELTSEKFLPNPFLPGERMYRTGDLAKMRPDGHIEYLGRIDHQVKIRGYRIELGEIEHQLLRHSDIKEAAVAAKTDQ
NNDQVLCAYVVSERDITQKDIKTFLAKELPEYMVPSYLLKLDELPLTPNGKVDLKALPEPDRSAGALLEYEPPRHELEEKMAAIWEDILNIEQIGINANIFDIGANSLNVMSFVSRL
YAELGFRVPFKDIFSKPTIKELSDFLKHAQDLLKDYTDDCMQLTRAEEGGKNLFCFPPAASMGIAYMGLAKHLKQHSVYSFNFIPSANRIRKYADIIKNIQGEGPYTLIGYSSGGIL
AFDVAKELNRQGYEVEDLIIIDSKYRTKAEKHQFTEEEYREEISKTFELEKYRDVEKLLSDYLVDLVMKSYVYIQNTVTTGAIDGHISYIKSSDNQRGENMMMWEKATSKTFTVV
QGAGTHMQMISKSHPDILERNARLIHDIINKTVKIGSRSHHHHHH

### 11) BacC4-C5[A-PCP-E-COM(Ins5062G)-C-A-PCP-TE]-H$_6$

MDSDEEMNALLDQNQGKQADYPQDQTVHQLFEQQADKTPEQTAVVYADEKLTYRELNERANQLARLLRDKGADADQPVAIMIEPSLEMIISMLAVLKAGAAYVPIEPEQLAKR
TNEILSDSRAAILLVKGSVKENVAFAGEIVNVADGLIDAKVASNLSASGSADQNAYIIYTSGSTGKPKGVFVRHGNVVNYTTWFMKEAGLTENDKAMLVSSYAFDLGYTSIFSAL
LSGSELHIARKECYTNAHRALKYIKENGITYIKLTPSLFNIFVNDPGFSAEKPCATLRLVVLGGEMINTRDVETFYNQYPDHVVMNHYGPTETTIGSVFKVIDPEHLDSFKECPVIG
TPIHNTNAYVLDENMKLLPEGVYGELCIAGAGVTGGYVNRPDETKEKFIENPFAPHTKMYRTGDLARRLSDGNIELAGRIDTQVKVRGYRIEPEEIKNRLLAHDDIKEAFIAARED
HKGAKQLCAYFTADAELPFEDIRTYLMHELPEYMIPSSFVQIEKMPLSANGKIDTAALPEPQPGKETEYEPPRNETEEKLVQIWEEVLGIDKIGITHHFFAAGGDSIKALQMISRLS
REGLSLEMKDLFANPQIKSLSRYVKAESDKSASYETVEGEVLLTPIQQEYFSLNKTDRNHYNHAVMLYRKNGFDESIVKRVFKEIIKHHDALRTVFTEEDGKIIQYNRGPDKQLF
DLFVYDVSSENDQPQKVYQLATELQQSIDIETGPLVKLALFKTNNGDHLLIIIHHLVVDGISWRILFEDLAIGYSQLANGEKVEFYPKTASYQAYARHIAEYAKSVKLLSEKQYWLK
AIAEGVEFLDMNENAGAFKVEDSRTFSTELEKEETKRLLRETNRAYHTEINDILITALLVAARDMNGQNQLRITLEGHGREQVADGIDISRTVGWFTSKYPVFIDLGQETDMSRTI
KMVKEHLRNVPNKGIGYGILKYLTRDSEIAKGAASPILFNYLGQLDEDINSGEFSSSHLSPGEAAGKGITREHPLEINAVVFRGKLAIQTTYNTRAYSEDVVRAFAQNYKEALKAV
IRHCAEREETEKgTPSDYGDKGISLDQLEEIKLKYKGMEIEKIYPLANMQRGMLFHALEDKESQAYFEQMAINMKGLIDERLFAETFNDIMERHEILRASIEYEITDEPRNVIIKDRK
INLDYHDLRKQSPAEREQVIQAYRKADREKGFRLNSEPLIRAALMRTEDDSYTFIWTNHHILLDGWSRGIIMGELFHMYHMKEARQKHRLEEARPYSDYIGWLQQQDKEAAKA
YWRNYLSGFTEKSSISVLAGSSGHAKYKRKEAVIEFPEQLTGRITELASRNNVTFHTVLQCIWGMLLARYNQTDDVVFGTVISGRDAQVTGIEKMVGLFINTVPTRIRLDKSQSF
KELIKSVQEQALEGRTYHDMNLSEVQSLSELKRELLDHILIFENYAVDQSAFETSGKRGAGFVFEEIHAEEQTNYGFNIVAVPGERLVIKLTYDGNIYHDHIIAGIKGHLQQVMEQ
VVQHEDQSLNDITVLSEAERNRLLYEWNDTKAEYPNQTIHRLFEEQAEKTPELAAVVSGNDKLTYRELNEKSNQLARYLRDKGVKADTIVAIMAERSPEMVVGIMGILKAGGAY
LPIDPDYPEERIKYMLEDSGAAIIPADHKQDLGTLHQEAVELTGDFSSYPADNLEPAGNADSLAYIIYTSGSTGKPKGVMIRQRGLVNYITWADRVYVQGEQLDFALYSSIAFDLT
VTSIFTPLISGNRVIVYRHSEDGEPLIRKVFRDQKAGIVKLTPSHLSLVKDMDASGSSIKRLIVGGEDLKTELAKEITERFHHNIEIYNEYGPTETVVGCMIYQYDAGWDRQVSVPI
GKPASNVQLYILDERQEVQPVGIAGELYISGDGVAKGYLNKPELTSEKFLPNPFLPGERMYRTGDLAKMRPDGHIEYLGRIDHQVKIRGYRIELGEIEHQLLRHSDIKEAAVAAK
TDQNNDQVLCAYVVSERDITQKDIKTFLAKELPEYMVPSYLLKLDELPLTPNGKVDLKALPEPDRSAGALLEYEPPRHELEEKMAAIWEDILNIEQIGINANIFDIGANSLNVMSFV
SRLYAELGFRVPFKDIFSKPTIKELSDFLKHAQDLLKDYTDDCMQLTRAEEGGKNLFCFPPAASMGIAYMGLAKHLKQHSVYSFNFIPSANRIRKYADIIKNIQGEGPYTLIGYSS
GGILAFDVAKELNRQGYEVEDLIIIDSKYRTKAEKHQFTEEEYREEISKTFELEKYRDVEKLLSDYLVDLVMKSYVYIQNTVTTGAIDGHISYIKSSDNQRGENMMMWEKATSKTF
TVVQGAGTHMQMISKSHPDILERNARLIHDIINKTVKIGSRSHHHHHH

### 12) BacC4-C5[A-PCP-E-COM(Δ5059)-C-A-PCP-TE]-$H_6$

MDSDEEMNALLDQNQGKQADYPQDQTVHQLFEQQADKTPEQTAVVYADEKLTYRELNERANQLARLLRDKGADADQPVAIMIEPSLEMIISMLAVLKAGAAYVPIEPEQLAKR
TNEILSDSRAAILLVKGSVKENVAFAGEIVNVADGLIDAKVASNLSASGSADQNAYIIYTSGSTGKPKGVFVRHGNVVNYTTWFMKEAGLTENDKAMLVSSYAFDLGYTSIFSAL
LSGSELHIARKECYTNAHRALKYIKENGITYIKLTPSLFNIFVNDPGFSAEKPCATLRLVVLGGEMINTRDVETFYNQYPDHVVMNHYGPTETTIGSVFKVIDPEHLDSFKECPVIG
TPIHNTNAYVLDENMKLLPEGVYGELCIAGAGVTGGYVNRPDETKEKFIENPFAPHTKMYRTGDLARRLSDGNIELAGRIDTQVKVRGYRIEPEEIKNRLLAHDDIKEAFIAARED
HKGAKQLCAYFTADAELPFEDIRTYLMHELPEYMIPSSFVQIEKMPLSANGKIDTAALPEPQPGKETEYEPPRNETEEKLVQIWEEVLGIDKIGITHHFFAAGGDSIKALQMISRLS
REGLSLEMKDLFANPQIKSLSRYVKAESDKSASYETVEGEVLLTPIQQEYFSLNKTDRNHYNHAVMLYRKNGFDESIVKRVFKEIIKHHDALRTVFTEEDGKIIQYNRGPDKQLF
DLFVYDVSSENDQPQKVYQLATELQQSIDIETGPLVKLALFKTNNGDHLLIIIHHLVVDGISWRILFEDLAIGYSQLANGEKVEFYPKTASYQAYARHIAEYAKSVKLLSEKQYWLK
AIAEGVEFLDMNENAGAFKVEDSRTFSTELEKEETKRLLRETNRAYHTEINDILITALLVAARDMNGQNQLRITLEGHGREQVADGIDISRTVGWFTSKYPVFIDLGQETDMSRTI
KMVKEHLRNVPNKGIGYGILKYLTRDSEIAKGAASPILFNYLGQLDEDINSGEFSSSHLSPGEAAGKGITREHPLEINAVVFRGKLAIQTTYNTRAYSEDVVRAFAQNYKEALKAV
IRHCAEREEEKTPSDYGDKGISLDQLEEIKLKYKGMEIEKIYPLANMQRGMLFHALEDKESQAYFEQMAINMKGLIDERLFAETFNDIMERHEILRASIEYEITDEPRNVIIKDRKIN
LDYHDLRKQSPAEREQVIQAYRKADREKGFRLNSEPLIRAALMRTEDDSYTFIWTNHHILLDGWSRGIIMGELFHMYHMKEARQKHRLEEARPYSDYIGWLQQQDKEAAKAY
WRNYLSGFTEKSPISVLAGSSGHAKYKRKEAVIEFPEQLTGRITELASRNNVTFHTVLQCIWGMLLARYNQTDDVVFGTVISGRDAQVTGIEKMVGLFINTVPTRIRLDKSQSFK
ELIKSVQEQALEGRTYHDMNLSEVQSLSELKRELLDHILIFENYAVDQSAFETSGKRGAGFVFEEIHAEEQTNYGFNIVAVPGERLVIKLTYDGNIYHDHIIAGIKGHLQQVMEQV
VQHEDQSLNDITVLSEAERNRLLYEWNDTKAEYPNQTIHRLFEEQAEKTPELAAVVSGNDKLTYRELNEKSNQLARYLRDKGVKADTIVAIMAERSPEMVVGIMGILKAGGAYL
PIDPDYPEERIKYMLEDSGAAIILADHKQDLGTLHQEAVELTGDFSSYPADNLEPAGNADSLAYIIYTSGSTGKPKGVMIRQRGLVNYITWADRVYVQGEQLDFALYSSIAFDLTV
TSIFTPLISGNRVIVYRHSEDGEPLIRKVFRDQKAGIVKLTPSHLSLVKDMDASGSSIKRLIVGGEDLKTELAKEITERFHHNIEIYNEYGPTETVVGCMIYQYDAGWDRQVSVPIG
KPASNVQLYILDERQEVQPVGIAGELYISGDGVAKGYLNKPELTSEKFLPNPFLPGERMYRTGDLAKMRPDGHIEYLGRIDHQVKIRGYRIELGEIEHQLLRHSDIKEAAVAAKTD
QNNDQVLCAYVVSERDITQKDIKTFLAKELPEYMVPSYLLKLDELPLTPNGKVDLKALPEPDRSAGALLEYEPPRHELEEKMAAIWEDILNIEQIGINANIFDIGANSLNVMSFVSR
LYAELGFRVPFKDIFSKPTIKELSDFLKHAQDLLKDYTDDCMQLTRAEEGGKNLFCFPPAASMGIAYMGLAKHLKQHSVYSFNFIPSANRIRKYADIIKNIQGEGPYTLIGYSSGGI
LAFDVAKELNRQGYEVEDLIIIDSKYRTKAEKHQFTEEEYREEISKTFELEKYRDVEKLLSDYLVDLVMKSYVYIQNTVTTGAIDGHISYIKSSDNQRGENMMMWEKATSKTFTVV
QGAGTHMQMISKSHPDILERNARLIHDIINKTVKIGSRSHHHHHH

### 13) BacC4-C5[A-PCP-E-COM(Ins5068G)-C-A-PCP-TE]-$H_6$

MDSDEEMNALLDQNQGKQADYPQDQTVHQLFEQQADKTPEQTAVVYADEKLTYRELNERANQLARLLRDKGADADQPVAIMIEPSLEMIISMLAVLKAGAAYVPIEPEQLAKR
TNEILSDSRAAILLVKGSVKENVAFAGEIVNVADGLIDAKVASNLSASGSADQNAYIIYTSGSTGKPKGVFVRHGNVVNYTTWFMKEAGLTENDKAMLVSSYAFDLGYTSIFSAL
LSGSELHIARKECYTNAHRALKYIKENGITYIKLTPSLFNIFVNDPGFSAEKPCATLRLVVLGGEMINTRDVETFYNQYPDHVVMNHYGPTETTIGSVFKVIDPEHLDSFKECPVIG
TPIHNTNAYVLDENMKLLPEGVYGELCIAGAGVTGGYVNRPDETKEKFIENPFAPHTKMYRTGDLARRLSDGNIELAGRIDTQVKVRGYRIEPEEIKNRLLAHDDIKEAFIAARED
HKGAKQLCAYFTADAELPFEDIRTYLMHELPEYMIPSSFVQIEKMPLSANGKIDTAALPEPQPGKETEYEPPRNETEEKLVQIWEEVLGIDKIGITHHFFAAGGDSIKALQMISRLS
REGLSLEMKDLFANPQIKSLSRYVKAESDKSASYETVEGEVLLTPIQQEYFSLNKTDRNHYNHAVMLYRKNGFDESIVKRVFKEIIKHHDALRTVFTEEDGKIIQYNRGPDKQLF
DLFVYDVSSENDQPQKVYQLATELQQSIDIETGPLVKLALFKTNNGDHLLIIIHHLVVDGISWRILFEDLAIGYSQLANGEKVEFYPKTASYQAYARHIAEYAKSVKLLSEKQYWLK
AIAEGVEFLDMNENAGAFKVEDSRTFSTELEKEETKRLLRETNRAYHTEINDILITALLVAARDMNGQNQLRITLEGHGREQVADGIDISRTVGWFTSKYPVFIDLGQETDMSRTI
KMVKEHLRNVPNKGIGYGILKYLTRDSEIAKGAASPILFNYLGQLDEDINSGEFSSSHLSPGEAAGKGITREHPLEINAVVFRGKLAIQTTYNTRAYSEDVVRAFAQNYKEALKAV
IRHCAEREETEKTPSDYgGDKGISLDQLEEIKLKYKGMEIEKIYPLANMQRGMLFHALEDKESQAYFEQMAINMKGLIDERLFAETFNDIMERHEILRASIEYEITDEPRNVIIKDRK
INLDYHDLRKQSPAEREQVIQAYRKADREKGFRLNSEPLIRAALMRTEDDSYTFIWTNHHILLDGWSRGIIMGELFHMYHMKEARQKHRLEEARPYSDYIGWLQQQDKEAAKA
YWRNYLSGFTEKSPISVLAGSSGHAKYKRKEAVIEFPEQLTGRITELASRNNVTFHTVLQCIWGMLLARYNQTDDVVFGTVISGRDAQVTGIEKMVGLFINTVPTRIRLDKSQSF
KELIKSVQEQALEGRTYHDMNLSEVQSLSELKRELLDHILIFENYAVDQSAFETSGKRGAGFVFEEIHAEEQTNYGFNIVAVPGERLVIKLTYDGNIYHDHIIAGIKGHLQQVMEQ
VVQHEDQSLNDITVLSEAERNRLLYEWNDTKAEYPNQTIHRLFEEQAEKTPELAAVVSGNDKLTYRELNEKSNQLARYLRDKGVKADTIVAIMAERSPEMVVGIMGILKAGGAY
LPIDPDYPEERIKYMLEDSGAAIILADHKQDLGTLHQEAVELTGDFSSYPADNLEPAGNADSLAYIIYTSGSTGKPKGVMIRQRGLVNYITWADRVYVQGEQLDFALYSSIAFDLT
VTSIFTPLISGNRVIVYRHSEDGEPLIRKVFRDQKAGIVKLTPSHLSLVKDMDASGSSIKRLIVGGEDLKTELAKEITERFHHNIEIYNEYGPTETVVGCMIYQYDAGWDRQVSVPI
GKPASNVQLYILDERQEVQPVGIAGELYISGDGVAKGYLNKPELTSEKFLPNPFLPGERMYRTGDLAKMRPDGHIEYLGRIDHQVKIRGYRIELGEIEHQLLRHSDIKEAAVAAK
TDQNNDQVLCAYVVSERDITQKDIKTFLAKELPEYMVPSYLLKLDELPLTPNGKVDLKALPEPDRSAGALLEYEPPRHELEEKMAAIWEDILNIEQIGINANIFDIGANSLNVMSFV
SRLYAELGFRVPFKDIFSKPTIKELSDFLKHAQDLLKDYTDDCMQLTRAEEGGKNLFCFPPAASMGIAYMGLAKHLKQHSVYSFNFIPSANRIRKYADIIKNIQGEGPYTLIGYSS
GGILAFDVAKELNRQGYEVEDLIIIDSKYRTKAEKHQFTEEEYREEISKTFELEKYRDVEKLLSDYLVDLVMKSYVYIQNTVTTGAIDGHISYIKSSDNQRGENMMMWEKATSKTF
TVVQGAGTHMQMISKSHPDILERNARLIHDIINKTVKIGSRSHHHHHH

### 14) BacC4-C5[A-PCP-E-COM(P5063A)-C-A-PCP-TE]-$H_6$

MDSDEEMNALLDQNQGKQADYPQDQTVHQLFEQQADKTPEQTAVVYADEKLTYRELNERANQLARLLRDKGADADQPVAIMIEPSLEMIISMLAVLKAGAAYVPIEPEQLTKR
TNEILSDSRAAILLVKGSVKENVAFAGEIVNVADGLIDAKVASNLSASGSADQNAYIIYTSGSTGKPKGVFVRHGNVVNYTTWFMKEAGLTENDKAMLVSSYAFDLGYTSIFSAL
LSGSELHIARKECYTNAHRALKYIKENGITYIKLTPSLFNIFVNDPGFSAEKPCATLRLVVLGGEMINTRDVETFYNQYPDHVVMNHYGPTETTIGSVFKVIDPEHLDSFKECPVIG
TPIHNTNAYVLDENMKLLPEGVYGELCIAGAGVTGGYVNRPDETKEKFIENPFAPHTKMYRTGDLARRLSDGNIELAGRIDTQVKVRGYRIEPEEIKNRLLAHDDIKEAFIAARED
HKGAKQLCAYFTADAELPFEDIRTYLMHELPEYMIPSSFVQIEKMPLSANGKIDTAALPEPQPGKETEYEPPRNETEEKLVQIWEEVLGIDKIGITHHFFAAGGDSIKALQMISRLS
REGLSLEMKDLFANPQIKSLSRYVKAESDKSASYETVEGEVLLTPIQQEYFSLNKTDRNHYNHAVMLYRKNGFDESIVKRVFKEIIKHHDALRTVFTEEDGKIIQYNRGPDKQLF
DLFVYDVSSENDQPQKVYQLATELQQSIDIETGPLVKLALFKTNNGDHLLIIIHHLVVDGISWRILFEDLAIGYSQLANGEKVEFYPKTASYQAYARHIAEYAKSVKLLSEKQYWLK
AIAEGVEFLDMNENAGAFKVEDSRTFSTELEKEETKRLLRETNRAYHTEINDILITALLVAARDMNGQNQLRITLEGHGREQVADGIDISRTVGWFTSKYPVFIDLGQETDMSRTI
KMVKEHLRNVPNKGIGYGILKYLTRDSEIAKGAASPILFNYLGQLDEDINSGEFSSSHLSPGEAAGKGITREHPLEINAVVFRGKLAIQTTYNTRAYSEDVVRAFAQNYKEALKAV
IRHCAEREETEKTASDYGDKGISLDQLEEIKLKYKGMEIEKIYPLANMQRGMLFHALEDKESQAYFEQMAINMKGLIDERLFAETFNDIMERHEILRASIEYEITDEPRNVIIKDRKI
NLDYHDLRKQSPAEREQVIQAYRKADREKGFRLNSEPLIRAALMRTEDDSYTFIWTNHHILLDGWSRGIIMGELFHMYHMKEARQKHRLEEARPYSDYIGWLQQQDKEAAKAY
WRNYLSGFTEKSPISVLAGSSGHAKYKRKEAVIEFPEQLTGRITELASRNNVTFHTVLQCIWGMLLARYNQTDDVVFGTVISGRDAQVTGIEKMVGLFINTVPTRIRLDKSQSFK
ELIKSVQEQALEGRTYHDMNLSEVQSLSELKRELLDHILIFENYAVDQSAFETSGKRGAGFVFEEIHAEEQTNYGFNIVAVPGERLVIKLTYDGNIYHDHIIAGIKGHLQQVMEQV
VQHEDQSLNDITVLSEAERNRLLYEWNDTKAEYPNQTIHRLFEEQAEKTPELAAVVSGNDKLTYRELNEKSNQLARYLRDKGVKADTIVAIMAERSPEMVVGIMGILKAGGAYL
PIDPDYPEERIKYMLEDSGAAIILADHKQDLGTLHQEAVELTGDFSSYPADNLEPAGNADSLAYIIYTSGSTGKPKGVMIRQRGLVNYITWADRVYVQGEQLDFALYSSIAFDLTV
TSIFTPLISGNRVIVYRHSEDGEPLIRKVFRDQKAGIVKLTPSHLSLVKDMDASGSSIKRLIVGGEDLKTELAKEITERFHHNIEIYNEYGPTETVVGCMIYQYDAGWDRQVSVPIG
KPASNVQLYILDERQEVQPVGIAGELYISGDGVAKGYLNKPELTSEKFLPNPFLPGERMYRTGDLAKMRPDGHIEYLGRIDHQVKIRGYRIELGEIEHQLLRHSDIKEAAVAAKTD
QNNDQVLCAYVVSERDITQKDIKTFLAKELPEYMVPSYLLKLDELPLTPNGKVDLKALPEPDRSAGALLEYEPPRHELEEKMAAIWEDILNIEQIGINANIFDIGANSLNVMSFVSR
LYAELGFRVPFKDIFSKPTIKELSDFLKHAQDLLKDYTDDCMQLTRAEEGGKNLFCFPPAASMGIAYMGLAKHLKQHSVYSFNFIPSANRIRKYADIIKNIQGEGPYTLIGYSSGGI
LAFDVAKELNRQGYEVEDLIIIDSKYRTKAEKHQFTEEEYREEISKTFELEKYRDVEKLLSDYLVDLVMKSYVYIQNTVTTGAIDGHISYIKSSDNQRGENMMMWEKATSKTFTVV
QGAGTHMQMISKSHPDILERNARLIHDIINKTVKIGSRSHHHHHH

### 15) BacC4-C5[A-PCP-E-COM(Ins5086GGS)-C-A-PCP-TE]-$H_6$

MDSDEEMNALLDQNQGKQADYPQDQTVHQLFEQQADKTPEQTAVVYADEKLTYRELNERANQLARLLRDKGADADQPVAIMIEPSLEMIISMLAVLKAGAAYVPIEPEQLAKR
TNEILSDSRAAILLVKGSVKENVAFAGEIVNVADGLIDAKVASNLSASGSADQNAYIIYTSGSTGKPKGVFVRHGNVVNYTTWFMKEAGLTENDKAMLVSSYAFDLGYTSIFSAL
LSGSELHIARKECYTNAHRALKYIKENGITYIKLTPSLFNIFVNDPGFSAEKPCATLRLVVLGGEMINTRDVETFYNQYPDHVVMNHYGPTETTIGSVFKVIDPEHLDSFKECPVIG
TPIHNTNAYVLDENMKLLPEGVYGELCIAGAGVTGGYVNRPDETKEKFIENPFAPHTKMYRTGDLARRLSDGNIELAGRIDTQVKVRGYRIEPEEIKNRLLAHDDIKEAFIAARED
HKGAKQLCAYFTADAELPFEDIRTYLMHELPEYMIPSSFVQIEKMPLSANGKIDTAALPEPQPGKETEYEPPRNETEEKLVQIWEEVLGIDKIGITHHFFAAGGDSIKALQMISRLS
REGLSLEMKDLFANPQIKSLSRYVKAESDKSASYETVEGEVLLTPIQQEYFSLNKTDRNHYNHAVMLYRKNGFDESIVKRVFKEIIKHHDALRTVFTEEDGKIIQYNRGPDKQLF
DLFVYDVSSENDQPQKVYQLATELQQSIDIETGPLVKLALFKTNNGDHLLIIIHHLVVDGISWRILFEDLAIGYSQLANGEKVEFYPKTASYQAYARHIAEYAKSVKLLSEKQYWLK
AIAEGVEFLDMNENAGAFKVEDSRTFSTELEKEETKRLLRETNRAYHTEINDILITALLVAARDMNGQNQLRITLEGHGREQVADGIDISRTVGWFTSKYPVFIDLGQETDMSRTI
KMVKEHLRNVPNKGIGYGILKYLTRDSEIAKGAASPILFNYLGQLDEDINSGEFSSSHLSPGEAAGKGITREHPLEINAVVFRGKLAIQTTYNTRAYSEDVVRAFAQNYKEALKAV
IRHCAEREETEKTPSDYGDKGISLDQLEEIKLKYKGggsMEIEKIYPLANMQRGMLFHALEDKESQAYFEQMAINMKGLIDERLFAETFNDIMERHEILRASIEYEITDEPRNVIIKD
RKINLDYHDLRKQSPAEREQVIQAYRKADREKGFRLNSEPLIRAALMRTEDDSYTFIWTNHHILLDGWSRGIIMGELFHMYHMKEARQKHRLEEARPYSDYIGWLQQQDKEAA
KAYWRNYLSGFTEKSPISVLAGSSGHAKYKRKEAVIEFPEQLTGRITELASRNNVTFHTVLQCIWGMLLARYNQTDDVVFGTVISGRDAQVTGIEKMVGLFINTVPTRIRLDKSQ
SFKELIKSVQEQALEGRTYHDMNLSEVQSLSELKRELLDHILIFENYAVDQSAFETSGKRGAGFVFEEIHAEEQTNYGFNIVAVPGERLVIKLTYDGNIYHDHIIAGIKGHLQQVME
QVVQHEDQSLNDITVLSEAERNRLLYEWNDTKAEYPNQTIHRLFEEQAEKTPELAAVVSGNDKLTYRELNEKSNQLARYLRDKGVKADTIVAIMAERSPEMVVGIMGILKAGGA
YLPIDPDYPEERIKYMLEDSGAAIILADHKQDLGTLHQEAVELTGDFSSYPADNLEPAGNADSLAYIIYTSGSTGKPKGVMIRQRGLVNYITWADRVYVQGEQLDFALYSSIAFDL
TVTSIFTPLISGNRVIVYRHSEDGEPLIRKVFRDQKAGIVKLTPSHLSLVKDMDASGSSIKRLIVGGEDLKTELAKEITERFHHNIEIYNEYGPTETVVGCMIYQYDAGWDRQVSVP
IGKPASNVQLYILDERQEVQPVGIAGELYISGDGVAKGYLNKPELTSEKFLPNPFLPGERMYRTGDLAKMRPDGHIEYLGRIDHQVKIRGYRIELGEIEHQLLRHSDIKEAAVAAK
TDQNNDQVLCAYVVSERDITQKDIKTFLAKELPEYMVPSYLLKLDELPLTPNGKVDLKALPEPDRSAGALLEYEPPRHELEEKMAAIWEDILNIEQIGINANIFDIGANSLNVMSFV
SRLYAELGFRVPFKDIFSKPTIKELSDFLKHAQDLLKDYTDDCMQLTRAEEGGKNLFCFPPAASMGIAYMGLAKHLKQHSVYSFNFIPSANRIRKYADIIKNIQGEGPYTLIGYSS
GGILAFDVAKELNRQGYEVEDLIIIDSKYRTKAEKHQFTEEEYREEISKTFELEKYRDVEKLLSDYLVDLVMKSYVYIQNTVTTGAIDGHISYIKSSDNQRGENMMMWEKATSKTF
TVVQGAGTHMQMISKSHPDILERNARLIHDIINKTVKIGSRS**HHHHHH**

**16) SBP-TycA[A-PCP-E-COM$^{D}$]**

**MDEKTTGWRGGHVVEGLAGELEQLRARLEHHPQGQREP**MVANQANLIDNKRELEQHALVPYAQGKSIHQLFEEQAEAFPDRVAIVFENRRLSYQELNRKANQLARALLEKG
VQTDSIVGVMMEKSIENVIAILAVLKAGGAYVPIDIEYPRDRIQYILQDSQTKIVLTQKSVSQLVHDVGYSGEVVVLDEEQLDARETANLHQPSKPTDLAYVIYTSGTTGKPKGTM
LEHKGIANLQSFFQNSFGVTEQDRIGLFASMSFDASVWEMFMALLSGASLYILSKQTIHDFAAFEHYLSENELTIITLPPTYLTHLTPERITSLRIMITAGSASSAPLVNKWKDKLR
YINAYGPTETSICATIWEAPSNQLSVQSVPIGKPIQNTHIYIVNEDLQLLPTGSEGELCIGGVGLARGYWNRPDLTAEKFVDNPFVPGEKMYRTGDLAKWLTDGTIEFLGRIDHQ
VKIRGHRIELGEIESVLLAHEHITEAVVIAREDQHAGQYLCAYYISQQEATPAQLRDYAAQKLPAYMLPSYFVKLDKMPLTPNDKIDRKALPEPDLTANQSQAAYHPPRTETESIL
VSIWQNVLGIEKIGIRDNFYSLGGDSIQAIQVVARLHSYQLKLETKDLLNYPTIEQVALFVKSTTRKSDQGIIAGNVPLTPIQKWFFGKNFTNTGHWNQSSVLYRPEGFDPKVIQS
VMDKIIEHHDALRMVYQHENGNVVQHNRGLGGQLYDFFSYNLTAQPDVQQAIEAETQRLHSSMNLQEGPLVKVALFQTLHGDHLFLAIHHLVVDGISWRILFEDLATGYAQAL
AGQAISLPEKTDSFQSWSQWLQEYANEADLLSEIPYWESLESQAKNVSLPKDYEVTDCKQKSVRNMRIRLHPEETEQLLKHANQAYQTEINDLLLAALGLAFAEWSKLAQIVIH
LEGHGREDIIEQANVARTVGWFTSQYPVLLDLKQTAPLSDYIKLTKENMRKIPRKGIGYDILKHVTLPENRGSLSFRVQPEVTFNYLGQFDADMRTELFTRSPYSGGNTLGADG
KNNLSPESEVYTALNITGLIEGGELVLTFSYSSEQYREESIQQLSQSYQKHLLAIIAHCTEKKEVERTPSDFSVKGLQMEEMDDIFELLANTLR

**17) TycB1[COM$^{A}$-A-PCP]-His$_{6}$**

MSVFSKEQVQDMYALTPMQEGMLFHALLDQEHNSHLVQMSISLQGDLDVGLFTDSLHVLVERYDVFRTLFLYEKLKQPLQVVLKQRPIPIEFYDLSACDESEKQLRYTQYKRA
DQERTFHLAKDPLMRVALFQMSQHDYQVIWSFHHILMDGWCFSIIFDDLLAIYLSLQNKTALSLEPVQPYSRFINWLEKQNKQAALNYWSDYLEAYEQKTTLPKKEAAFAKAFQ
PTQYRFSLNRTLTKQLGTIASQNQVTLSTVIQTIWGVLLQKYNAAHDVLFGSIVSGRPTDIVGIDKMVGLFINTIPFRVQAKAGQTFSELLQAVHKRTLQSQPYEHVPLYDIQTQS
VLKQELIDHLLVIENYPLVEALQKKALNQQIGFTITAVEMFEPTNYDLTVMVMPKEELAFRFDYNAALFDEQVVQKLAGHLQQIADCVANNSGVELCQIPLLTEAETSQLLAKRTE
TAADYPAATMHELFSRQAEKTPEQVAVVFADQHLTYRELDEKSNQLARFLRKKGIGTGSLVGTLLDRSLDMIVGILGVLKAGGAFVPIDPELPAERIAYMLTHSRVPLVVTQNHL
RAKVTTPTETIDINTAVIGEESRAPIESLNQPHDLFYIIYTSGTTGQPKGVMLEHRNMANLMHFTFDQTNIAFHEKVLQYTTCSFDVCYQEIFSTLLSGGQLYLITNELRRHVEKLF
AFIQEKQISILSLPVSFLKFIFNEQDYAQSFPRCVKHIITAGEQLVVTHELQKYLRQHRVFLHNHYGPSETHVVTTCTMDPGQAIPELPPIGKPISNTGIYILDEGLQLKPEGIVGEL
YISGANVGRGYLHQPELTAEKFLDNPYQPGERMYRTGDLARWLPDGQLEFLGRIDHQVKIRGHRIELGEIESRLLNHPAIKEAVVIDRADETGGKFLCAYVVLQKALSDEEMRA
YLAQALPEYMIPSFFVTLERIPVTPNGKTDRRALPKPEGSAKTKADYVAPTTELEQKLVAIWEQILGVSPIGIQDHFFTLGGHSLKAIQLISRIQKECQADVPLRVLFEQPTIQALAA
YVEGSRS**HHHHHH**

**18) SBP-TycA[A-PCP-E-COM$^{D}$(ΔHlx)]**

**MDEKTTGWRGGHVVEGLAGELEQLRARLEHHPQGQREP**MVANQANLIDNKRELEQHALVPYAQGKSIHQLFEEQAEAFPDRVAIVFENRRLSYQELNRKANQLARALLEKG
VQTDSIVGVMMEKSIENVIAILAVLKAGGAYVPIDIEYPRDRIQYILQDSQTKIVLTQKSVSQLVHDVGYSGEVVVLDEEQLDARETANLHQPSKPTDLAYVIYTSGTTGKPKGTM
LEHKGIANLQSFFQNSFGVTEQDRIGLFASMSFDASVWEMFMALLSGASLYILSKQTIHDFAAFEHYLSENELTIITLPPTYLTHLTPERITSLRIMITAGSASSAPLVNKWKDKLR
YINAYGPTETSICATIWEAPSNQLSVQSVPIGKPIQNTHIYIVNEDLQLLPTGSEGELCIGGVGLARGYWNRPDLTAEKFVDNPFVPGEKMYRTGDLAKWLTDGTIEFLGRIDHQ
VKIRGHRIELGEIESVLLAHEHITEAVVIAREDQHAGQYLCAYYISQQEATPAQLRDYAAQKLPAYMLPSYFVKLDKMPLTPNDKIDRKALPEPDLTANQSQAAYHPPRTETESIL
VSIWQNVLGIEKIGIRDNFYSLGGDSIQAIQVVARLHSYQLKLETKDLLNYPTIEQVALFVKSTTRKSDQGIIAGNVPLTPIQKWFFGKNFTNTGHWNQSSVLYRPEGFDPKVIQS
VMDKIIEHHDALRMVYQHENGNVVQHNRGLGGQLYDFFSYNLTAQPDVQQAIEAETQRLHSSMNLQEGPLVKVALFQTLHGDHLFLAIHHLVVDGISWRILFEDLATGYAQAL
AGQAISLPEKTDSFQSWSQWLQEYANEADLLSEIPYWESLESQAKNVSLPKDYEVTDCKQKSVRNMRIRLHPEETEQLLKHANQAYQTEINDLLLAALGLAFAEWSKLAQIVIH
LEGHGREDIIEQANVARTVGWFTSQYPVLLDLKQTAPLSDYIKLTKENMRKIPRKGIGYDILKHVTLPENRGSLSFRVQPEVTFNYLGQFDADMRTELFTRSPYSGGNTLGADG
KNNLSPESEVYTALNITGLIEGGELVLTFSYSSEQYREESIQQLSQSYQKHLLAIIAHCTEKKEVERTPSDFSVKGLQ

**19) SBP-TycA[A-PCP-E-COM$^{D}$(ΔLinker)]**

**MDEKTTGWRGGHVVEGLAGELEQLRARLEHHPQGQREP**MVANQANLIDNKRELEQHALVPYAQGKSIHQLFEEQAEAFPDRVAIVFENRRLSYQELNRKANQLARALLEKG
VQTDSIVGVMMEKSIENVIAILAVLKAGGAYVPIDIEYPRDRIQYILQDSQTKIVLTQKSVSQLVHDVGYSGEVVVLDEEQLDARETANLHQPSKPTDLAYVIYTSGTTGKPKGTM
LEHKGIANLQSFFQNSFGVTEQDRIGLFASMSFDASVWEMFMALLSGASLYILSKQTIHDFAAFEHYLSENELTIITLPPTYLTHLTPERITSLRIMITAGSASSAPLVNKWKDKLR
YINAYGPTETSICATIWEAPSNQLSVQSVPIGKPIQNTHIYIVNEDLQLLPTGSEGELCIGGVGLARGYWNRPDLTAEKFVDNPFVPGEKMYRTGDLAKWLTDGTIEFLGRIDHQ
VKIRGHRIELGEIESVLLAHEHITEAVVIAREDQHAGQYLCAYYISQQEATPAQLRDYAAQKLPAYMLPSYFVKLDKMPLTPNDKIDRKALPEPDLTANQSQAAYHPPRTETESIL
VSIWQNVLGIEKIGIRDNFYSLGGDSIQAIQVVARLHSYQLKLETKDLLNYPTIEQVALFVKSTTRKSDQGIIAGNVPLTPIQKWFFGKNFTNTGHWNQSSVLYRPEGFDPKVIQS
VMDKIIEHHDALRMVYQHENGNVVQHNRGLGGQLYDFFSYNLTAQPDVQQAIEAETQRLHSSMNLQEGPLVKVALFQTLHGDHLFLAIHHLVVDGISWRILFEDLATGYAQAL
AGQAISLPEKTDSFQSWSQWLQEYANEADLLSEIPYWESLESQAKNVSLPKDYEVTDCKQKSVRNMRIRLHPEETEQLLKHANQAYQTEINDLLLAALGLAFAEWSKLAQIVIH
LEGHGREDIIEQANVARTVGWFTSQYPVLLDLKQTAPLSDYIKLTKENMRKIPRKGIGYDILKHVTLPENRGSLSFRVQPEVTFNYLGQFDADMRTELFTRSPYSGGNTLGADG
KNNLSPESEVYTALNITGLIEGGELVLTFSYSSEQYREESIQQLSQSYQKHLLAIIAHCTEKKEGLQMEEMDDIFELLANTLR

**20) SBP-TycA[A-PCP-E-COM$^{D}$(VER(GGS)$_{3}$TPSD)]**

**MDEKTTGWRGGHVVEGLAGELEQLRARLEHHPQGQREP**MVANQANLIDNKRELEQHALVPYAQGKSIHQLFEEQAEAFPDRVAIVFENRRLSYQELNRKANQLARALLEKG
VQTDSIVGVMMEKSIENVIAILAVLKAGGAYVPIDIEYPRDRIQYILQDSQTKIVLTQKSVSQLVHDVGYSGEVVVLDEEQLDARETANLHQPSKPTDLAYVIYTSGTTGKPKGTM
LEHKGIANLQSFFQNSFGVTEQDRIGLFASMSFDASVWEMFMALLSGASLYILSKQTIHDFAAFEHYLSENELTIITLPPTYLTHLTPERITSLRIMITAGSASSAPLVNKWKDKLR
YINAYGPTETSICATIWEAPSNQLSVQSVPIGKPIQNTHIYIVNEDLQLLPTGSEGELCIGGVGLARGYWNRPDLTAEKFVDNPFVPGEKMYRTGDLAKWLTDGTIEFLGRIDHQ
VKIRGHRIELGEIESVLLAHEHITEAVVIAREDQHAGQYLCAYYISQQEATPAQLRDYAAQKLPAYMLPSYFVKLDKMPLTPNDKIDRKALPEPDLTANQSQAAYHPPRTETESIL
VSIWQNVLGIEKIGIRDNFYSLGGDSIQAIQVVARLHSYQLKLETKDLLNYPTIEQVALFVKSTTRKSDQGIIAGNVPLTPIQKWFFGKNFTNTGHWNQSSVLYRPEGFDPKVIQS
VMDKIIEHHDALRMVYQHENGNVVQHNRGLGGQLYDFFSYNLTAQPDVQQAIEAETQRLHSSMNLQEGPLVKVALFQTLHGDHLFLAIHHLVVDGISWRILFEDLATGYAQAL
AGQAISLPEKTDSFQSWSQWLQEYANEADLLSEIPYWESLESQAKNVSLPKDYEVTDCKQKSVRNMRIRLHPEETEQLLKHANQAYQTEINDLLLAALGLAFAEWSKLAQIVIH
LEGHGREDIIEQANVARTVGWFTSQYPVLLDLKQTAPLSDYIKLTKENMRKIPRKGIGYDILKHVTLPENRGSLSFRVQPEVTFNYLGQFDADMRTELFTRSPYSGGNTLGADG
KNNLSPESEVYTALNITGLIEGGELVLTFSYSSEQYREESIQQLSQSYQKHLLAIIAHCTEKKEVERggsggsggsTPSDFSVKGLQMEEMDDIFELLANTLR

**21) SBP-TycA[A-PCP-E-COM$^{D}$(ΔLinker_sm)]**

**MDEKTTGWRGGHVVEGLAGELEQLRARLEHHPQGQREP**MVANQANLIDNKRELEQHALVPYAQGKSIHQLFEEQAEAFPDRVAIVFENRRLSYQELNRKANQLARALLEKG
VQTDSIVGVMMEKSIENVIAILAVLKAGGAYVPIDIEYPRDRIQYILQDSQTKIVLTQKSVSQLVHDVGYSGEVVVLDEEQLDARETANLHQPSKPTDLAYVIYTSGTTGKPKGTM
LEHKGIANLQSFFQNSFGVTEQDRIGLFASMSFDASVWEMFMALLSGASLYILSKQTIHDFAAFEHYLSENELTIITLPPTYLTHLTPERITSLRIMITAGSASSAPLVNKWKDKLR
YINAYGPTETSICATIWEAPSNQLSVQSVPIGKPIQNTHIYIVNEDLQLLPTGSEGELCIGGVGLARGYWNRPDLTAEKFVDNPFVPGEKMYRTGDLAKWLTDGTIEFLGRIDHQ
VKIRGHRIELGEIESVLLAHEHITEAVVIAREDQHAGQYLCAYYISQQEATPAQLRDYAAQKLPAYMLPSYFVKLDKMPLTPNDKIDRKALPEPDLTANQSQAAYHPPRTETESIL
VSIWQNVLGIEKIGIRDNFYSLGGDSIQAIQVVARLHSYQLKLETKDLLNYPTIEQVALFVKSTTRKSDQGIIAGNVPLTPIQKWFFGKNFTNTGHWNQSSVLYRPEGFDPKVIQS
VMDKIIEHHDALRMVYQHENGNVVQHNRGLGGQLYDFFSYNLTAQPDVQQAIEAETQRLHSSMNLQEGPLVKVALFQTLHGDHLFLAIHHLVVDGISWRILFEDLATGYAQAL
AGQAISLPEKTDSFQSWSQWLQEYANEADLLSEIPYWESLESQAKNVSLPKDYEVTDCKQKSVRNMRIRLHPEETEQLLKHANQAYQTEINDLLLAALGLAFAEWSKLAQIVIH
LEGHGREDIIEQANVARTVGWFTSQYPVLLDLKQTAPLSDYIKLTKENMRKIPRKGIGYDILKHVTLPENRGSLSFRVQPEVTFNYLGQFDADMRTELFTRSPYSGGNTLGADG
KNNLSPESEVYTALNITGLIEGGELVLTFSYSSEQYREESIQQLSQSYQKHLLAIIAHCTEKKETPSDFSVKGLQMEEMDDIFELLANTLR

**22) SBP-TycA[A-PCP-E-COM$^{D}$(VERgTPSD)]**

**MDEKTTGWRGGHVVEGLAGELEQLRARLEHHPQGQREP**MVANQANLIDNKRELEQHALVPYAQGKSIHQLFEEQAEAFPDRVAIVFENRRLSYQELNRKANQLARALLEKG
VQTDSIVGVMMEKSIENVIAILAVLKAGGAYVPIDIEYPRDRIQYILQDSQTKIVLTQKSVSQLVHDVGYSGEVVVLDEEQLDARETANLHQPSKPTDLAYVIYTSGTTGKPKGTM
LEHKGIANLQSFFQNSFGVTEQDRIGLFASMSFDASVWEMFMALLSGASLYILSKQTIHDFAAFEHYLSENELTIITLPPTYLTHLTPERITSLRIMITAGSASSAPLVNKWKDKLR
YINAYGPTETSICATIWEAPSNQLSVQSVPIGKPIQNTHIYIVNEDLQLLPTGSEGELCIGGVGLARGYWNRPDLTAEKFVDNPFVPGEKMYRTGDLAKWLTDGTIEFLGRIDHQ
VKIRGHRIELGEIESVLLAHEHITEAVVIAREDQHAGQYLCAYYISQQEATPAQLRDYAAQKLPAYMLPSYFVKLDKMPLTPNDKIDRKALPEPDLTANQSQAAYHPPRTETESIL
VSIWQNVLGIEKIGIRDNFYSLGGDSIQAIQVVARLHSYQLKLETKDLLNYPTIEQVALFVKSTTRKSDQGIIAGNVPLTPIQKWFFGKNFTNTGHWNQSSVLYRPEGFDPKVIQS
VMDKIIEHHDALRMVYQHENGNVVQHNRGLGGQLYDFFSYNLTAQPDVQQAIEAETQRLHSSMNLQEGPLVKVALFQTLHGDHLFLAIHHLVVDGISWRILFEDLATGYAQAL
AGQAISLPEKTDSFQSWSQWLQEYANEADLLSEIPYWESLESQAKNVSLPKDYEVTDCKQKSVRNMRIRLHPEETEQLLKHANQAYQTEINDLLLAALGLAFAEWSKLAQIVIH
LEGHGREDIIEQANVARTVGWFTSQYPVLLDLKQTAPLSDYIKLTKENMRKIPRKGIGYDILKHVTLPENRGSLSFRVQPEVTFNYLGQFDADMRTELFTRSPYSGGNTLGADG
KNNLSPESEVYTALNITGLIEGGELVLTFSYSSEQYREESIQQLSQSYQKHLLAIIAHCTEKKEVERgTPSDFSVKGLQMEEMDDIFELLANTLR

**23) SBP-TycA[A-PCP-E-COM$^{D}$(ΔV1059)]**

**MDEKTTGWRGGHVVEGLAGELEQLRARLEHHPQGQREP**MVANQANLIDNKRELEQHALVPYAQGKSIHQLFEEQAEAFPDRVAIVFENRRLSYQELNRKANQLARALLEKG
VQTDSIVGVMMEKSIENVIAILAVLKAGGAYVPIDIEYPRDRIQYILQDSQTKIVLTQKSVSQLVHDVGYSGEVVVLDEEQLDARETANLHQPSKPTDLAYVIYTSGTTGKPKGTM
LEHKGIANLQSFFQNSFGVTEQDRIGLFASMSFDASVWEMFMALLSGASLYILSKQTIHDFAAFEHYLSENELTIITLPPTYLTHLTPERITSLRIMITAGSASSAPLVNKWKDKLR
YINAYGPTETSICATIWEAPSNQLSVQSVPIGKPIQNTHIYIVNEDLQLLPTGSEGELCIGGVGLARGYWNRPDLTAEKFVDNPFVPGEKMYRTGDLAKWLTDGTIEFLGRIDHQ
VKIRGHRIELGEIESVLLAHEHITEAVVIAREDQHAGQYLCAYYISQQEATPAQLRDYAAQKLPAYMLPSYFVKLDKMPLTPNDKIDRKALPEPDLTANQSQAAYHPPRTETESIL
VSIWQNVLGIEKIGIRDNFYSLGGDSIQAIQVVARLHSYQLKLETKDLLNYPTIEQVALFVKSTTRKSDQGIIAGNVPLTPIQKWFFGKNFTNTGHWNQSSVLYRPEGFDPKVIQS
VMDKIIEHHDALRMVYQHENGNVVQHNRGLGGQLYDFFSYNLTAQPDVQQAIEAETQRLHSSMNLQEGPLVKVALFQTLHGDHLFLAIHHLVVDGISWRILFEDLATGYAQAL
AGQAISLPEKTDSFQSWSQWLQEYANEADLLSEIPYWESLESQAKNVSLPKDYEVTDCKQKSVRNMRIRLHPEETEQLLKHANQAYQTEINDLLLAALGLAFAEWSKLAQIVIH
LEGHGREDIIEQANVARTVGWFTSQYPVLLDLKQTAPLSDYIKLTKENMRKIPRKGIGYDILKHVTLPENRGSLSFRVQPEVTFNYLGQFDADMRTELFTRSPYSGGNTLGADG
KNNLSPESEVYTALNITGLIEGGELVLTFSYSSEQYREESIQQLSQSYQKHLLAIIAHCTEKKEERTPSDFSVKGLQMEEMDDIFELLANTLR

### 24) SBP-TycA[A-PCP-E-COM$^{D}$(P1063A)]

**MDEKTTGWRGGHVVEGLAGELEQLRARLEHHPQGQREP**MVANQANLIDNKRELEQHALVPYAQGKSIHQLFEEQAEAFPDRVAIVFENRRLSYQELNRKANQLARALLEKG
VQTDSIVGVMMEKSIENVIAILAVLKAGGAYVPIDIEYPRDRIQYILQDSQTKIVLTQKSVSQLVHDVGYSGEVVVLDEEQLDARETANLHQPSKPTDLAYVIYTSGTTGKPKGTM
LEHKGIANLQSFFQNSFGVTEQDRIGLFASMSFDASVWEMFMALLSGASLYILSKQTIHDFAAFEHYLSENELTIITLPPTYLTHLTPERITSLRIMITAGSASSAPLVNKWKDKLR
YINAYGPTETSICATIWEAPSNQLSVQSVPIGKPIQNTHIYIVNEDLQLLPTGSEGELCIGGVGLARGYWNRPDLTAEKFVDNPFVPGEKMYRTGDLAKWLTDGTIEFLGRIDHQ
VKIRGHRIELGEIESVLLAHEHITEAVVIAREDQHAGQYLCAYYISQQEATPAQLRDYAAQKLPAYMLPSYFVKLDKMPLTPNDKIDRKALPEPDLTANQSQAAYHPPRTETESIL
VSIWQNVLGIEKIGIRDNFYSLGGDSIQAIQVVARLHSYQLKLETKDLLNYPTIEQVALFVKSTTRKSDQGIIAGNVPLTPIQKWFFGKNFTNTGHWNQSSVLYRPEGFDPKVIQS
VMDKIIEHHDALRMVYQHENGNVVQHNRGLGGQLYDFFSYNLTAQPDVQQAIEAETQRLHSSMNLQEGPLVKVALFQTLHGDHLFLAIHHLVVDGISWRILFEDLATGYAQAL
AGQAISLPEKTDSFQSWSQWLQEYANEADLLSEIPYWESLESQAKNVSLPKDYEVTDCKQKSVRNMRIRLHPEETEQLLKHANQAYQTEINDLLLAALGLAFAEWSKLAQIVIH
LEGHGREDIIEQANVARTVGWFTSQYPVLLDLKQTAPLSDYIKLTKENMRKIPRKGIGYDILKHVTLPENRGSLSFRVQPEVTFNYLGQFDADMRTELFTRSPYSGGNTLGADG
KNNLSPESEVYTALNITGLIEGGELVLTFSYSSEQYREESIQQLSQSYQKHLLAIIAHCTEKKEVERTASDFSVKGLQMEEMDDIFELLANTLR

### 25) TycB1[COM$^{A}$(S5Bpa)-A-PCP]

SVFBpaKEQVQDMYALTPMQEGMLFHALLDQEHNSHLVQMSISLQGDLDVGLFTDSLHVLVERYDVFRTLFLYEKLKQPLQVVLKQRPIPIEFYDLSACDESEKQLRYTQYKRA
DQERTFHLAKDPLMRVALFQMSQHDYQVIWSFHHILMDGWCFSIIFDDLLAIYLSLQNKTALSLEPVQPYSRFINWLEKQNKQAALNYWSDYLEAYEQKTTLPKKEAAFAKAFQ
PTQYRFSLNRTLTKQLGTIASQNQVTLSTVIQTIWGVLLQKYNAAHDVLFGSIVSGRPTDIVGIDKMVGLFINTIPFRVQAKAGQTFSELLQAVHKRTLQSQPYEHVPLYDIQTQS
VLKQELIDHLLVIENYPLVEALQKKALNQQIGFTITAVEMFEPTNYDLTVMVMPKEELAFRFDYNAALFDEQVVQKLAGHLQQIADCVANNSGVELCQIPLLTEAETSQLLAKRTE
TAADYPAATMHELFSRQAEKTPEQVAVVFADQHLTYRELDEKSNQLARFLRKKGIGTGSLVGTLLDRSLDMIVGILGVLKAGGAFVPIDPELPAERIAYMLTHSRVPLVVTQNHL
RAKVTTPTETIDINTAVIGEESRAPIESLNQPHDLFYIIYTSGTTGQPKGVMLEHRNMANLMHFTFDQTNIAFHEKVLQYTTCSFDVCYQEIFSTLLSGGQLYLITNELRRHVEKLF
AFIQEKQISILSLPVSFLKFIFNEQDYAQSFPRCVKHIITAGEQLVVTHELQKYLRQHRVFLHNHYGPSETHVVTTCTMDPGQAIPELPPIGKPISNTGIYILDEGLQLKPEGIVGEL
YISGANVGRGYLHQPELTAEKFLDNPYQPGERMYRTGDLARWLPDGQLEFLGRIDHQVKIRGHRIELGEIESRLLNHPAIKEAVVIDRADETGGKFLCAYVVLQKALSDEEMRA
YLAQALPEYMIPSFFVTLERIPVTPNGKTDRRALPKPEGSAKTKADYVAPTTELEQKLVAIWEQILGVSPIGIQDHFFTLGGHSLKAIQLISRIQKECQADVPLRVLFEQPTIQALAA
YVEGSRSHHHHHH

### Supporting References

[1] G. W. Heberlig, J. J. La Clair, M. D. Burkart, Nature, 2025, 638, 261-269.

[2] J. Rüschenbaum, W. Steinchen, F. Mayerthaler, A. L. Feldberg, H. D. Mootz, Angew Chem Int Ed Engl, 2022, 61, e202212994.
